# Protein multiple conformations prediction using multi-objective evolution algorithm

**DOI:** 10.1101/2023.04.21.537776

**Authors:** Ming-Hua Hou, Si-Rong Jin, Xin-Yue Cui, Chun-Xiang Peng, Kai-Long Zhao, Le Song, Gui-Jun Zhang

## Abstract

**Motivation:** With the breakthrough of AlphaFold2 and the publication of AlphaFold DB, the protein structure prediction has made remarkable progress, which may further promote many potential applications of proteomics in all areas of life. However, it should be noted that AlphaFold2 models tend to represent only a single static structure, and accurately predicting multiple conformations remains a challenge. Therefore, it is essential to develop methods for predicting multiple conformations, which enable us to gain knowledge of multiple conformational states and the broader conformational landscape to better understand the mechanism of action.

**Results:** In this work, we proposed a multiple conformational states folding method using the distance-based multi-objective evolutionary algorithm framework, named MultiSFold. First, a multi-objective energy landscape with multiple competing constraints generated by deep learning is constructed. Then, an iterative modal exploration and exploitation strategy based on multi-objective optimization, geometric optimization and structural similarity clustering is designed to perform conformational sampling. Finally, the final population is generated using a loop-specific perturbation strategy to adjust the spatial orientations. MultiSFold was compared with state-of-the-art methods on a developed benchmark testset containing 81 proteins with two representative conformational states. Based on the proposed metric, the success ratio of MultiSFold predicting multiple conformations was 70.4% while that of AlphaFold2 was 9.88%, which may indicate that conformational sampling combined with knowledge gained through deep learning has the potential to produce conformations spanned the range between two experimental structures. In addition, MultiSFold was tested on 244 human proteins with low structural accuracy in AlphaFold DB to test whether it could further improve the accuracy of static structures. The experimental results demonstrate that the TM-score of MultiSFold is 2.97% and 7.72% higher than that of AlphaFold2 and RoseTTAFold, respectively, supporting our hypothesis that multiple competing optimization objectives can further assist conformational search to improve prediction accuracy.

## 1 Introduction

The AlphaFold2 (Jumper *et al*, 2021) from DeepMind has achieved an astonishing accuracy in predicting the three-dimensional coordinates of a protein, which represents a dramatic advance in structural biology (Subramaniam and Kleywegt, 2022). Fierce competition has produced a range of comparable approaches to improve the method even further and help illuminate proteomes and their many interactions (Jones and Thornton, 2022). However, this does not mean that all structural prediction problems can be solved using AlphaFold2. For example, where a protein is known to have multiple conformations, AlphaFold2 usually only produces one of them and the output conformation cannot be reliably controlled (Tunyasuvunakool *et al*, 2022). When only static structures are available, the dynamic processes crucial to protein function are difficult to elucidate (Henzler-Wildman *et al*, 2007; Greener *et al*, 2017).

DeepMind collaborated with the European Bioinformatics Institute (EMBL-EBI) of the European Molecular Biology Laboratory to provide structure predictions at the proteomic scale (Tunyasuvunakool *et al*, 2021), resulting in the AlphaFold Protein Structure Database resource (AlphaFold DB; https://alphafold.ebi.ac.uk). The initial release of AlphaFold DB contains over 360,000 predicted structures across 21 model-organism proteomes, and now it provides open access to over 200 million protein structure predictions to accelerate scientific research (Tunyasuvunakool *et al*, 2022). This effort has generated great interest in the life science community as accurate protein prediction has many potential proteomic applications in all areas of life (Jones and Thornton, 2022). However, it has been reported that the accuracy of proteins in the AlphaFold DB is largely affected by whether they have at least part of the sequence can be homology modeled from a structure in the PDB (Thornton *et al*, 2021). When the target proteins with no close homolog in the PDB, the unreliable prediction models generated by AlphaFold2 may be further improved through combining the information captured by competitive methods. For example, another end-to-end prediction method, RoseTTAFold (Baek *et al*, 2021), obtains the prediction model with a three-track network in which information at the one-dimensional sequence level, the two-dimensional distance map level, and the three-dimensional coordinate level is transformed and integrated. In addition, the AlphaFold2 prediction model is a static structure, whereas a protein can exist in one conformation in its "inactive" state and another conformation in its "active" state (Ramanathan *et al*, 2014; Modi and Dunbrack, 2019; Weis and Kobilka, 2018; Xie *et al*, 2020). If an active conformation is predicted by AlphaFold2, then using it to target the inactive state will not work (Skolnick *et al*, 2021).

Dynamic interconversion between multiple conformations underpins the functions of integral membrane proteins in all domains of life (Boehr *et al*, 2009; Shaw *et al*, 2010; Cournia *et al*, 2015; Campbell *et al*, 2016). For example, the vectorial translocation of substrates by transporters is mediated by movements that open and close extra- and intracellular gates (Drew and Boudker, 2016; Kazmier *et al*, 2017). Due to the limitations of experimental techniques such as nuclear magnetic resonance (NMR) and computational methods based on molecular dynamics (MD), various non-MD methods have been used to explore protein dynamics (Greener *et al*, 2017; Zacharias, 2017). CONCOORD (de Groot *et al*, 1997, 1998, 1999) is a distance geometry method to predict the conformational transitions of proteins from an input structure. tCONCOORD (Seeliger *et al*, 2007) extends CONCOORD and gives better sampling of proteins with large conformational changes by predicting hydrogen bonds in the structure that are liable to break. ExProSE (Greener *et al*, 2017) to rapidly explore conformational ensembles using experimental data on two separate conformational states of a protein as input. Deep learning yielded significant improvements in structure prediction and the accuracy of predicted model is close to the experimental static structure (Skolnick *et al*, 2021), which makes it possible to replace the distance information from the experimental structure in the study of conformational transition to a certain extent. Recently, attempts have been made to drive AlphaFold2 to predict diverse conformational ensembles of transporters and receptors by restricting the depth of the input multiple sequence alignments (MSA) (Del Alamo *et al*, 2022). However, the predictive performance is undoubtedly limited by the trained model designed for static structure prediction. In fact, the distance information captured by the model changes due to the adjustment of the input MSA depth, which ultimately results in a different output model. Thereby, it may be a feasible way to predict multiple conformational states using knowledge learned from different predictors rather than from a fixed model by changing the input.

With the arrival of AlphaFold, the toolbox of structural biology has been augmented with a new and formidable set of computational tools, which provides us with an opportunity to gain insight into conformational dynamics (Subramaniam and Kleywegt, 2022). In this work, we proposed a multiple conformational states folding method using the distance-based multi-objective evolutionary algorithm, named MultiSFold. An iterative modal exploration and exploitation strategy combined with a loop-specific dihedral angle optimization strategy to search the conformational space by considering multiple competitive distance-based constrained objectives at the same time. Experimental results on proteins with multiple experimental conformations show that the success ratio of MultiSFold was significantly higher than AlphaFold2, and an ensemble that overlaps both conformations in different states of some proteins was obtained. The ensemble also represents possible intermediates on a transition pathway between two states, which may help to understand the mechanism of conformational transitions and to identify possible sterical barriers (Zacharias, 2017). In addition, MultiSFold tested on 244 human proteins with low-accuracy single static structure of AlphaFold DB, which is support our hypothesis that more optimization objectives are needed in the conformation search to improve the prediction accuracy of a protein structure. Especially, for proteins with pLDDT ≤80 in AlphaFold DB, the quality of these models can be significantly improved by our approach.

## 2 Methods

The pipeline of MultiSFold is illustrated in **Figure 1**. MultiSFold is a multi-objective population optimization method based on multiple distance information, focusing not only on the accuracy of the final structure but also on the generation of multiple models with diversity that may reflect conformational transitions. In addition to the query sequence, the variable-length fragment library generated by our previously proposed method VFlib (Feng *et al*, 2022) and several inter-residue distance maps are required as input. The initial population is generated by randomly fragment assembly. Then, the modal exploration and exploitation strategy is used to iteratively optimize the conformation guided by multiple distance-based objectives in the population. Finally, models generated after a loop-specific dihedral angle optimization were clustered, and the lowest-potential conformation in each cluster is selected as the candidate model. The Framework and specific details of MultiSFold are in **Supplementary Text S1**.

**Fig. 1.**
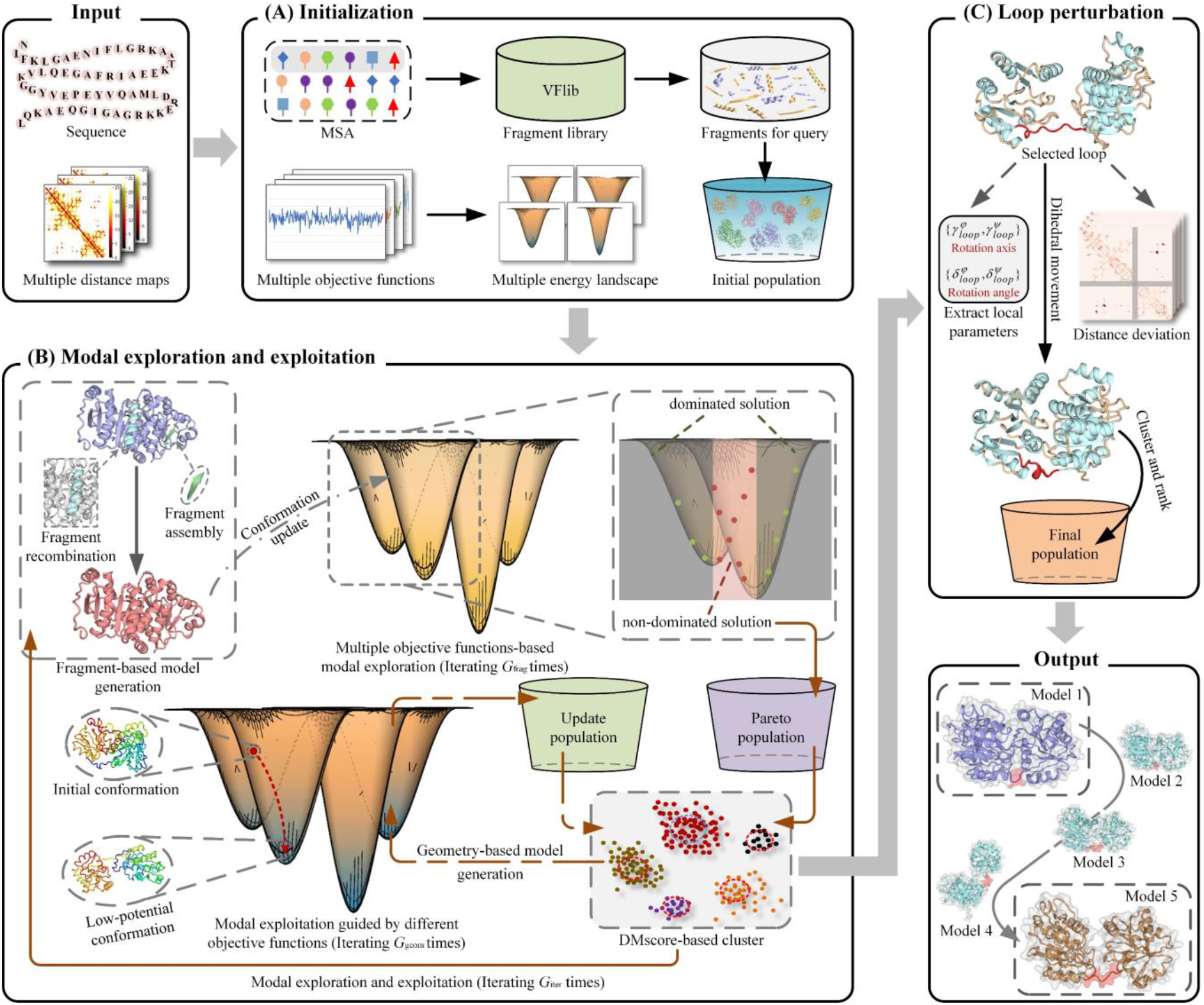
The pipeline of MultiSFold.

### 2.1 Initialization

In this work, the input variable length fragment library will be first generated based on the query sequence and the corresponding MSAs through our previously developed VFlib (Feng *et al*, 2022). Then, several input distance maps generated by different predictors and a consensus distance map based on them will be used to guide the conformation sampling. The predictors used here are two state-of-the-art methods AlphaFold2 and RoseTTAFold as well as our in-house predictor DeepMDisPre, which is an inter-residue multiple distances prediction method based on an improved network which integrates triangle update, axial attention and ResNet to predict multiple distances of residue pairs. The meta-distance map obtained using predicted inter-residue distance may provide more diverse and abundant spatial constraint information for protein folding, which is defined as follows:

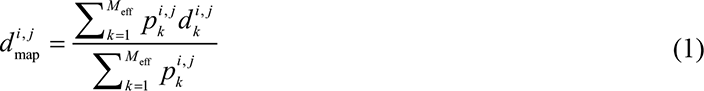

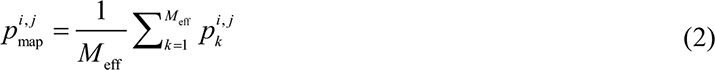

where 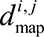 and 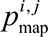 are the distance and confidence between the *i-*th and *j*-th residues of the meta-distance map, respectively; 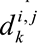 and 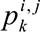 are the distance and confidence between the *i*-th and *j*-th residues in *k*-th input map, respectively; *i* and *j* are the residue indexes; M_eff_ is the number of effective inter-residue distance for the *i*-th and *j*-th residues in all input distance maps.

### 2.2 modal exploration and exploitation

Allostery in proteins is an essential phenomenon in biological processes, and the central physiological function of some proteins is typically achieved by conformational transitions between different states (Turkan et al, 2022). Knowledge of the conformational ensemble span a range of conformations, including the conformations that correspond to different functional states, could give insight into the mechanisms of regulation. Thereby, a modal exploration and exploitation strategy is designed to explore the conformational transitions through a combination of conformational sampling and knowledge gained through deep learning, which is illustrated in **Figure 1B**. In addition, it may be desirable to alleviate the impact of imprecise energy functions through multiple competitive constraints, so as to improve the prediction accuracy of conformation in a certain state.

This strategy first generates decoys through fragment recombination and assembly during the modal exploration stage. A multi-objective optimization process is then used to select decoys for building a Pareto non-dominated solution set, which in turn updates the population. This population is then used to enter the next fragment-based conformation update until the preset number (*G*_frag_) is reached. Once the modal exploration stage is completed, the population has explored some potential basins in the energy landscape. To rapidly update the decoys to reach the nearest energy minimum point, a distance-based geometric optimization method is used. During the modal exploitation stage, each decoy in the population would be selected as the base conformation, and the energy-minimization is applied to generate low-potential models guided by multiple distance constraints. Given iteration cycles (*G*_iter_) were performed for the process of fragment-based modal exploration and geometric-based modal exploitation, with switching between them accomplished through DMscore-based cluster (Zhao et al, 2021) and conformation selection.

#### 2.2.1 multi-objective sampling

In this strategy, multiple conformational states prediction is regarded as a multi-objective optimization problem, which is formulated as follows:

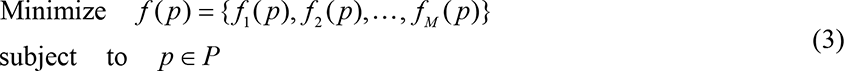

where *f_n_* (*p*), *n* ∈{1, 2, …, *M*} is the *n*-th objective function; *M* represents the total number of objective functions; *p* is the decision vector in the decision space *P*. In this work, *P* ={ *p*_1_, *p*_2_, …,*p_NP_*} represents a set of protein decoys, *NP* is the number of decoys in the population. *f_n_* (*p*) represents the distance potentials 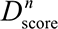 constructed by the inter-residue distance from *n*-th objective, which is defined as follows:

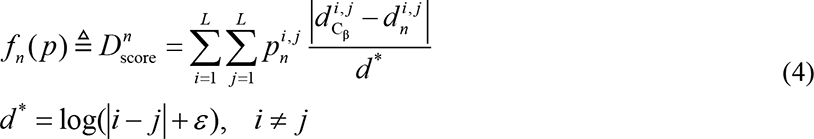

where *L* is the length of the protein sequence; 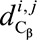 is the calculated distance between Cβ atom (C_α_ for glycine) of the residue pair (*i*, *j*) in the evaluated decoy; 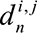 and 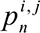 are the distance and confidence between the *i*-th and *j*-th residues in *n*-th distance map, respectively; *n*∈{1, 2,…,*M*} is the index of maps; *ε* is an infinitely small quantity to avoid *d*\* being zero.

In multi-objective optimization, there does not typically exist a feasible solution that minimizes all objective functions simultaneously, and the Pareto dominance is usually used to compare the solutions. A solution *p*_1_ ∈ *P* is said to (Pareto) dominate another solution *p*_2_ ∈ *P*, if

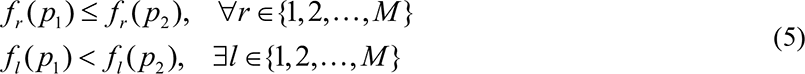

where *r* and *l* represent the index of objective functions.

A solution *p*\*∈ *P* is called a Pareto minimizer or a nondominated solution, if there does not exist another solution that dominates it; that is, there exists no other solution that would decrease some objectives without causing simultaneous increase in at least one other objective. The nondominated decoys (the solution *p* *) will be selected from the population to build a Pareto nondominated solution set.

#### 2.2.2 DMscore-based clustering and conformation selection

Although diversity has been considered as much as possible, the resulting population of each stage may still converge to several local regions and some decoys may be similar. These decoys with similar structures may be further accumulated in the whole algorithm, resulting in bias in the final population, which makes it difficult to predict different conformational states. Thereby, it is necessary to cluster and select decoys before the stage switching. In this strategy, the K-medoids algorithm based on DMscore was used to cluster decoys. DMscore (Zhao et al, 2021) measures the similarity by calculating the difference of Euclidean distance of the corresponding residue pairs in two models, and the clustering algorithm adaptively determine the number of clusters according to the multi-order nearest distance analysis (Hong et al, 2010). DMscore and the specific clustering process are described in the **Supplementary Text S2**. Then, the decoys of each cluster will be sorted by the *E*_total_, which is defined as follows:

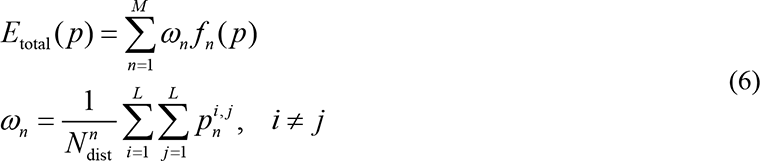

where *M* is the number of objective functions; *L* is the length of the protein sequence; 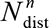 is the number of the effective inter-residue distance of n-th map. Each sorted cluster will be selected in turn and the lowest energy conformation of that cluster will be extracted into the population until a preset number *N*_select_ is reached. In addition, clustering and conformation selection process will also be performed if the number of nondominated solutions exceeds the preset threshold *N*_pareto_.

### 2.3 Loop-specific sampling

The structure of the protein loop region connected with the regular α-helix or β-sheet is flexible and irregular, and its small changes will produce dramatic effect on the entire topology (Liu et al, 2020, 2022). Especially, the domains of multidomain proteins frequently interact with each other to perform higher-level functions, and therefore, multidomain proteins usually have multiple conformational states (Zhang et al, 2022; Peng et al, 2023). According to multiple inter-residue distance, a loop-specific dihedral angle optimization strategy is used to adjust the topology efficiently. For each conformation, the secondary structure is first calculated by DSSP (Kabsch and Sander, 1983), and then the dihedral optimization is performed in turn for the loop regions connected with α-helix or β-sheet at both ends. The aim of this dihedral optimization is to find a set of rotation angles acting on the loop regions, to generate a conformation that satisfies the distance constraint to the greatest extent. In this strategy, multiple constraints are used to guide the dihedral optimization to produce more diverse models in order to span different conformational states. The details of the loop-specific dihedral angle rotation model construction and the dihedral optimization are shown in **Supplementary Text S3**.

## 3 Results and discussion

### 3.1 Benchmark testsets

In order to study the ability of MultiSFold in predicting multiple conformations and modeling conformational transitions, a multi-conformation protein benchmark testset is constructed according to the following criteria: (1) all proteins in the PDB (2022-08) are clustered at 100% sequence identity cut-off; (2) select the clusters in which there is at least one pair of proteins with TM-score ≤ 0.8; (3) the two proteins with the minimum TM-score in each cluster is selected as a representative of that cluster; (4) these clusters are screened out, if there are residue missing at domain boundary for representative structures in these clusters; (5) the final 81 clusters are generated by clustering the clusters generated in previous 4 steps with a 30% sequence identity cut-off. The detailed information of multi-conformation protein targets is shown in **Supplementary Table S2**.

In addition, MultiSFold was tested on a single static structure benchmark testset of 244 human proteins and participated in a blind test of CAMEO to verify our hypothesis that multiple competing optimization objectives may be required in the conformation search to improve the prediction accuracy of a single static structure. The benchmark testset was constructed from AlphaFold Protein Structure Database (AlphaFold DB, 2022-05) as follows. 7,186 out of 23,391 proteins from the Homo sapiens proteome in the AlphaFold DB were firstly picked by culling out proteins that did not have the experimental structure. For the selected proteins with multiple different experimental structures, sequence coverage and structural similarity were used to select one of them as the native structure to evaluate model quality. Then, CD-HIT (Fu et al, 2012) was used to cluster with a 30% sequence identity cut-off, and the representative protein of each cluster constituted 5,158 non-redundant proteins. In order to test the performance of our algorithm on proteins with low-accuracy single static AlphaFold model, proteins were selected if the TM-scores between structures from the AlphaFold DB and PDB were less than 0.9. Finally, 244 non-redundant proteins with a length ranging from 50 to 500 residues were selected as the benchmark testset. The detailed information of benchmark testset and CAMEO blind test targets are shown in **Supplementary Tables S3** and **S4**, respectively. The root mean square deviation (RMSD) and TM-score are used to evaluate the predicted model’s accuracy.

### 3.2 Results of predicting multiple conformations

An important limitation is that the predicted structural models may not provide insights into conformational dynamics. For multidomain and allosteric proteins, for which function is intimately connected to changes in tertiary and quaternary structure spanning multiple conformational states. AlphaFold2 predictions typically generate one of the states of the protein, but understanding the mechanism of action will require knowledge of the broader conformational landscape. In this section, MultiSFold is tested on the multiple conformation protein benchmark testset to investigate whether MultiSFold has the ability to span different conformational states, and compared with state-of-the-art methods AlphaFold2 and RoseTTAFold. The structure models of MultiSFold are from the final population, and the models of MultiSFold^∗^ represent the five lowest-potential decoys from each of the five clusters generated through DMscore-based clustering. The structure models of AlphaFold2 and RoseTTAFold are predicted using its latest standalone package, respectively. The results are summarized in **Supplementary Table S5** and the details for each protein with multiple conformational states are shown in **Supplementary Table S6**.

In this test, models predicted by each method were used to calculate the TM-score to the native structures in different conformational states, and then maximum TM-score is taken. **Figure 2A** shows the resulting TM-score boxplot for 81 proteins in two highly distinct conformational states. For the 81 proteins in conformational state 1, the average TM-score of MultiSFold is 0.774, which is 6.46% and 13.8% higher than that of AlphaFold2 (0.727) and RoseTTAFold (0.680) respectively. The average TM-score of MultiSFold∗ (0.756) is 3.99% and 11.2% higher than that of AlphaFold2 and RoseTTAFold, respectively. In conformational state 2 of the 81 proteins, the average TM-score of MultiSFold is 0.762, which is 4.38% and 14.9% higher than that of AlphaFold2 (0.730) and RoseTTAFold (0.663) respectively. The average TM-score of MultiSFold^∗^ (0.748) is 2.47% and 12.8% higher than that of AlphaFold2 and RoseTTAFold, respectively. It’s worth noting that even if all predictive models capture the same conformational state for a given protein, program statistics may still generate a TM-score for an alternative conformational state that was not identified by the models. Therefore, relying solely on the TM-score and a threshold to determine if multiple conformations have been successfully predicted may not provide an objective assessment.

**Fig. 2.**
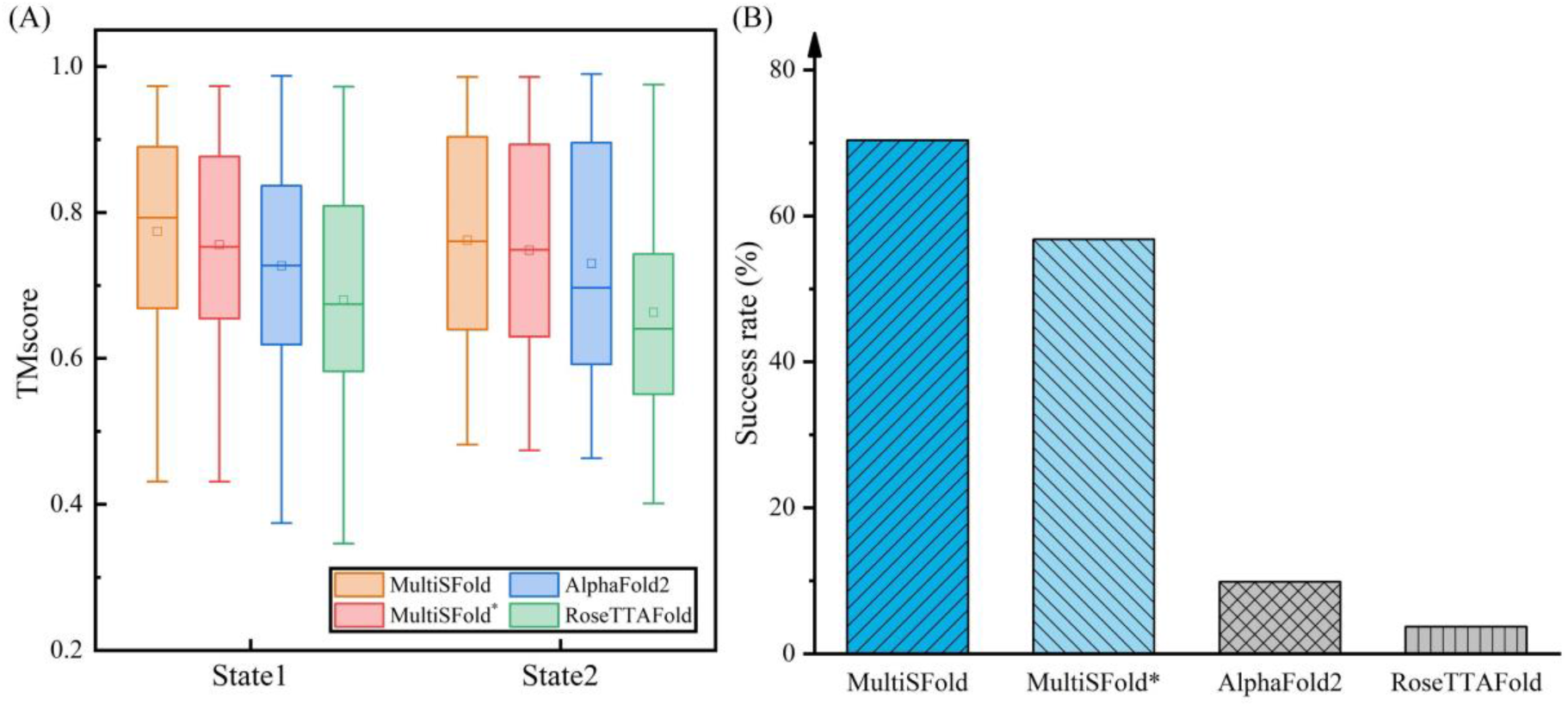
The result of predicting multiple conformations. The models of MultiSFold are from the final population, and the models of MultiSFold^∗^, AlphaFold2 and RoseTTAFold are from the final output of three methods respectively. (A) TM-score boxplot for different methods. The square and black horizontal line in the box represent the average and median TM-scores. (B) Prediction success rate histogram for different methods. The ordinate represents the ratio of successful predictions to all target proteins, and the abscissa indicates different methods.

In order to provide a more intuitive analysis, we proposed a success rate metric to assess multiple conformations modeling performance. For each prediction model, the higher of two TM-scores calculated using two native structures with different conformational states is taken as the final TM-score of the model. The prediction model is then labeled according to the conformational state corresponding to that TM-score. The TM-score of a method for a given conformational state is determined by selecting the maximum TM-score among the prediction models with that state label, provided it exceeds the similarity (TM-score) of the two native structures being compared. If a method produces prediction results for both states of a specific protein and the similarity (TM-score) between the corresponding prediction models is less than 0.9, the method is considered likely to have successfully predicted multiple conformations of that protein. Finally, the success rate of multiple conformation prediction for each method is defined by the percentage of successfully predicted proteins in all targets. The results on the multiple conformation benchmark proteins showed that MultiSFold, MultiSFold*, AlphaFold2 and RoseTTAFold successfully predicted 57, 46, 8, and 3 out of the 81 targets, respectively. As illustrated in **Figure 2B**, the prediction success rate of MultiSFold reached 70.4%, significantly better than that of AlphaFold2 (9.88%) and RoseTTAFold (3.70%).

### 3.3 Comparison with AlphaFold2 and RoseTTAFold on single static structures

MultiSFold is compared with two state-of-the-art methods AlphaFold2 and RoseTTAFold on the single static structure benchmark testset, where templates with a sequence identity ≥30% to the query were excluded. In this test, the distance maps used in MultiSFold are obtained from AlphaFold2, RoseTTAFold and our in-house DeepMDisPre (Zhao et al, 2022), respectively. For each method, the top one model is selected as the final model. In order to objectively evaluate the performance of methods and investigate the complementarity of our method to existing protein structure prediction tools, we divided the test set into 3 subsets, 0-80, 80-90, and 90-100, based on the average pLDDT of the structures in AlphaFold DB. The predicted results of methods on the benchmark testset are summarized in **Table 1**, and the detailed results of each protein are presented in **Supplementary Table S7**.

The results on all benchmark proteins showed that our method outperformed AlphaFold2 and RoseTTAFold, with the average TM-score (0.712) being 2.97% and 7.72% higher than that of AlphaFold2 (0.691) and RoseTTAFold (0.661), respectively. The average RMSD of our method is 8.31 Å, which is reduced by 11.0% and 15.0% compared with that of AlphaFold2 (9.35Å) and RoseTTAFold (9.73 Å), respectively. The *p*-values of student’s t test were 1.58E-09 and 1.46E-25, and the difference was statistically significant. **Figure 3A** presents the head-to-head TM-score and RMSD comparison between methods, where MultiSFold achieved a higher TM-score than AlphaFold2 and RoseTTAFold for 67.6% and 84.4% of total targets, and a lower RMSD for 63.5% and 72.5% of total targets, respectively.

**Fig. 3.**
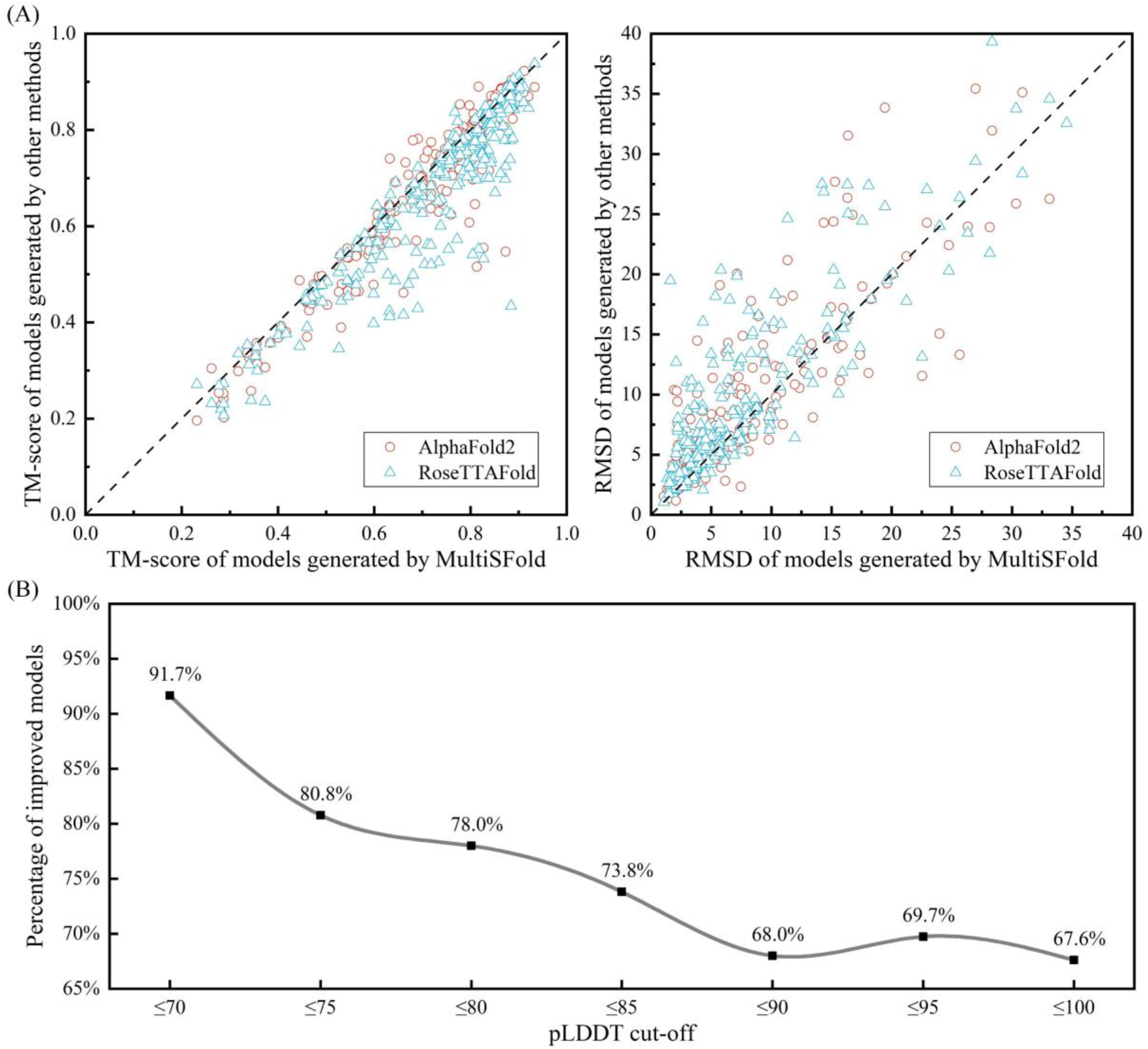
Performance of MultiSFold on the benchmark testset. (A) Head-to-head TM-score and RMSD comparison of MultiSFold with AlphaFold2 and RoseTTAFold. (B) The relationship between the average pLDDT cutoff and the percentage of improved models relative to AlphaFold2.

The results on 3 subsets proteins showed that the average TM-score of our method is 1.65%, 3.11% and 6.17% higher than that of AlphaFold2 in the range of 90-100, 80-90, and 0-80, respectively. Compared with RoseTTAFold, the average TM-score of our method is improved by 6.81%, 7.28%, and 11.41%, respectively. This result suggests that the improvement rate of our method relative to the two comparison methods shows an

upward trend with the decrease of the average pLDDT. In order to clarify this relationship, we further refine the division of the threshold and show the proportion of models improved by our method relative to AlphaFold2 at different average pLDDT cutoffs in **Figure 3B**. With the average pLDDT threshold decreases, our method generated more high-quality models, especially when average pLDDT is less than 80, the percentage of models with higher quality exceeds 75%.

### 3.4 Ablation experiments

We design two ablation experiments to examine the effect of iteration strategy and loop-specific perturbation strategy. The first is MultiSFold-w/o-iteration that uses one round of modal exploration and exploitation. The second is the MultiSFold-w/o-loop, which replaces the loop-specific perturbation strategy with fragment-based modal exploration. The predicted results of different versions of MultiSFold on the benchmark testset are summarized in **Supplementary Table S10**, and the detailed results of each protein are listed in **Supplementary Table S11**.

The average TM-score and RMSD of MultiSFold on all target proteins are 0.712 and 8.31Å, respectively. The average TM-score (0.690) is decreased by 3.09% and the average RMSD (9.418Å) is increased by 13.36%, when the loop-specific perturbation strategy was removed. If only one round of modal exploration and exploitation was considered, the average TM-score (0.678) is decreased by 4.78% and the average RMSD (9.665Å) is increased by 16.25%. Furthermore, **Figure 4A** illustrates the TM-score cut-off threshold of their results, suggesting that MultiSFold outperformed other methods in predicting high-quality structures. Specifically, using a TM-score cut-off threshold of ≥0.8, MultiSFold predicted high-quality structures for 95 out of 244 proteins, while MultiSFold-w/o-iteration and MultiSFold-w/o-loop predicted high-quality structures for only 68 and 78 proteins, respectively. **Figure 4B** presents the results of ablation experiments performed on three subsets, divided based on the average pLDDT of structures in AlphaFold DB. The outcomes demonstrate that the application of both strategies was effective in enhancing prediction accuracy across all subsets.

**Fig. 4.**
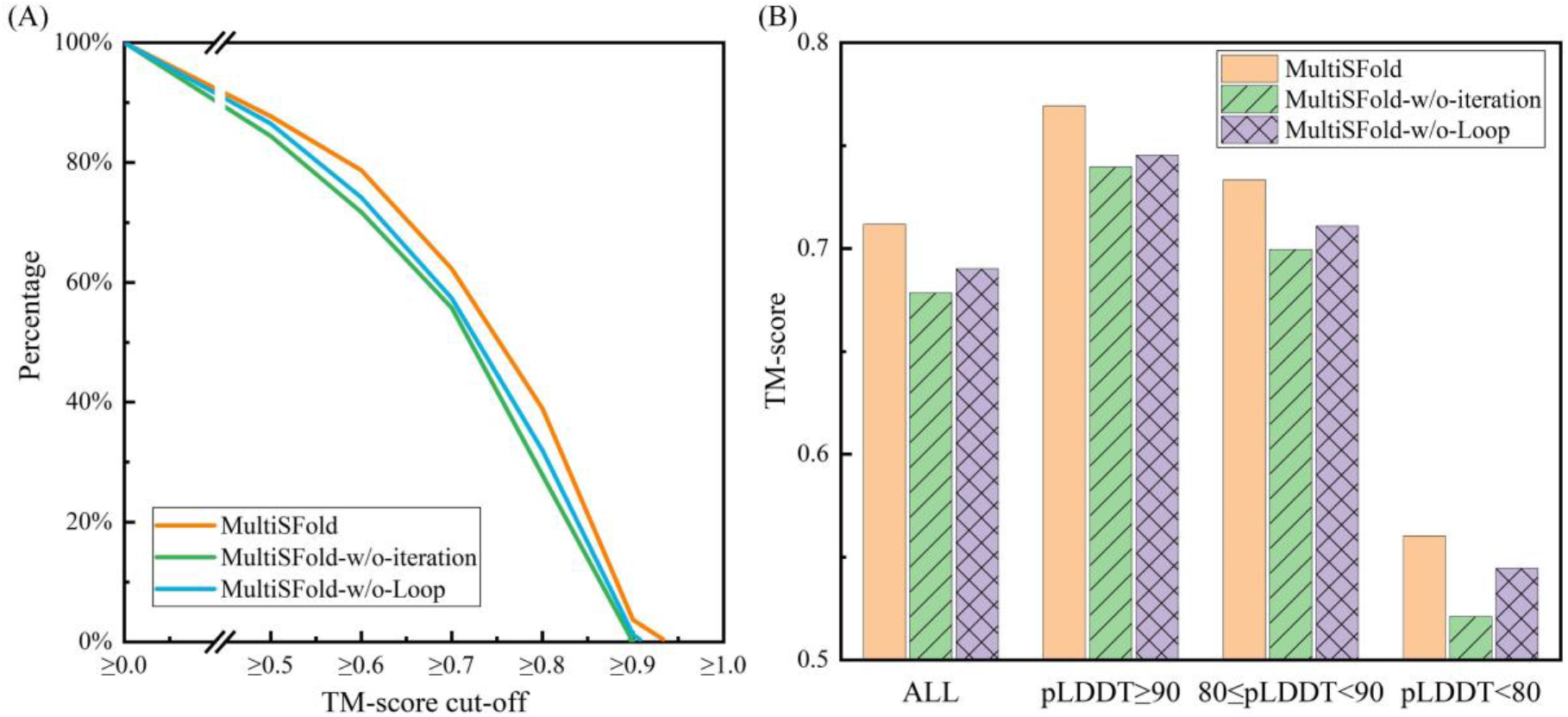
Comparison of MultiSFold, MultiSFold-w/o-iteration and MultiSFold-w/o-loop. (A) Percentage of models at different TM-score cut-off thresholds. (B) Results on the benchmark testset and three subsets divided based on the pLDDT of the structures in AlphaFold DB.

### 3.5 Results of modeling conformational transitions

The Shwachman-Bodian-Diamond syndrome protein (SBDS), which is deficient in the inherited leukemia-predisposition disorder Shwachman-Diamond syndrome (SDS) (Boocock et al, 2003), cooperates with the GTPase elongation factor-like 1 (EFL1) to activate nascent 60S ribosomal subunits for translation by catalyzing eviction of the ribosome antiassociation factor eukaryotic initiation factor 6 (eIF6) (Senger et al, 2001; Finch et al, 2011). Upon EFL1 binding, SBDS is repositioned around helix 69, thus facilitating a conformational switch in EFL1 that displaces eIF6 by competing for an overlapping binding site on the 60S ribosomal subunit (Weis et al, 2015). As illustrated in **Figure 5A**, SBDS domain II undergoes a 60° rotation with a pivot point through the N terminus of helix α5 and domain III rotates 180°, thus transferring the conformational structure from the "closed" state to the "open" state. For AlphaFold2, only the "open" conformation was successfully modeled and the "closed" conformation was not, while RoseTTAFold only successfully predicted the "closed" conformation. Interestingly, MultiSFold successfully modeled the "open" and "closed" conformations using the distance constraints from two methods, and achieves better TM-scores of 0.638 and 0.703 respectively.

**Fig. 5.**
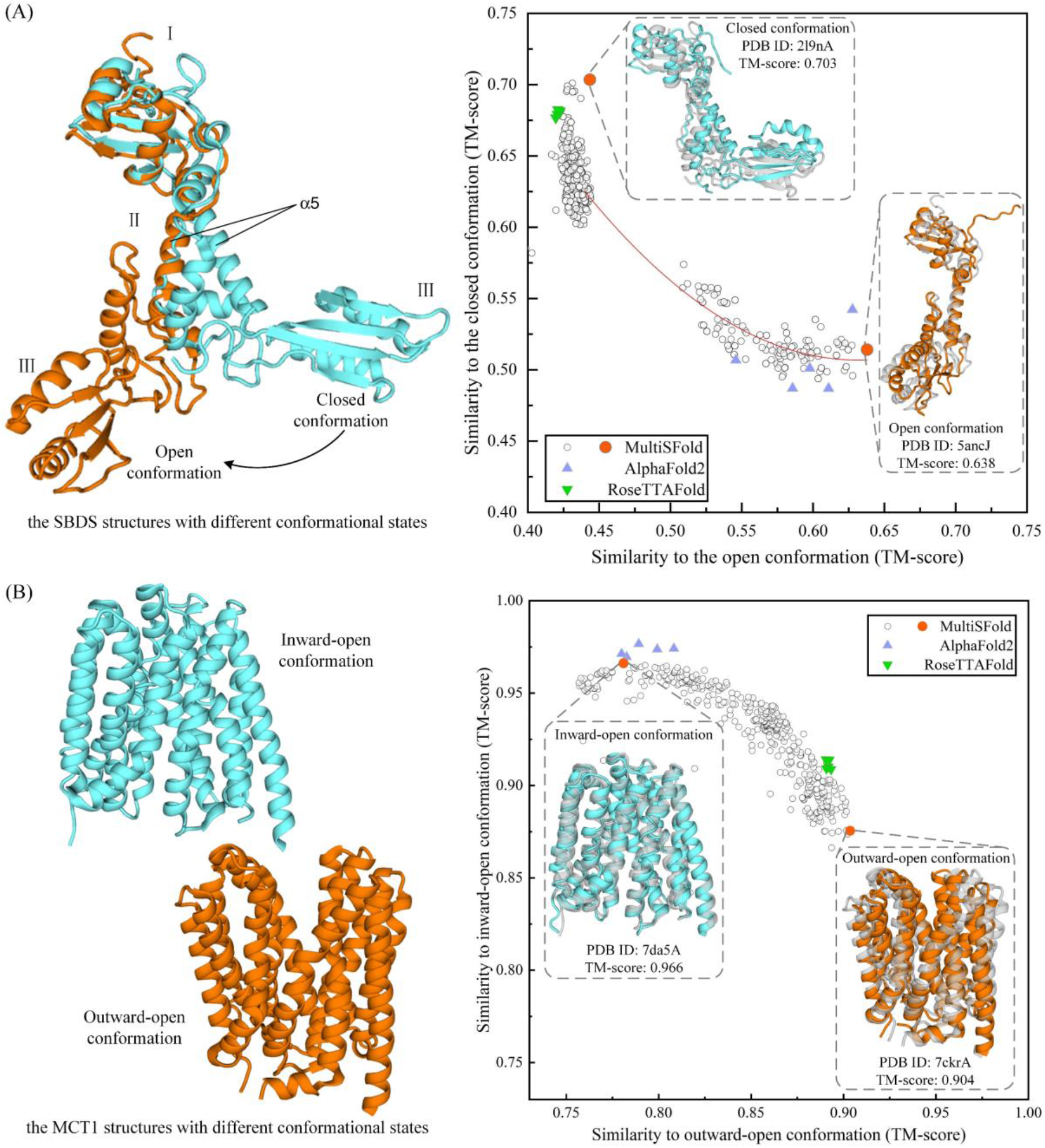
Two representative examples of modeling conformational transitions. (A) Representative models of the protein SBDS in closed and open conformations. (B) Representative models of the transporter MCT1 in inward-open and outward-open conformations. The experimental structures of proteins with different conformational states are shown in the left. The right scattergram shows the comparison of prediction models with closed/open or inward-open/outward-open experimental structures, experimental structures shown in gray and prediction model shown in color.

As the gatekeepers and surveillance systems of the cells, membrane proteins often use conformational changes to regulate transport activity or signaling to soluble messengers. Monocarboxylate transporter 1 (MCT1, alias SLC16A1) is a proton coupled monocarboxylate transporter that catalyzes the movement of many monocarboxylates, such as lactate and pyruvate, across the plasma membrane through an ordered mechanism (Garcia et al, 1994; Ritzhaupt et al, 1998). This substrate translocation is achieved through rigid-body rotation of the two domains, which forms two distinct conformational states (inward-open and outward-open, as show in **Figure 5B**) and exposes the central substrate binding sites alternatingly to either side of the membrane (Wang et al, 2021). For AlphaFold2, only the "inward-open" conformation was successfully modeled and the "outward-open" conformation was not, while RoseTTAFold only successfully predicted the "outward-open" conformation. MultiSFold successfully modeled the "inward-open" and "outward-open" conformations, and achieves TM-scores of 0.966 and 0.904 respectively.

## 4 Conclusion

The protein structure prediction has made remarkable progress with the arrival of deep learning end-to-end techniques such as AlphaFold2. However, although AlphaFold2 can accurately predict the conformation of main chain and side chain, it is specific to the specific conformation state (Skolnick et al, 2021). For the proteins with multiple conformational states, the AlphaFold2 models do not provide insight into conformational dynamics (Subramaniam and Kleywegt, 2022). In this work, we try to produce multiple conformations containing different states through conformational sampling combined with knowledge gained through deep learning. In addition, multiple competing constraints may help to further improve the accuracy of a single static structure.

This work develops a multiple conformational states folding method using the distance-based multi-objective evolutionary algorithm, named MultiSFold. For the query sequence, the method first constructs multiple competing constraints from the distance information generated in different ways. The initial population is generated by randomly fragment assembly using the variable-length fragment library generated by in-house method VFlib. Then, the population is updated by the iteration of a multi-objective optimization strategy for modal exploration and a geometric optimization strategy for modal exploitation. Finally, a loop-specific perturbation strategy is used to adjust the spatial orientation between the secondary structures to generate the final population.

MultiSFold was tested on a developed benchmark testset, which contains 81 proteins with two representative conformational states. Experimental results show that the prediction success rate of MultiSFold reached 70.4%, significantly better than that of AlphaFold2 (9.88%) and RoseTTAFold (3.70%). On two representative cases, SBDS and MCT1, MultiSFold successfully predicted both two conformational states, while AlphaFold2 and RoseTTAFold can only predict one of them. This may be due to the fact that multiple competing constraints cover different conformational states information, and the multi-objective optimization algorithm makes it have the ability to span the range between two conformational states. More interestingly, the ensemble generated by MultiSFold may represents possible intermediates on a transition pathway between two states, which may help to understand the mechanism of conformational transitions (Zacharias, 2017). In addition, complete information with multiple competitive constraints may reduce the noise of a single constraint and further improve the prediction accuracy of a single static structure. The experimental results on 244 human proteins from AlphaFold DB show that the MultiSFold models obtained better average TM-score than AlphaFold2, especially when average pLDDT is less than 80.

While these results seem to support to some extent the notion that knowledge gained through deep learning can guide conformational sampling to predict multiple conformational states, the information of distinct conformational states is not an explicit objective built into the existing deep learning methods. It may be important to further develop artificial intelligence methods capable of learning the conformational flexibility intrinsic to protein structures, which is also a direction of our research exploration.

## 5 Acknowledgments

This work is supported by the National Key R&D Program of China (2022ZD0115103), the National Nature Science Foundation of China (62173304), and the Key Project of Zhejiang Provincial Natural Science Foundation of China (LZ20F030002).

## Supplementary Texts

### Text S1. Details of MultiSFold

The framework of MultiSFold is described as **Figure S1**. The sequence and several distance maps are used as input, and MultiSFold finally outputs the predicted three-dimensional structures of the target sequence.

**Figure S1.**
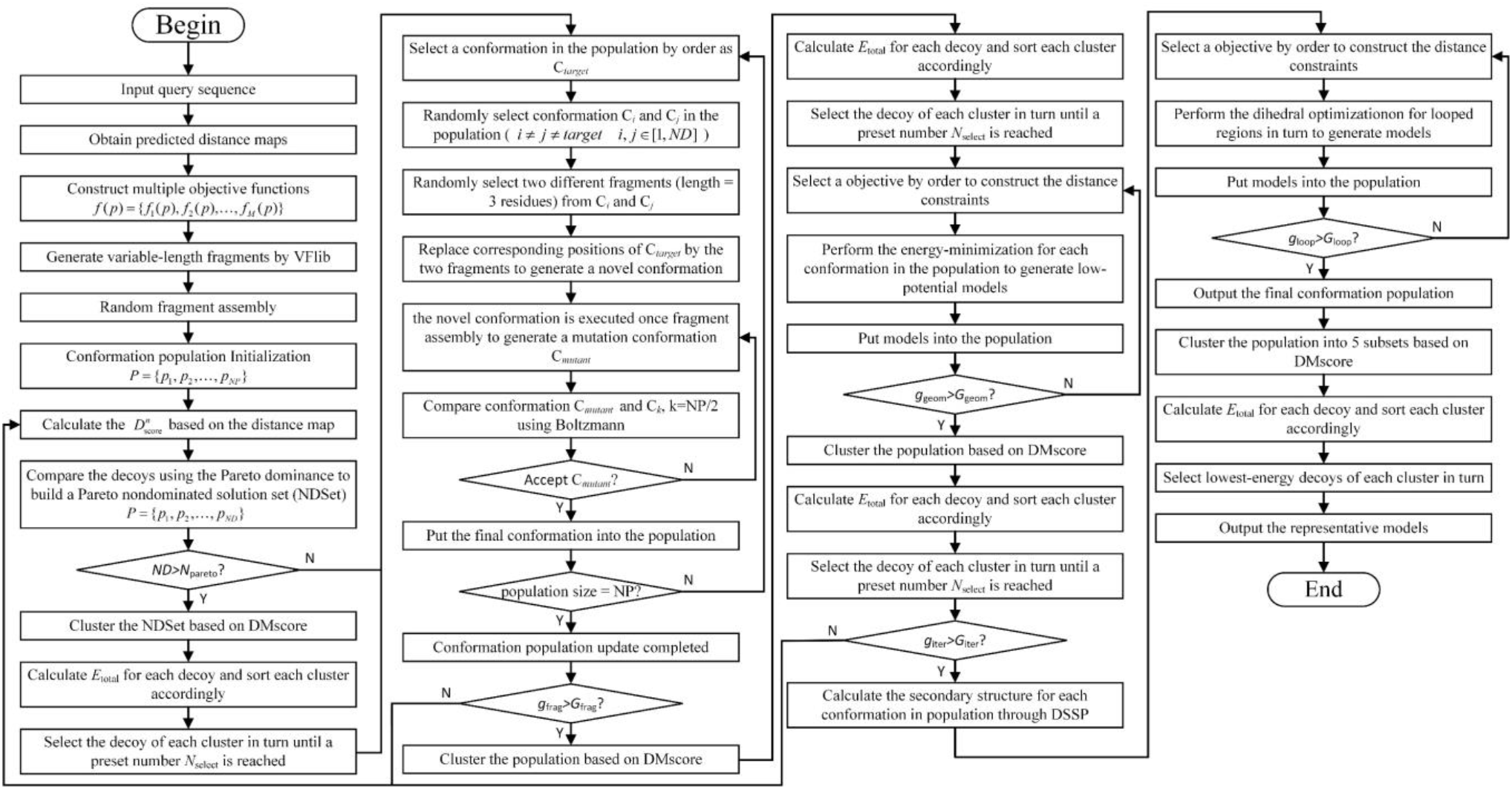
The framework of MultiSFold.

#### The initialization of MultiSFold

First, the meta distance map is generated based on several input distance maps, and multiple objective functions *f* (*p*) ={ *f*_1_ (*p*), *f*_2_ (*p*), ..., *f_M_* (*p*)} were constructed based on all distance maps. Then, the fragments with variable length of the query sequence will be generated through our previously developed method VFlib ^[S1]^. Finally, the initial population *P* ={ *p*_1_, *p*_2_, …, *p_NP_*} is generated through random fragment assembly.

#### The modal exploration and exploitation of MultiSFold

This stage is mainly for the process of fragment-based modal exploration and geometric-based modal exploitation, and these two processes perform *G*_iter_ times of iterations.

In the process of fragment-based modal exploration, the 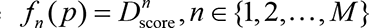 of each decoy will be firstly obtained based on a scoring model using multiple distance maps. Then, the Pareto dominance is used to compare the decoys in population, which is illustrated as follows: A solution *p*_1_ ∈ *P* is said to (Pareto) dominate another solution *p*_2_ ∈ *P*, if

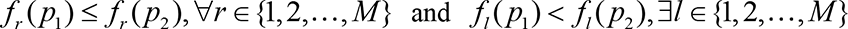

where *r* and *l* represent the index of objective functions. A solution *p*^∗^ ∈ *P* is called a Pareto minimizer or a nondominated solution, if there does not exist another solution that dominates it; that is, there exists no other solution that would decrease some objectives without causing simultaneous increase in at least one other objective. The nondominated decoys (the solution *p*^∗^) will be selected from the population to build a Pareto nondominated solution set. If the number *ND* of nondominated solutions (NDSet) exceeds the preset threshold *N*_pareto_, the DMscore-based clustering will be performed which is described in the **Supplementary Text S2**, and the decoys of each cluster will be sorted by the *E*_total_. Each sorted cluster will be selected in turn and the lowest energy conformation of that cluster be extracted into the population until a preset number *N*_select_ is reached. Based on the selected decoys, fragment recombination and assembly are performed to generate new decoys for populating the population.

In fragment recombination, each preserved decoy in the population will be selected in turn as the target conformation, *C*_*target*_. Two mutually different decoys are randomly selected, namely, *C*_*rand*1_ and *C*_*rand*2_, where *rand*1 ≠ *rand*2 ≠ *target*. Afterward, two mutually different fragments (length 3 residues) are randomly selected from *C*_*rand*1_ and *C*_*rand*2_, and then the corresponding positions of *C*_*target*_ are replaced by the two fragments to generate a novel conformation. Then, the novel conformation is executed fragment assembly. In fragment assembly, select different fragment libraries based on the number of the current iteration *g*_frag_. When the number of the current iteration *g*_frag_ is less than 1/5 of the maximum number of iterations *G*_frag_, one fragment in the 6 to 21-mer fragment libraries is randomly selected to replace the corresponding position; otherwise, the 3 to 5-mer fragment libraries is randomly selected to perform the operation. Finally, the mutation conformation *C*_mutant_ is generated and compared with target conformation *C*_*target*_ and received through Metropolis Monte Carlo. After filling up the population, it will enter the next iteration until the number of iterations is greater than the preset number *G*_frag_. The DMscore-based clustering and conformation selection are performed to select some representative decoys before modal exploitation to reduce the computational cost.

In the process of geometric-based modal exploitation, each decoy in the population would be selected as the base conformation, and the energy-minimization is applied *G*_geom_ times to generate low-potential models guided by multiple distance constraints. The DMscore-based clustering and conformation selection are performed to select some representative decoys from these low-potential models.

#### The Loop-specific sampling

According to multiple inter-residue distance, a loop-specific dihedral angle optimization strategy is used to adjust the topology efficiently. For each conformation, the secondary structure is first calculated by DSSP, and then the dihedral optimization is performed in turn for the loop regions connected with α-helix or β-sheet at both ends. The aim of this dihedral optimization is to find a set of rotation angles acting on the loop regions, to generate a conformation that satisfies the distance constraint to the greatest extent. In this strategy, multiple constraints are used to guide the dihedral optimization to produce more diverse models in order to span different conformational states. The details of the loop-specific dihedral angle rotation model construction and the dihedral optimization are shown in **Supplementary Text S3**. Finally, the DMscore-based clustering and conformation selection are performed to select five representative models and the lowest-energy conformation is selected as the prediction model.

**The parameters of MultiSFold** are listed in **Table S1**.

**Table S1.** Parameter descriptions in MultiSFold.

| Parameter | value | descriptions |
| --- | --- | --- |
| $M$ | 4 | Number of objective functions |
| $NP$ | $25*(M+2)$ | Population size |
| $N_{\text{pareto}}$ | $0.8*NP$ | Number threshold of nondominated decoys |
| $N_{\text{select}}$ | $5*(M+2)$ | Number of selected decoys |
| $G_{\text{iter}}$ | 3 | Generation of modal exploration and exploitation |
| $G_{\text{frag}}$ | 200 | Generation of fragment-based modal exploration stage |
| $G_{\text{geom}}$ | $M$ | Generation of geometric-based modal exploitation stage |
| $G_{\text{loop}}$ | $M$ | Generation of loop-specific sampling stage |

### Text S2. DMscore-based cluster

DMscore ^[S2]^ measures the similarity by calculating the difference of Euclidean distance of the corresponding residue pairs in two models, which is defined as follows:

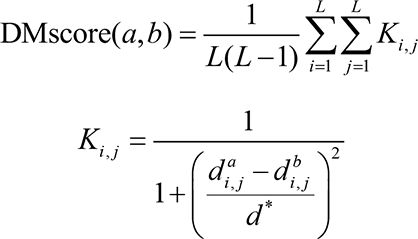

where *L* is the length of the protein structure; 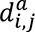 and 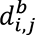 are the distances between the *i*-th and *j*-th residues in model *a* and model *b*, respectively; and *d\** is the normalized scale used to eliminate the inherent dependence of the score on protein size, which is defined as the logarithm of the residue sequence distance, namely:

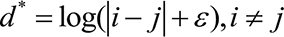

where *ε* is an infinitely small quantity to avoid *d\** being zero. The value of DMscore is between (0,1]. The higher value indicates higher similarity between the two models; 1 indicates a perfect match between two models.

A clustering algorithm is designed to adaptively determine the number of clusters according to the multi-order nearest distance analysis. The number of clusters is adaptively counted by step information. Then, the DMscore between the conformations is taken as the distance and clustered. 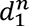 is the first-nearest neighbor distance of the *n-*th conformation in the population, 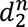 is the second-nearest neighbor distance, and so on. These distances satisfy:

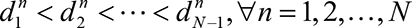

where *n* is the index of the decoy in population; *N* is the size of population. The mean square value and the mean value of the *m-*th nearest neighbor distances of all conformations are calculated as follows:

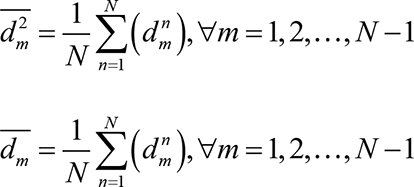

where *m* represents the nearest neighbor order number. The distribution of individuals determines the number of divisible populations. The distribution degree of the population is analyzed by calculating the variance, as follows:

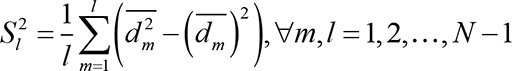

where *m* and *l* represent the *m-*th and the *l-*th nearest neighbor distances, respectively. The **Figure S2** intuitively shows first-order shortest neighbor distance and second-order shortest neighbor distance.

**Figure S2.**
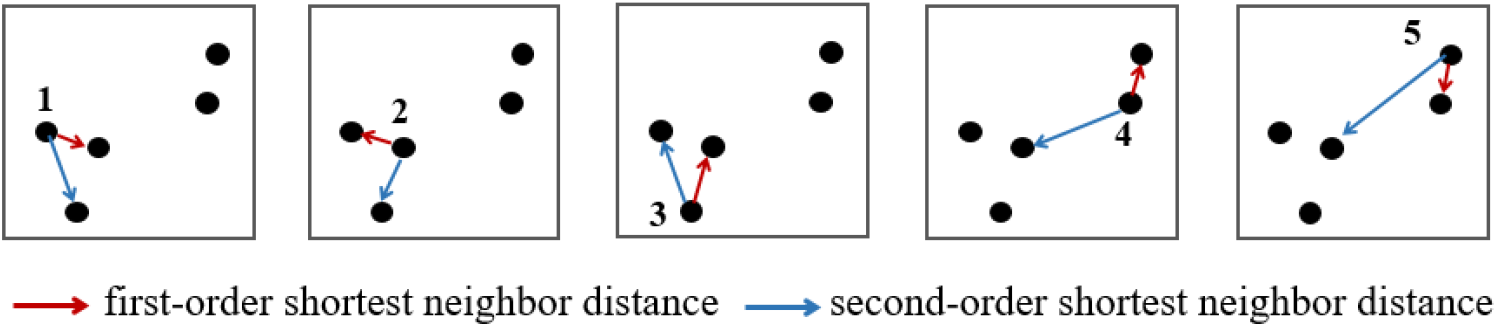
Schematic schematic diagram of first-order shortest neighbor distance and second-order shortest neighbor distance.

In order to visually analyze the cluster algorithm, the example in two dimensions was shown in the **Figure S3**. **Figure S3(A)** is the single cluster, **Figure S3(B)-(D)** are multiple clusters. Through the comparison, it can be found that once there is more than one population, the variance calculated by the nearest neighbor distance will have a corresponding step. This is because of that the distance between individuals within the same population varies slightly, but the distance between individuals varies greatly when crossing from one population to another. Therefore, the slope of the variance of the m-th nearest neighbor distance will fluctuate greatly. In this way, all the step information is captured and the global distribution of the current population can be obtained.

**Figure S3.**
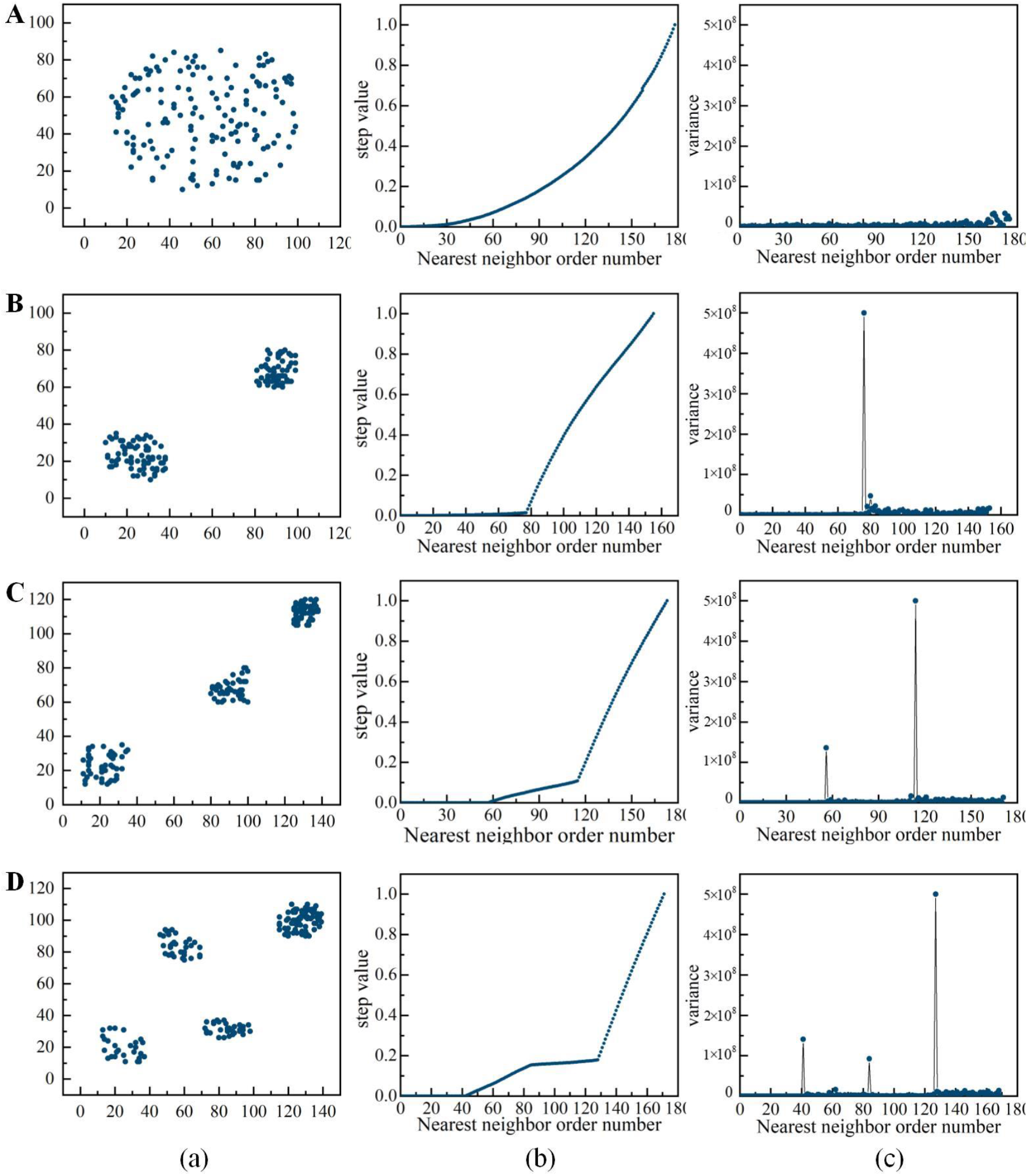
Schematic of DMscore-based cluster. (a) Population distribution. (b) The relationship between the nearest neighbor order number and the variance. (c) The slope of the variance changes.

### Text S3. Loop-specific sampling

In this strategy, multiple distance constraints are constructed firstly based on different objectives. Then, the secondary structure for each conformation is calculated by DSSP, and the dihedral optimization is performed in turn for the loop regions connected with α-helix or β-sheet at both ends. For the selected loop region, a loop-specific dihedral angle rotation model is constructed as shown in **Figure S4**. The set of dihedral angle rotation axes Γ of the loop region is defined as follows ^[S3, S4]^:

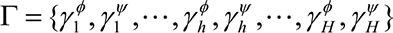

where 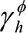 and 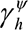 are the unit axes of the N-C_α_ and C_α_-C atomic bonds of the *h*-th residue in the selected loop region, corresponding to the dihedral angles *ϕ* and *ψ*, respectively. *H* is the number of residues in the selected loop region.

**Figure S4.**
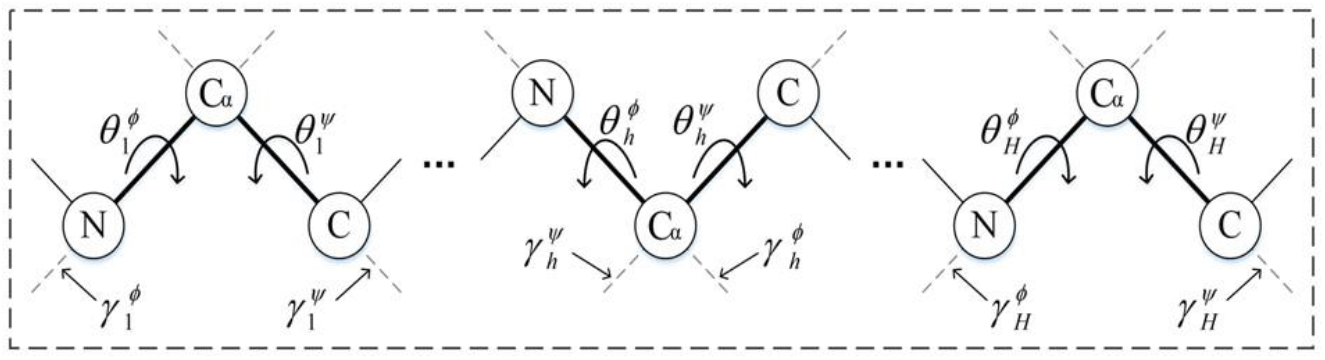
The dihedral angle rotation model of loop region.

The inter-residue distance with residues on both sides of the loop region are selected as the constraints. The coordinates of the C_β_ (C_α_ for glycine) atomics of the *m*-th selected residue pair are expressed as 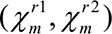, and the coordinate transformation and differential evolution algorithm are used to generate a set of rotation angles 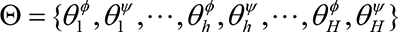. The rotation matrix of the coordinate transformation for the rotation points can be calculated by 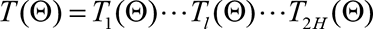, which is defined as follows:

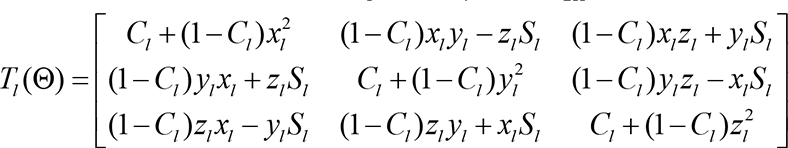

where *T_l_* (Θ) is the rotation matrix corresponding to the *l*-th rotation axis; C*_l_* and S*_l_* are cos*θ_l_* and sin*θ_l_*, respectively; *θ_l_* is the *l-*th rotation angle; (x*_l_*; y*_l_*; z*_l_*) is the unit vector of the *l*-th rotation axis; and *l* ∈{1, 2,…, 2*H*}.

With the assumption that the residues on the left of the loop region remain fixed and the residues on the right are rotated, the coordinates of 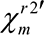 obtained by the rotation transformation of 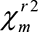 can be calculated by 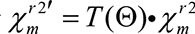.

## Supplementary Figures

**Figure S5.**
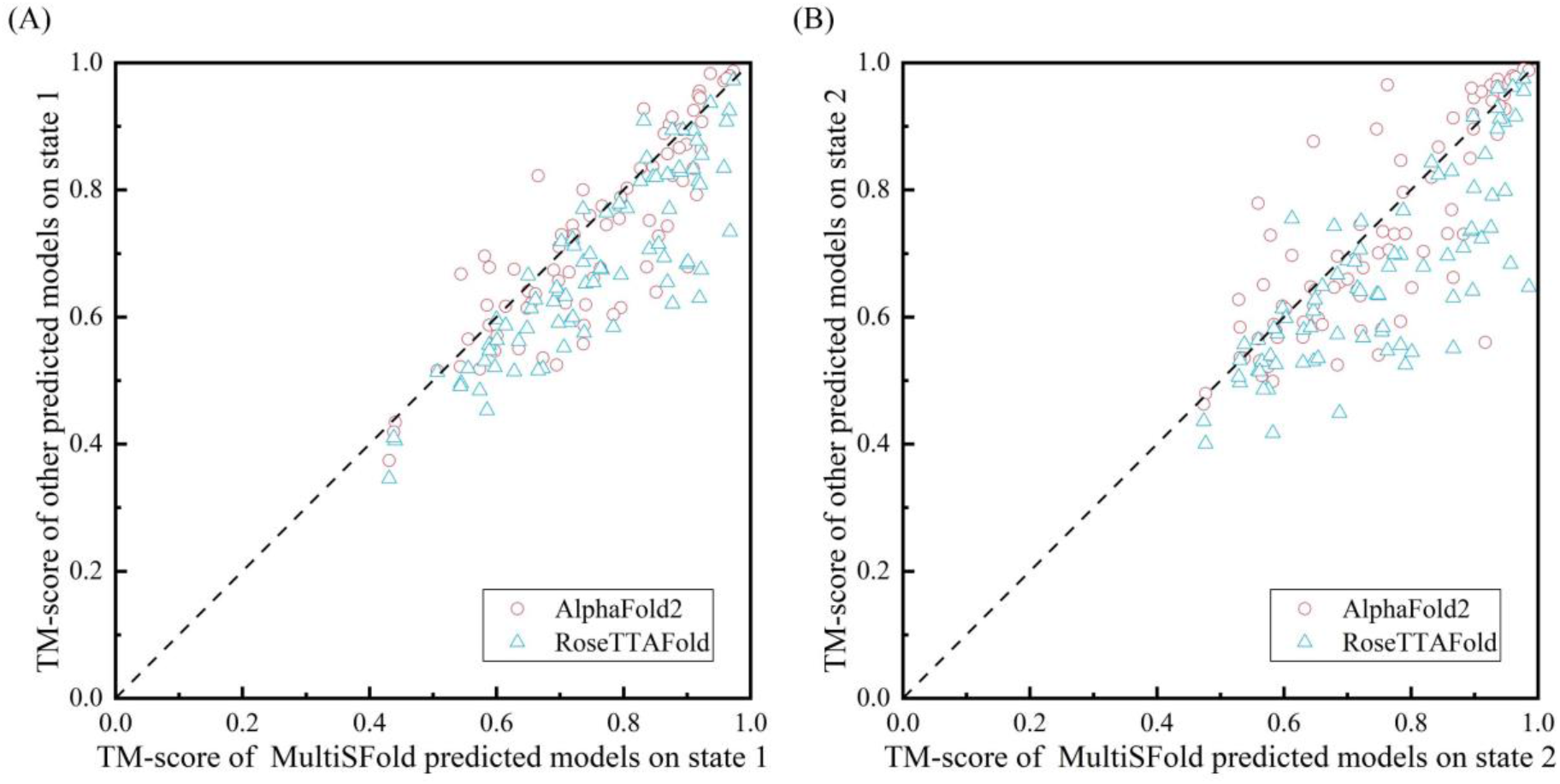
Head-to-head TM-score comparison of MultiSFold with AlphaFold2 and RoseTTAFold on distinct conformational states of 81 proteins.

**Figure S6.**
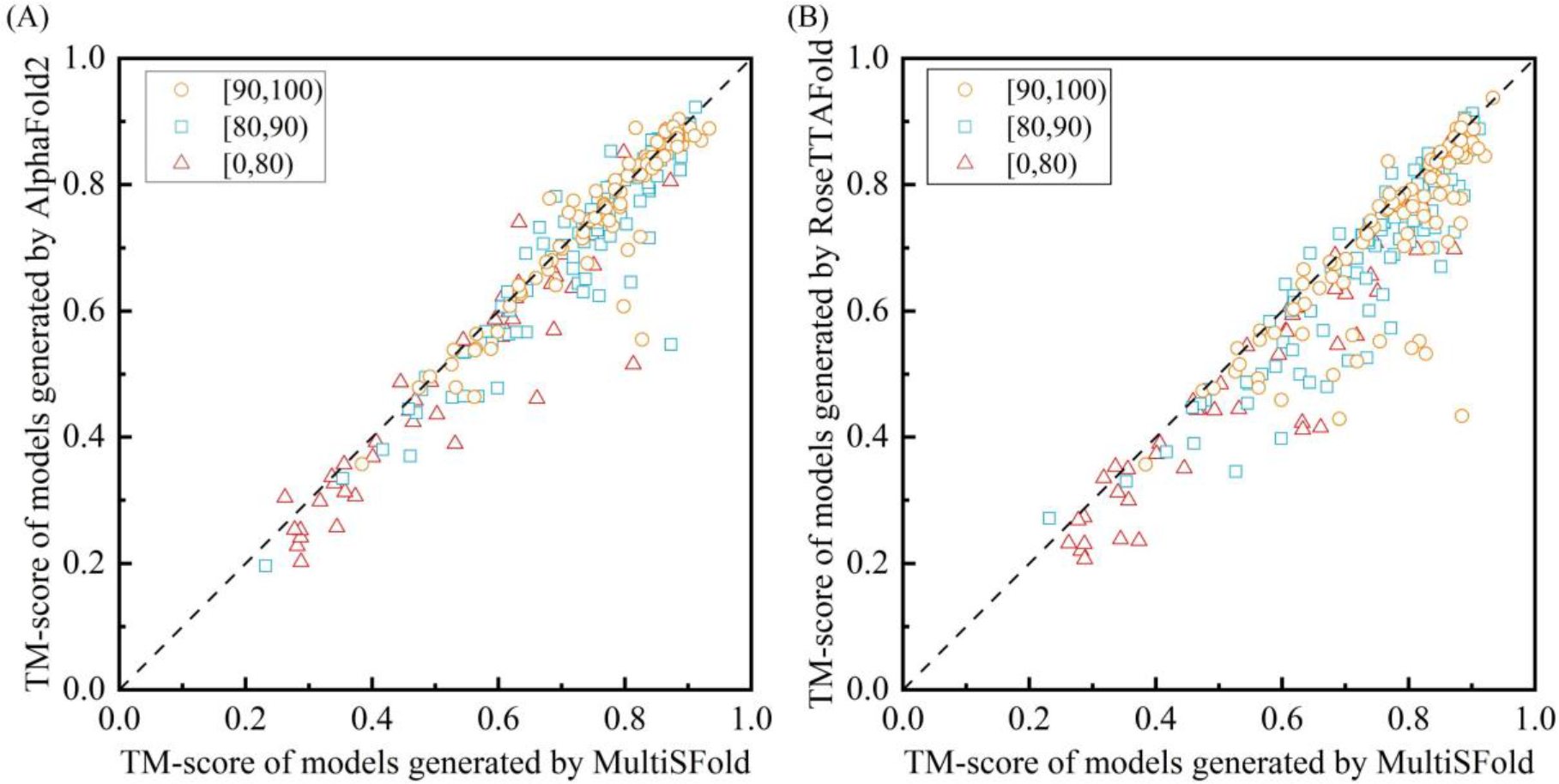
Performance of MultiSFold on the 244 human proteins. (A) Head-to-head TM-score comparison of MultiSFold with AlphaFold2 on three subsets, 0-80, 80-90 and 90-100, which are divided based on the average pLDDT of the structures in the AlphaFold DB. (B) Head-to-head TM-score comparison of MultiSFold with RoseTTAFold on three subsets.

## Supplementary Tables

**Table S2.**
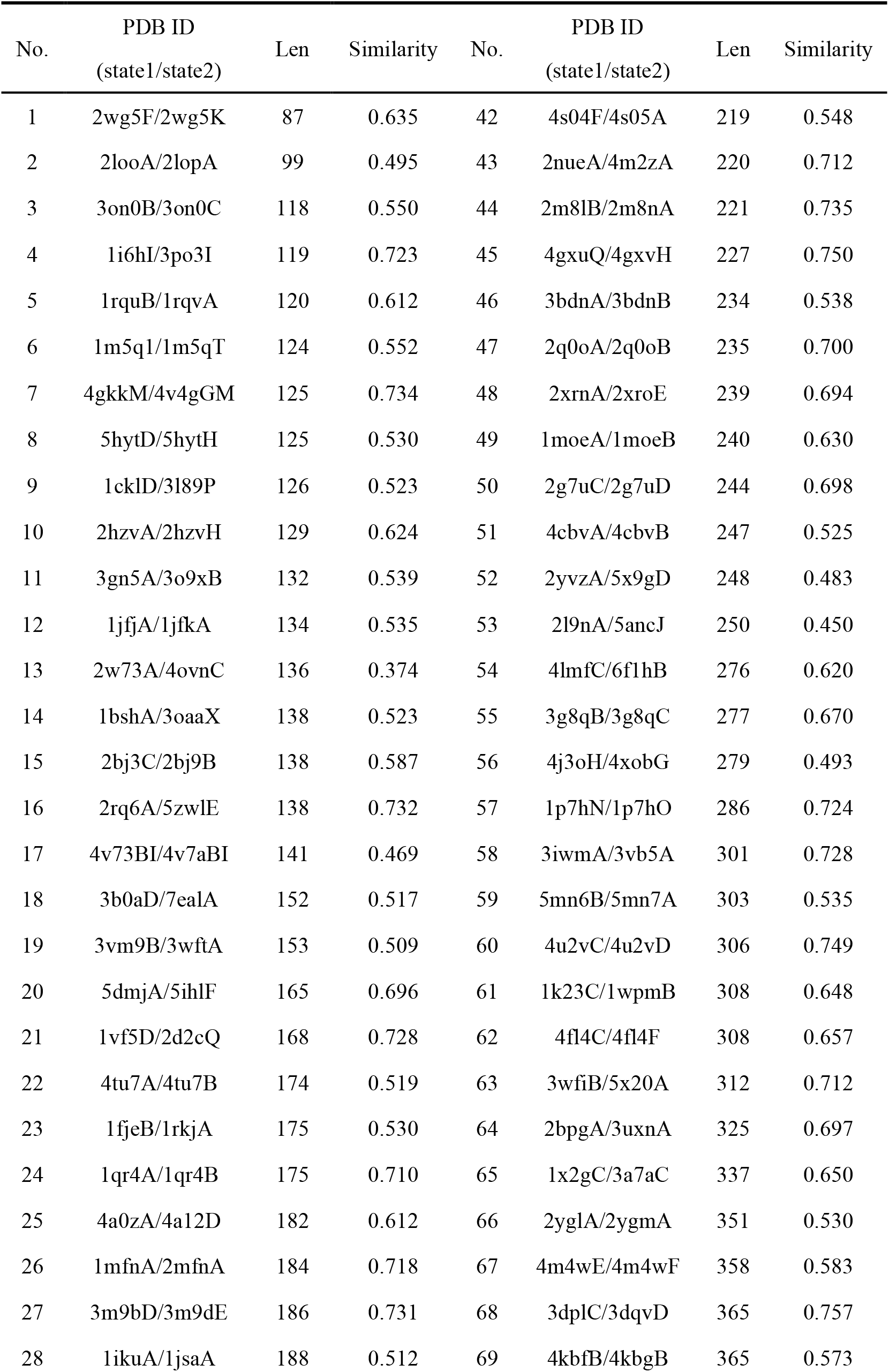
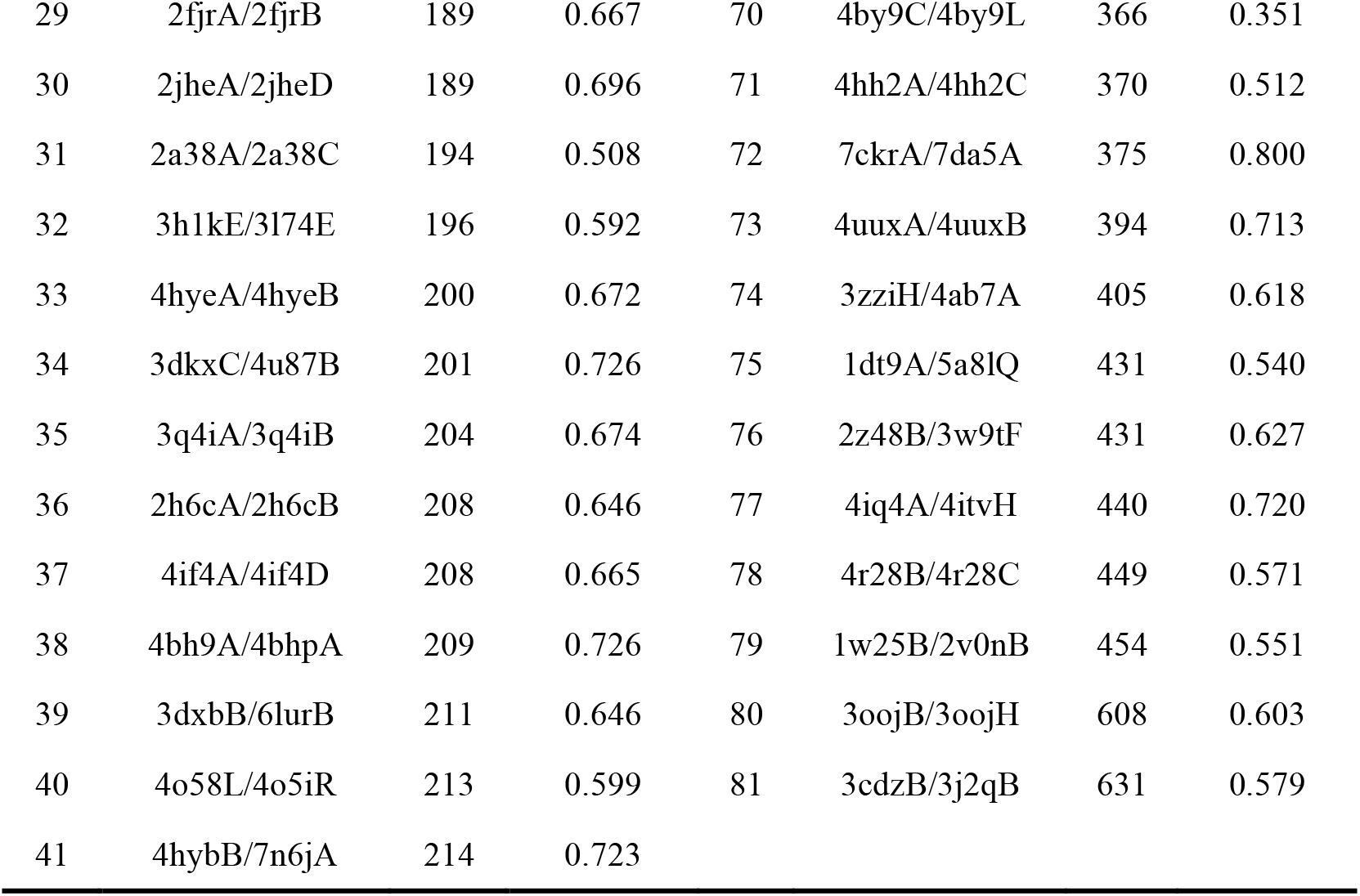
Details of 81 proteins with multiple conformational states.

**Table S3.**
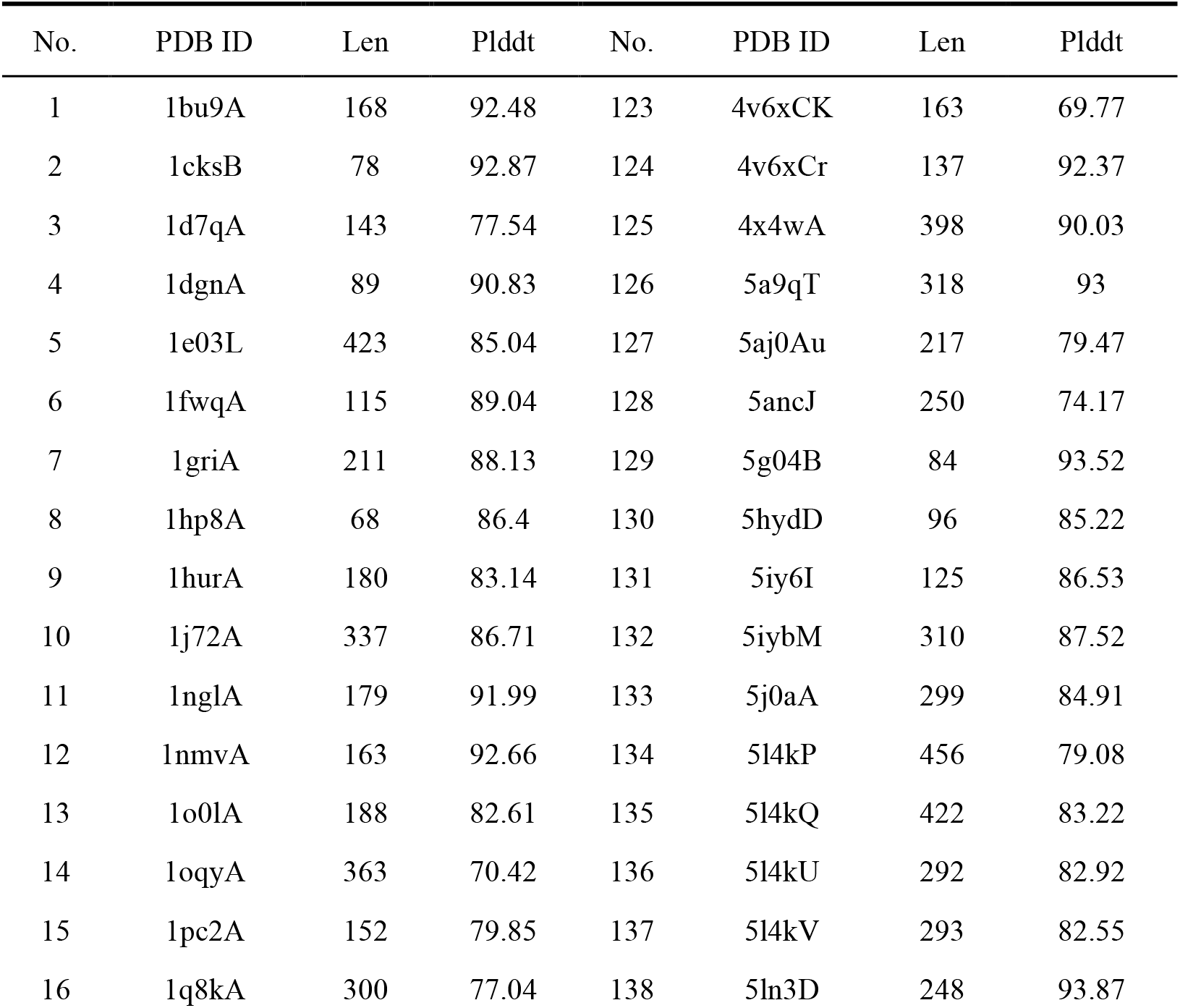
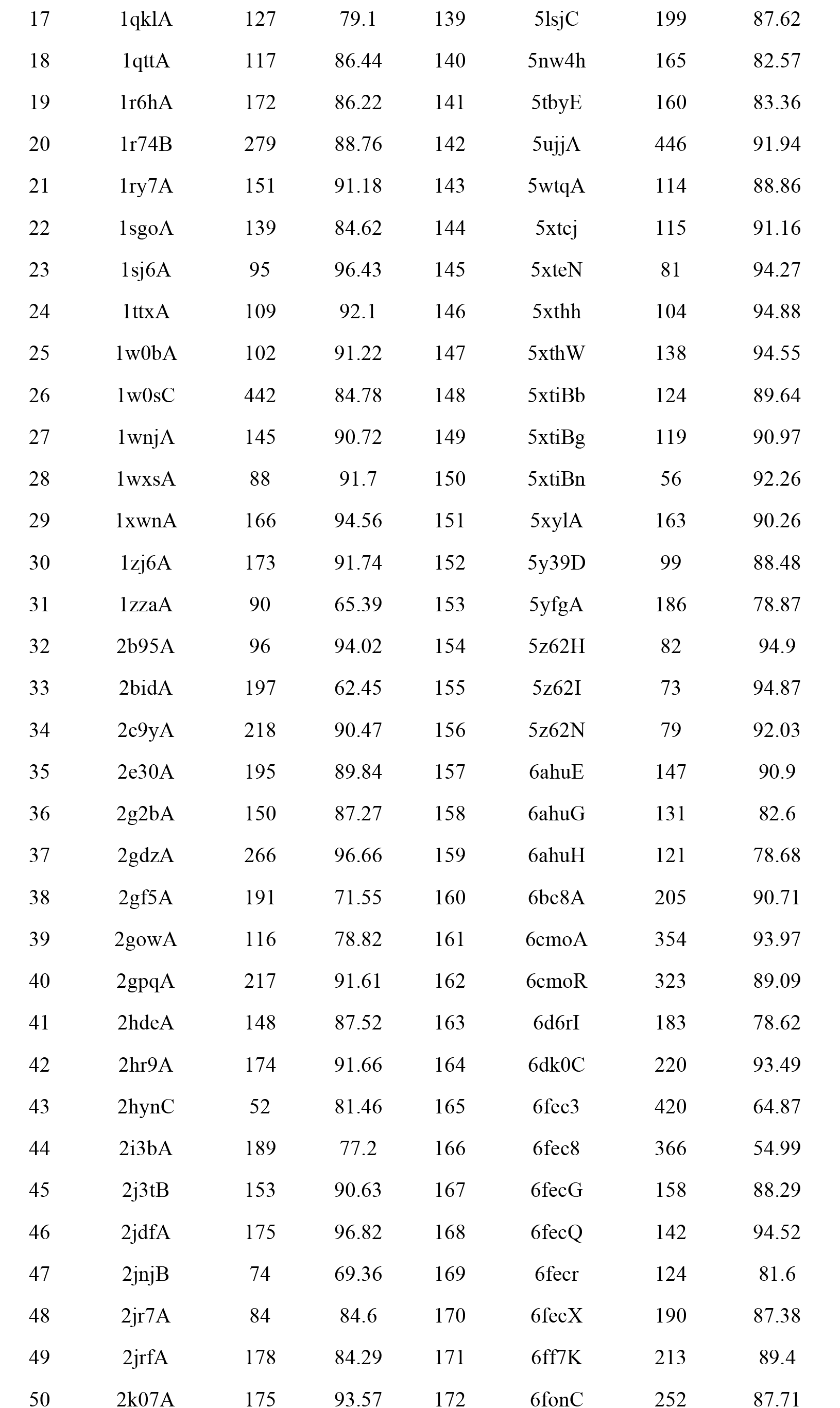
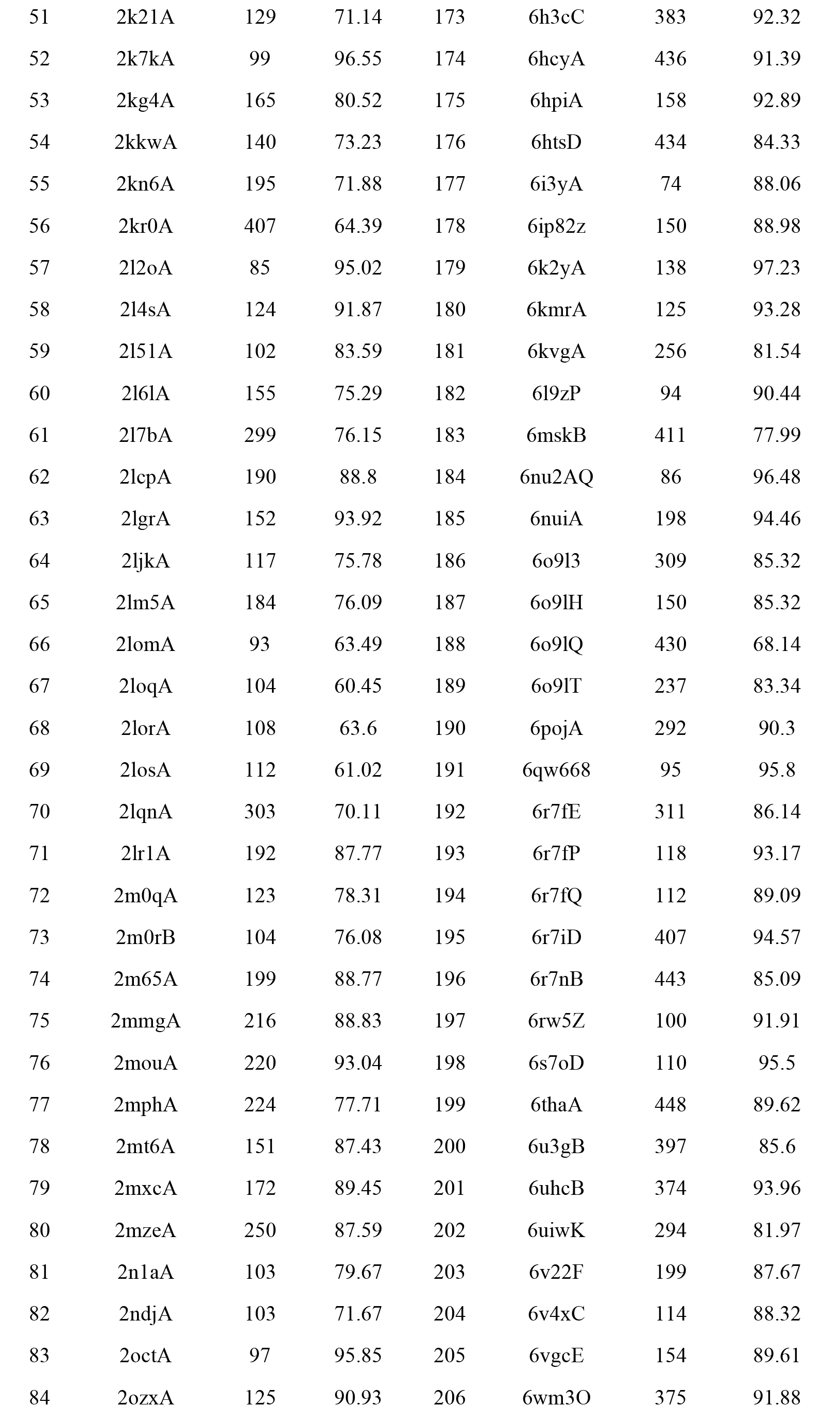
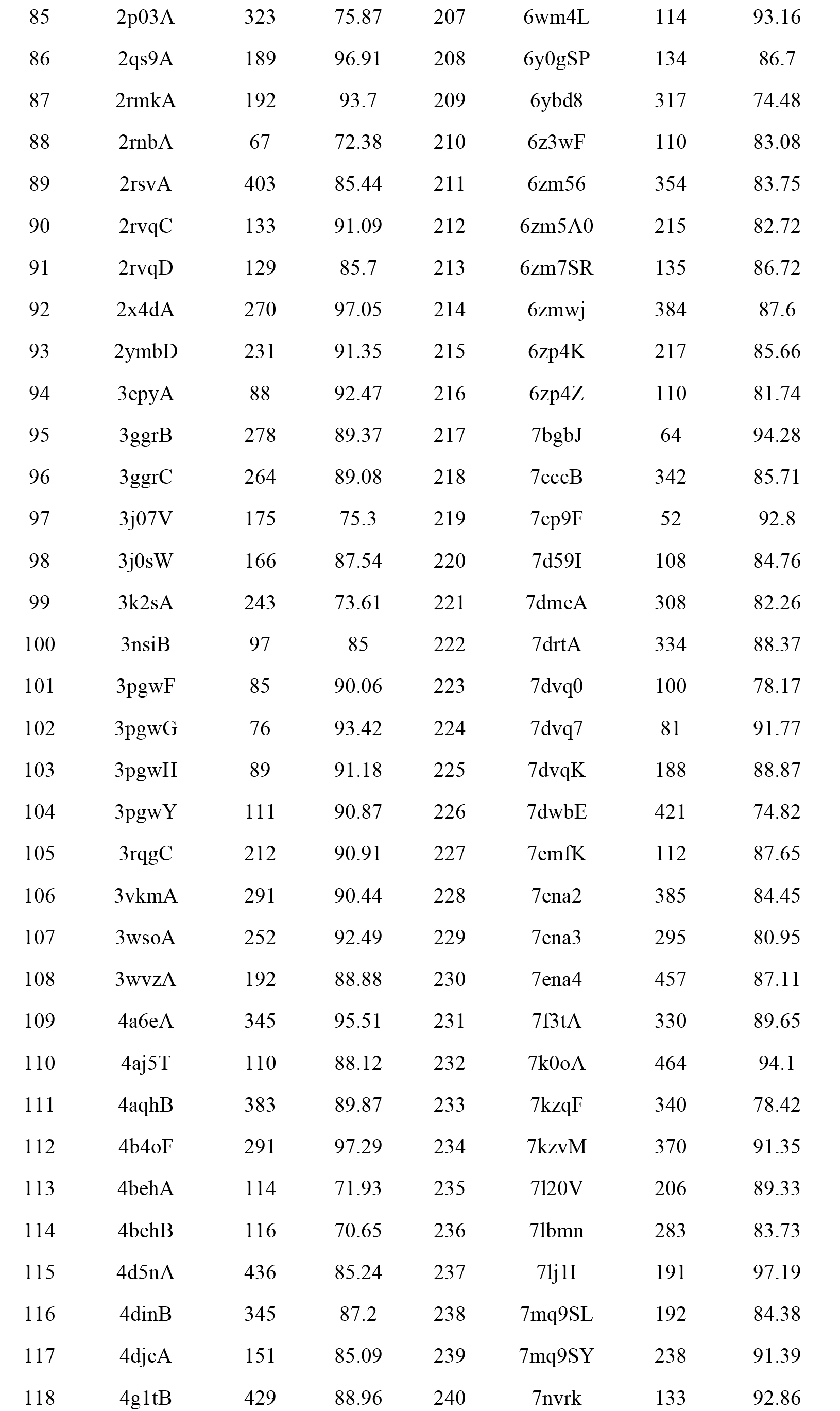
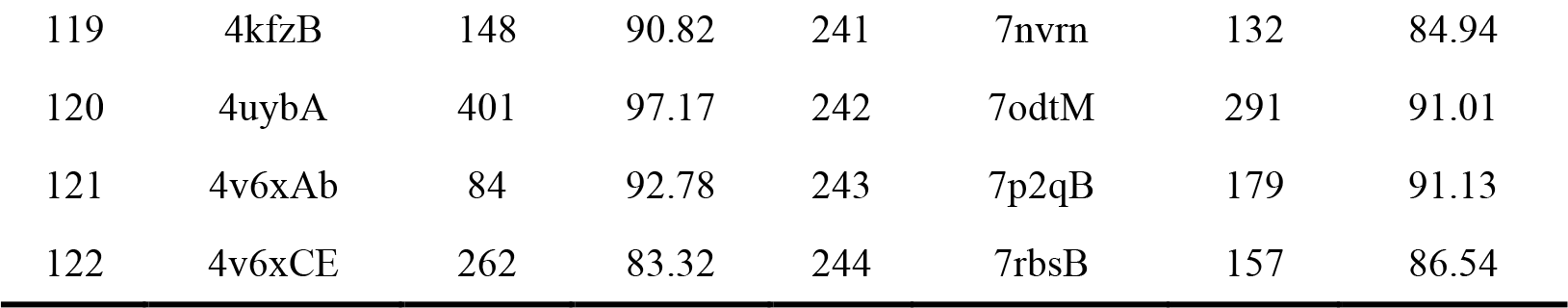
Details of 244 human proteins from AFDB.

**Table S4.**
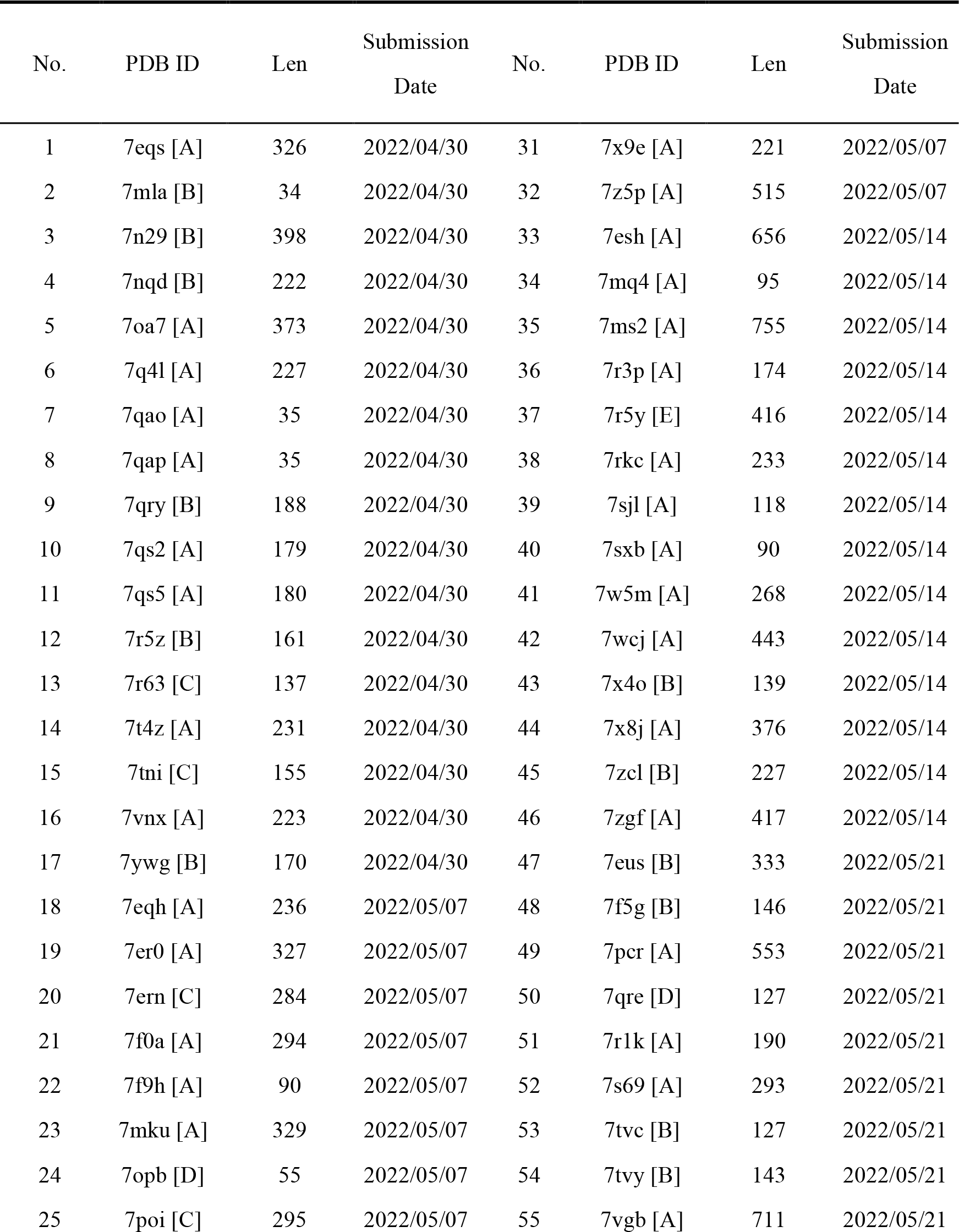
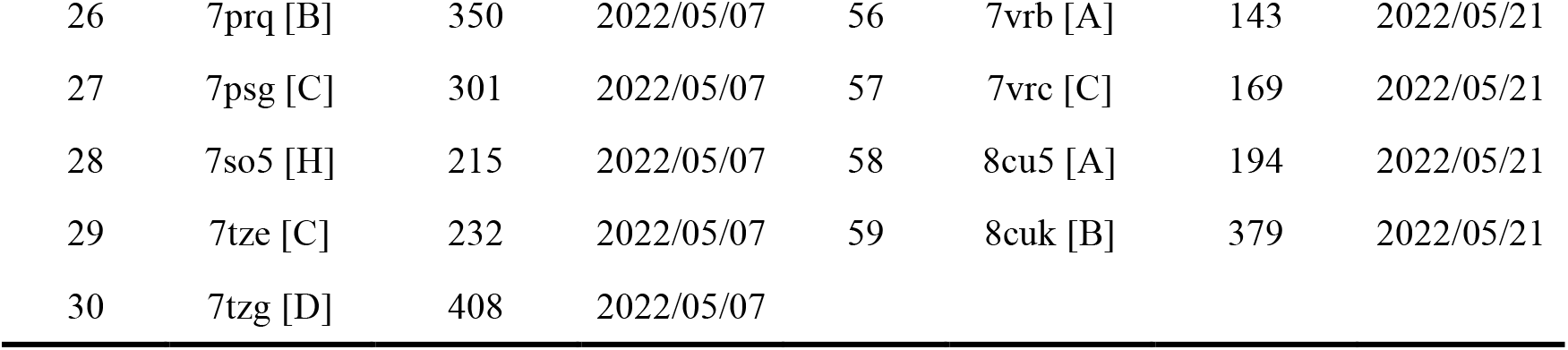
Details of target proteins from CAMEO (2022/04/30∼2022/05/21).

**Table S5.** Summary of experimental results for predicting multiple conformations.

| Method | State1 |  |  | State2 |  |  | Success ratio (%) |
| --- | --- | --- | --- | --- | --- | --- | --- |
| | TM-score | #TM≥0.5 | #TM≥0.8 | TM-score | #TM≥0.5 | #TM≥0.8 | |
| MultiSFold | 0.774 | 78/81 | 37/81 | 0.762 | 79/81 | 36/81 | 70.37 |
| MultiSFold* | 0.756 | 78/81 | 33/81 | 0.748 | 79/81 | 30/81 | 56.79 |
| AlphaFold2 | 0.727 | 78/81 | 28/81 | 0.730 | 78/81 | 27/81 | 9.88 |
| RoseTTAFold | 0.680 | 74/81 | 21/81 | 0.663 | 74/81 | 15/81 | 3.70 |

**Table S6.**
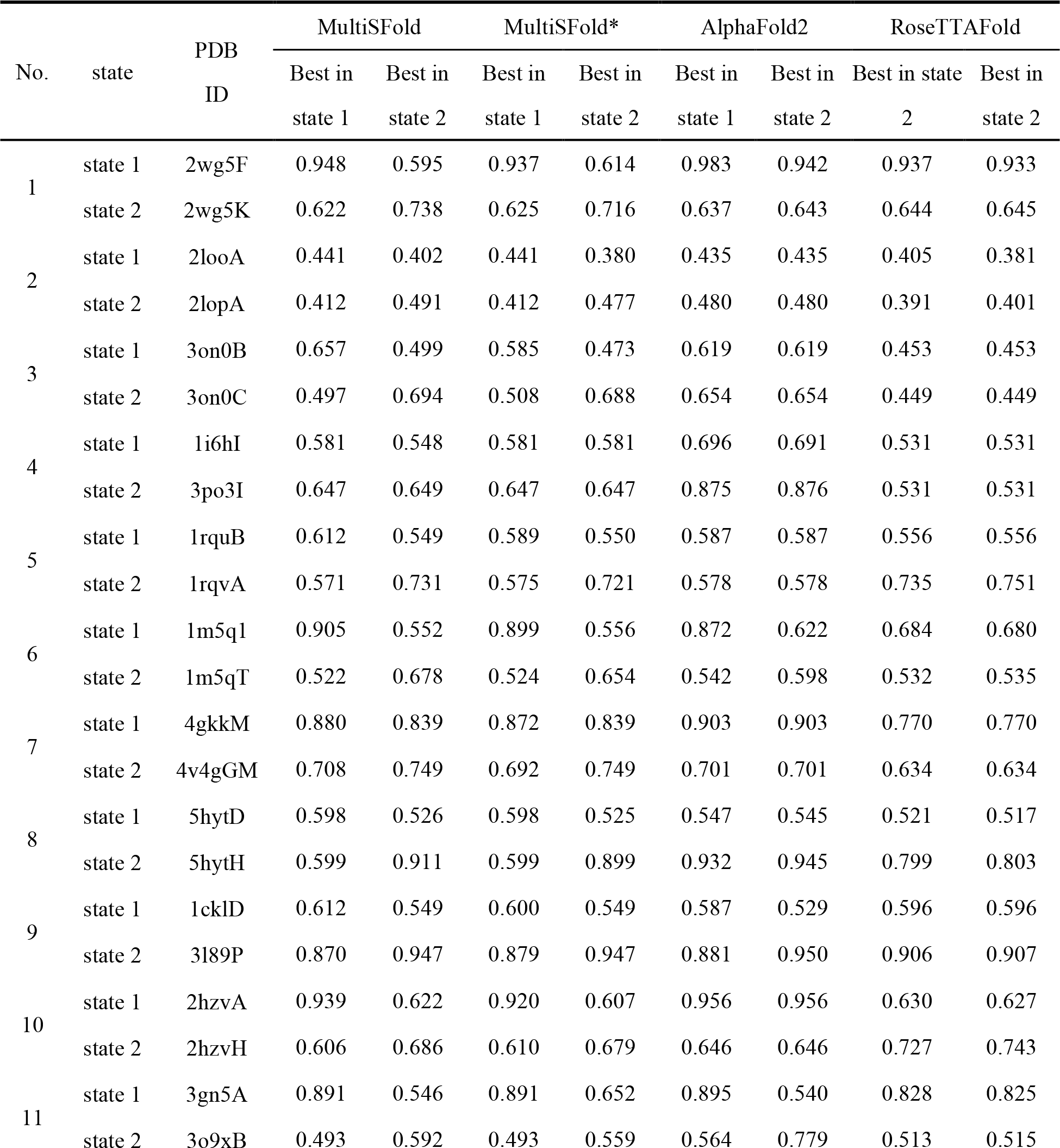
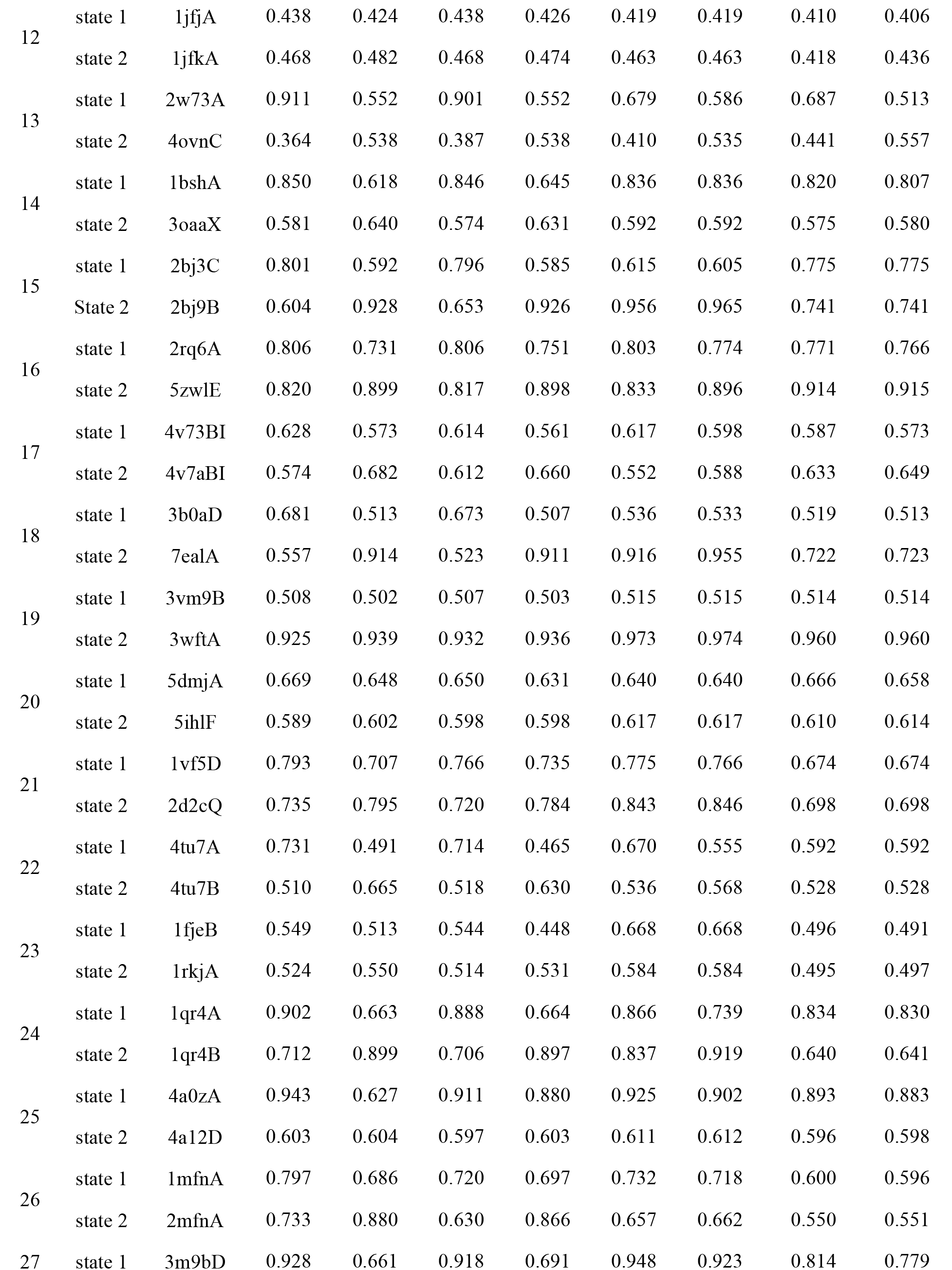
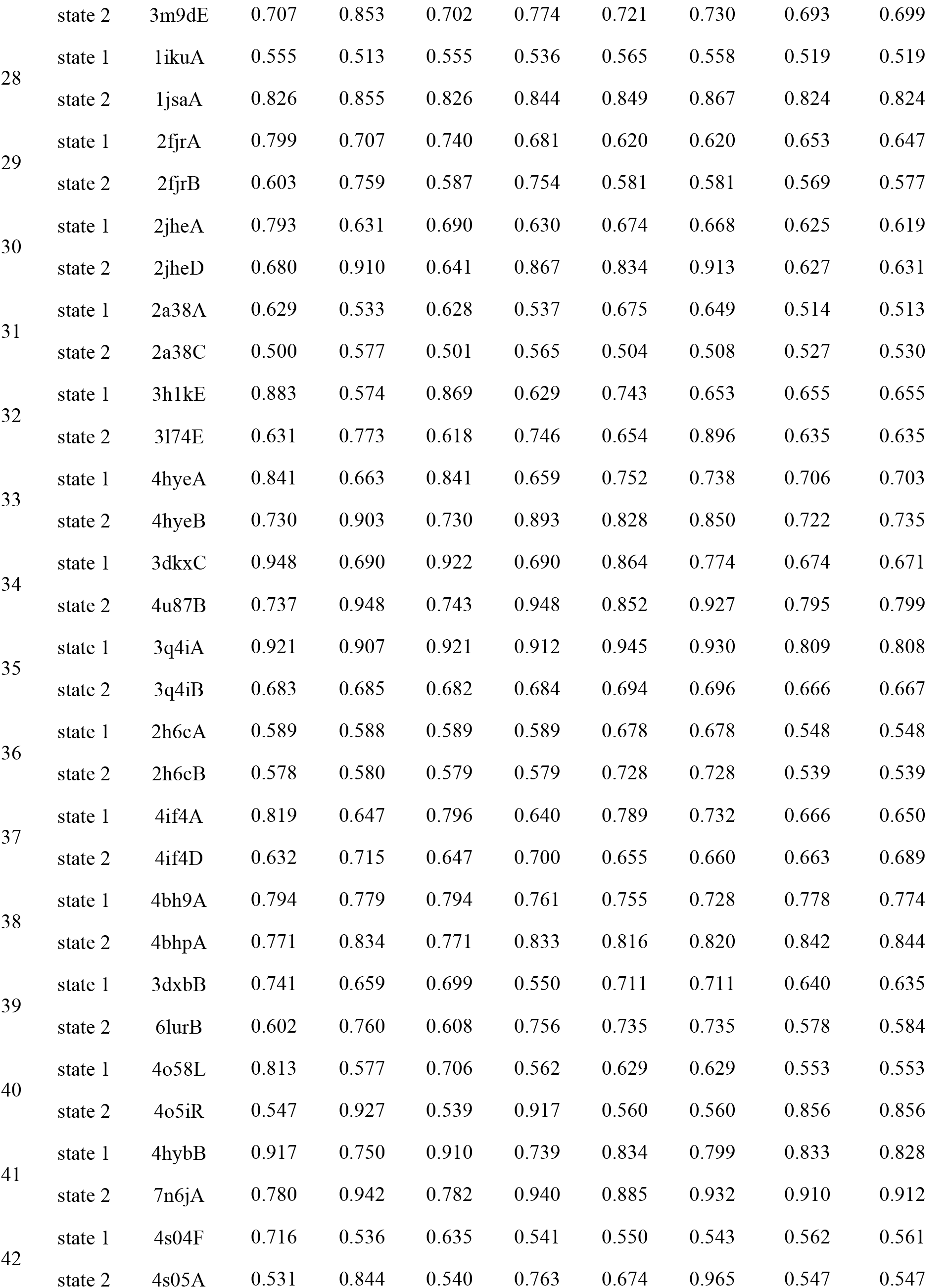
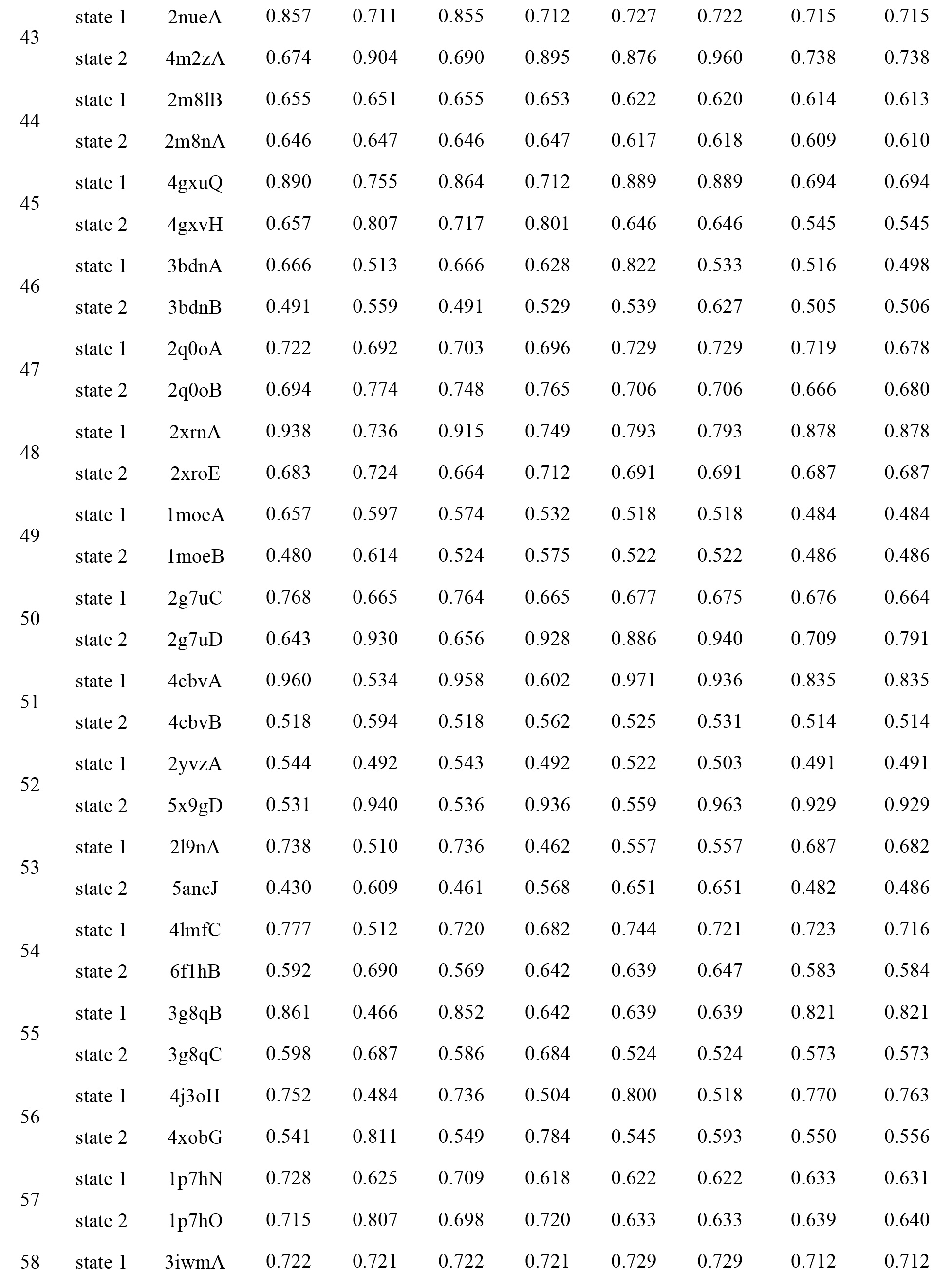
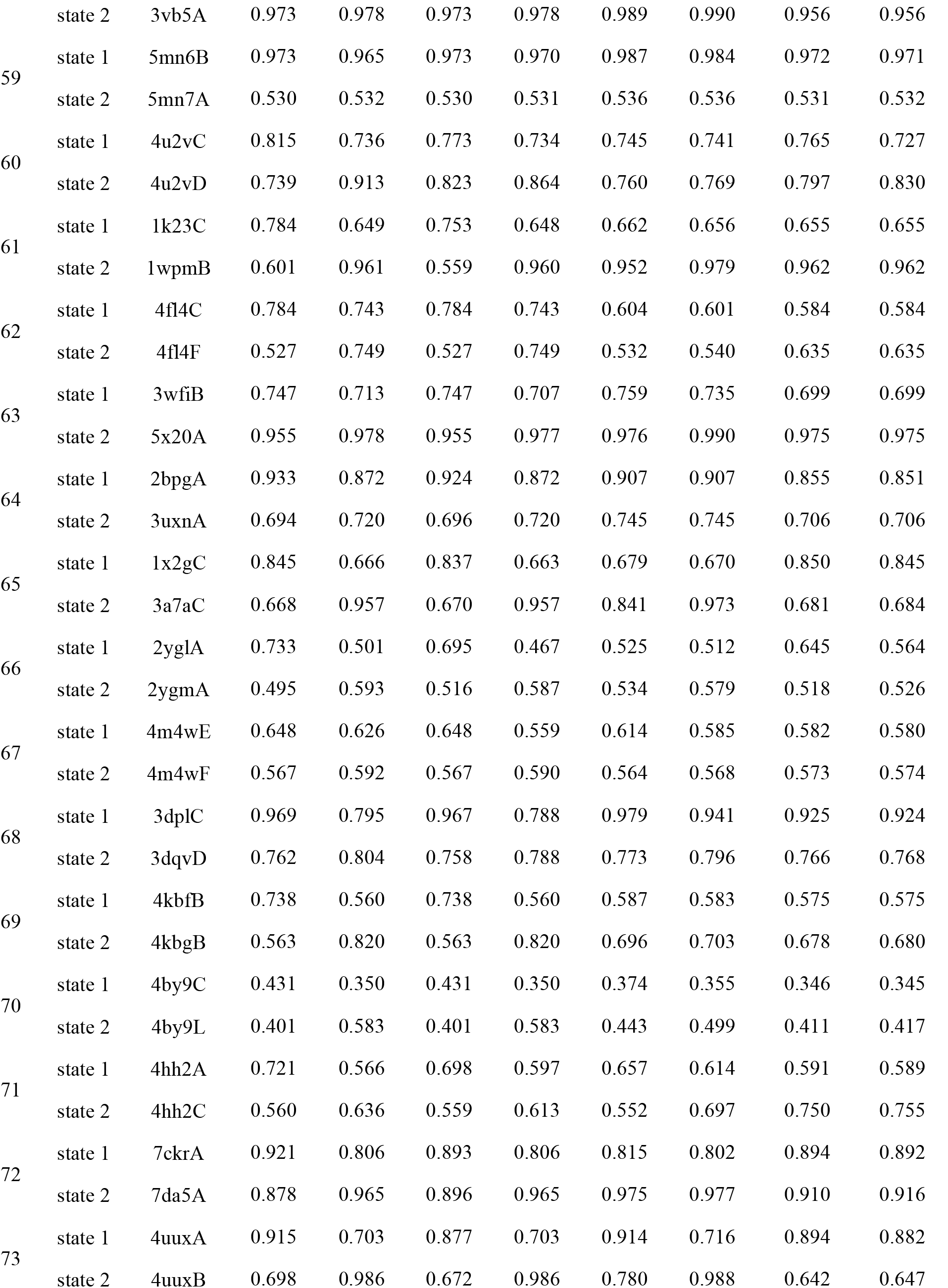
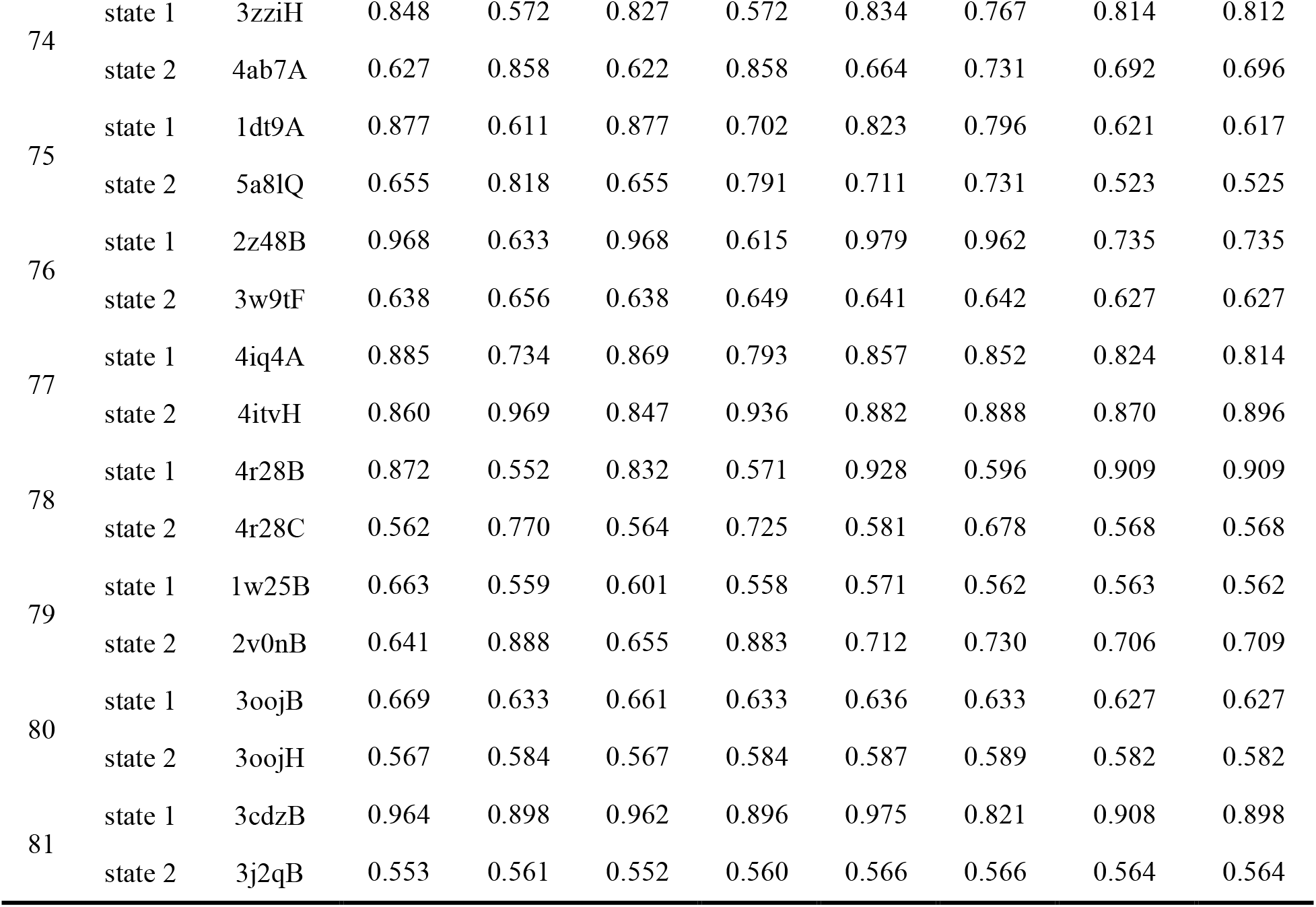
Details of experimental results for predicting multiple conformations.

**Table S7.**
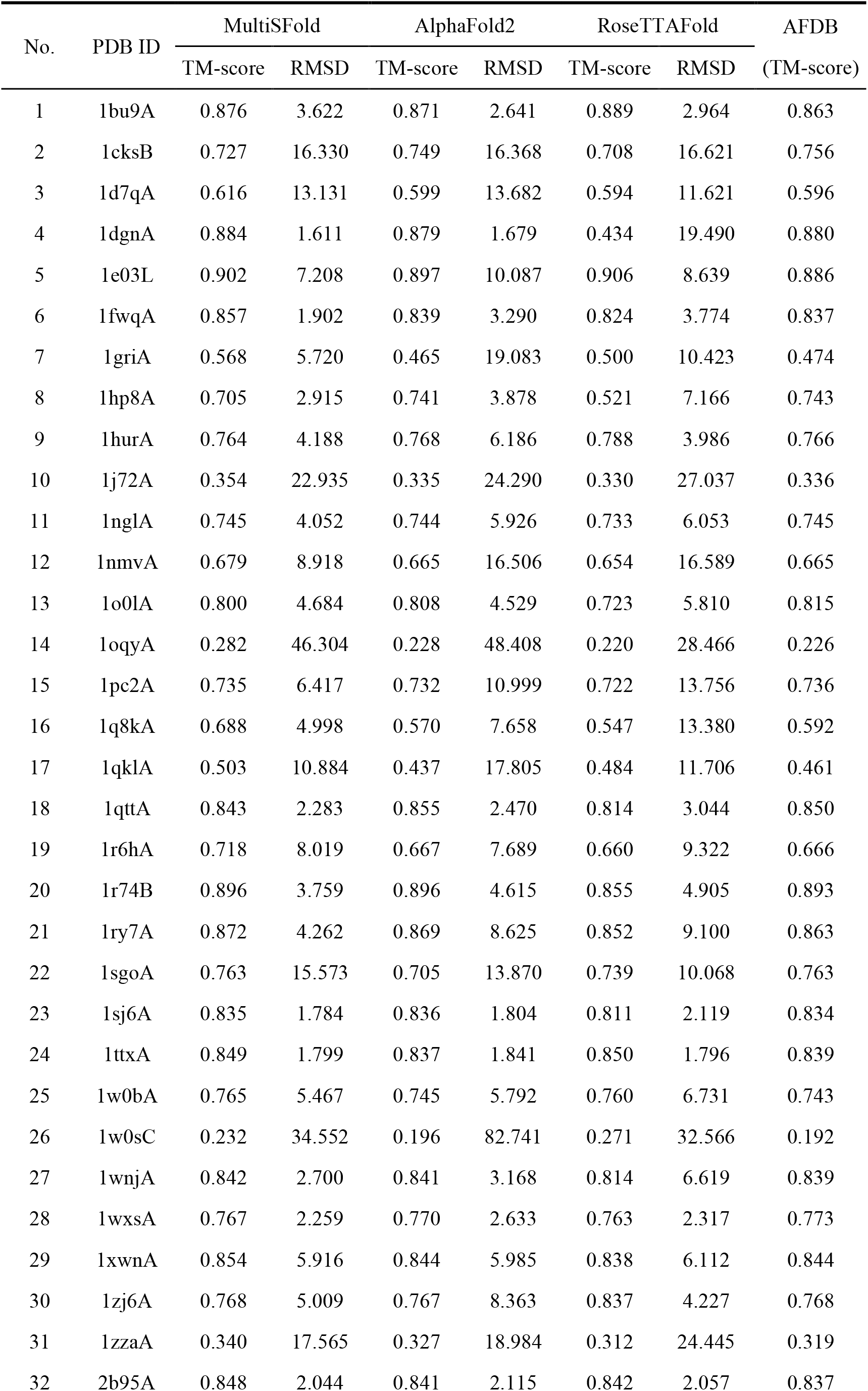
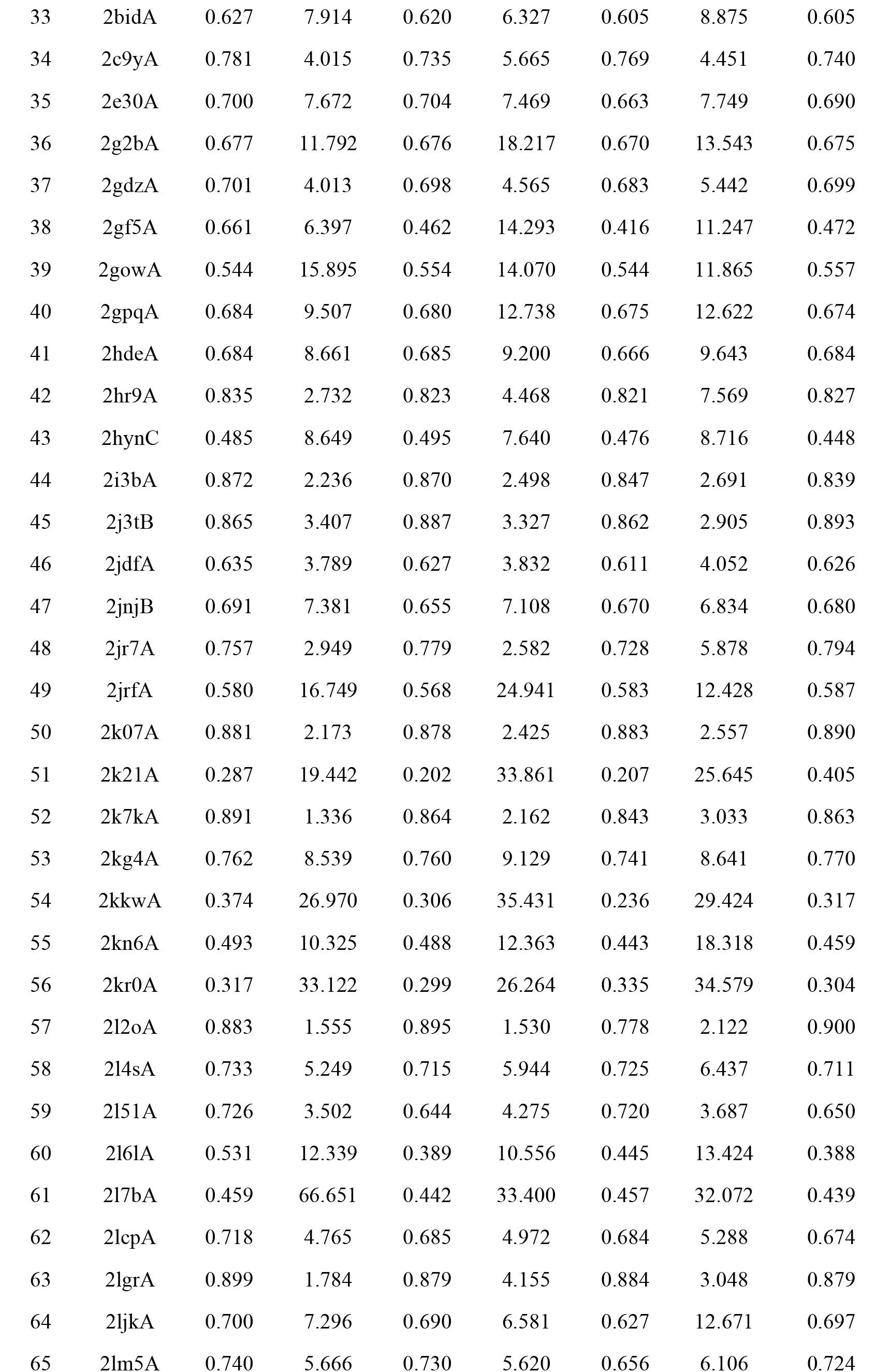
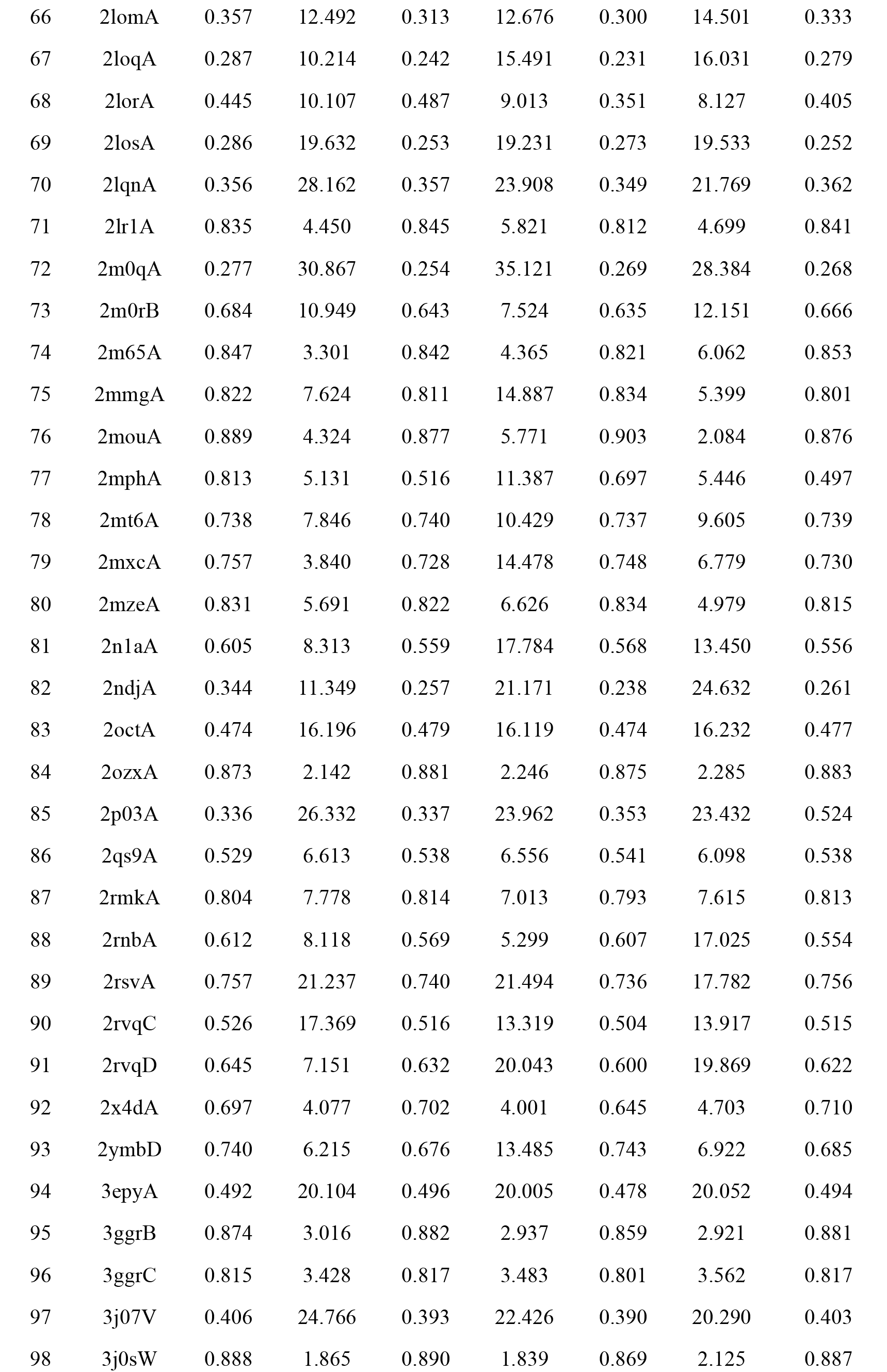
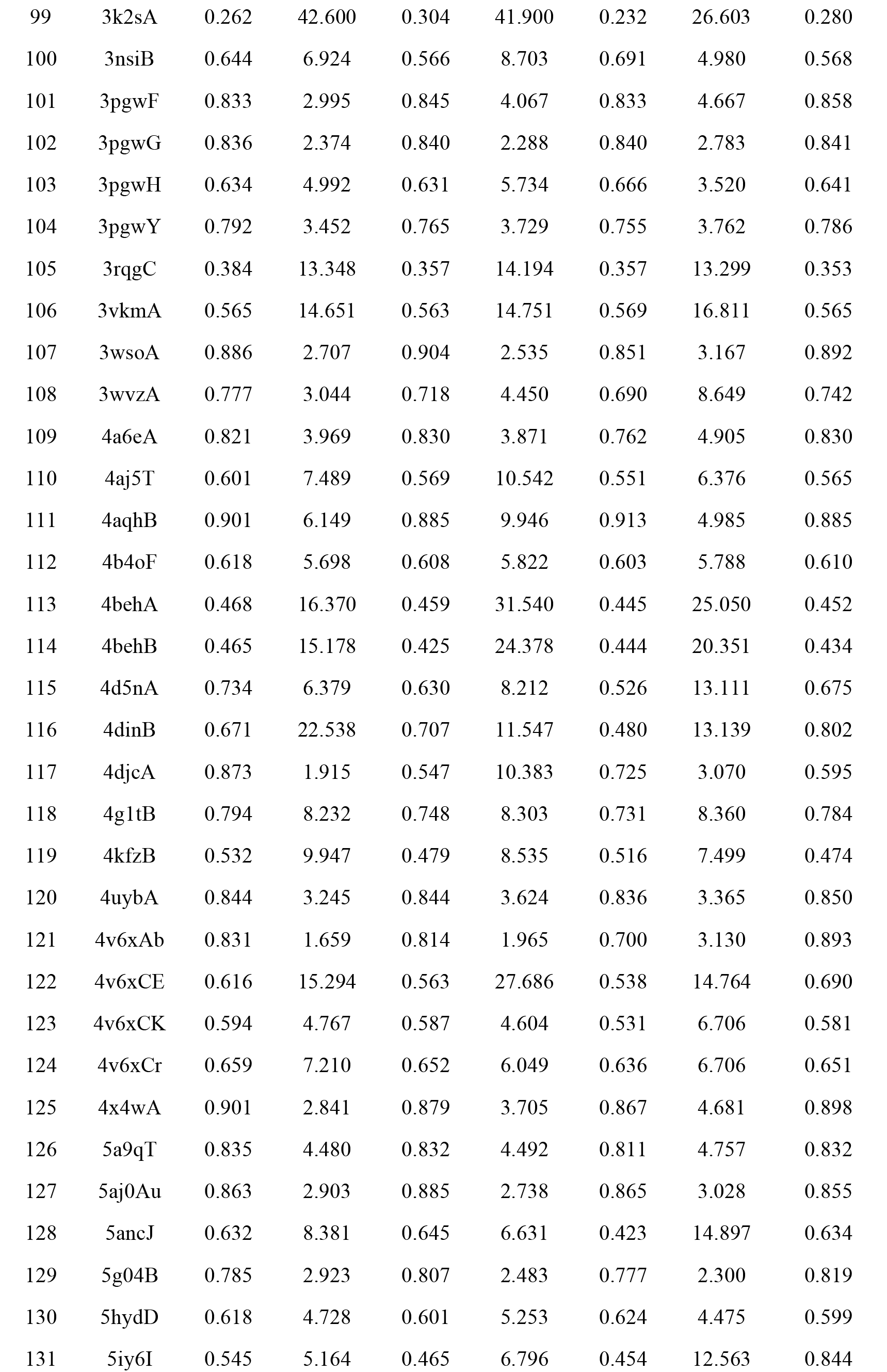
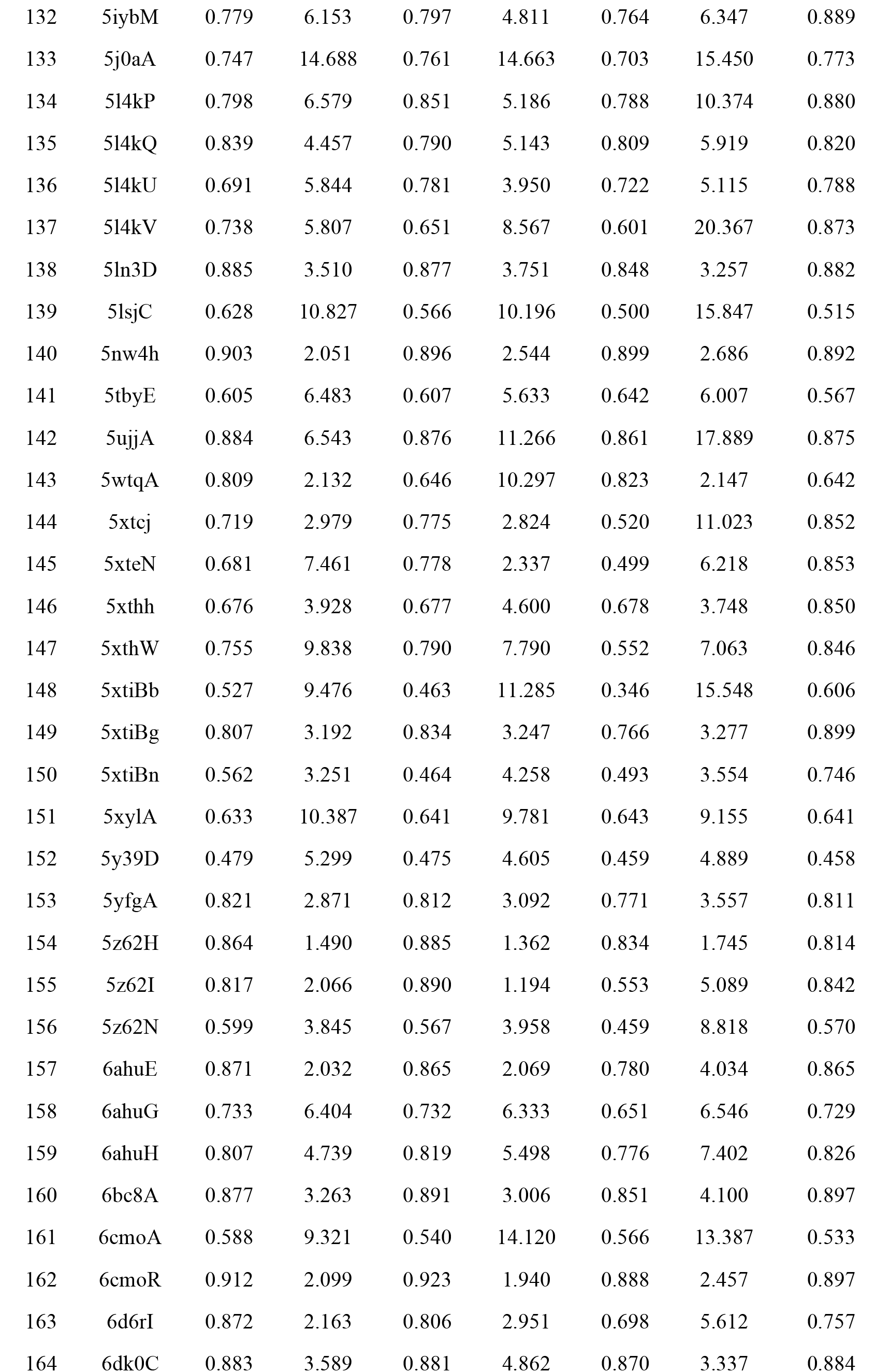
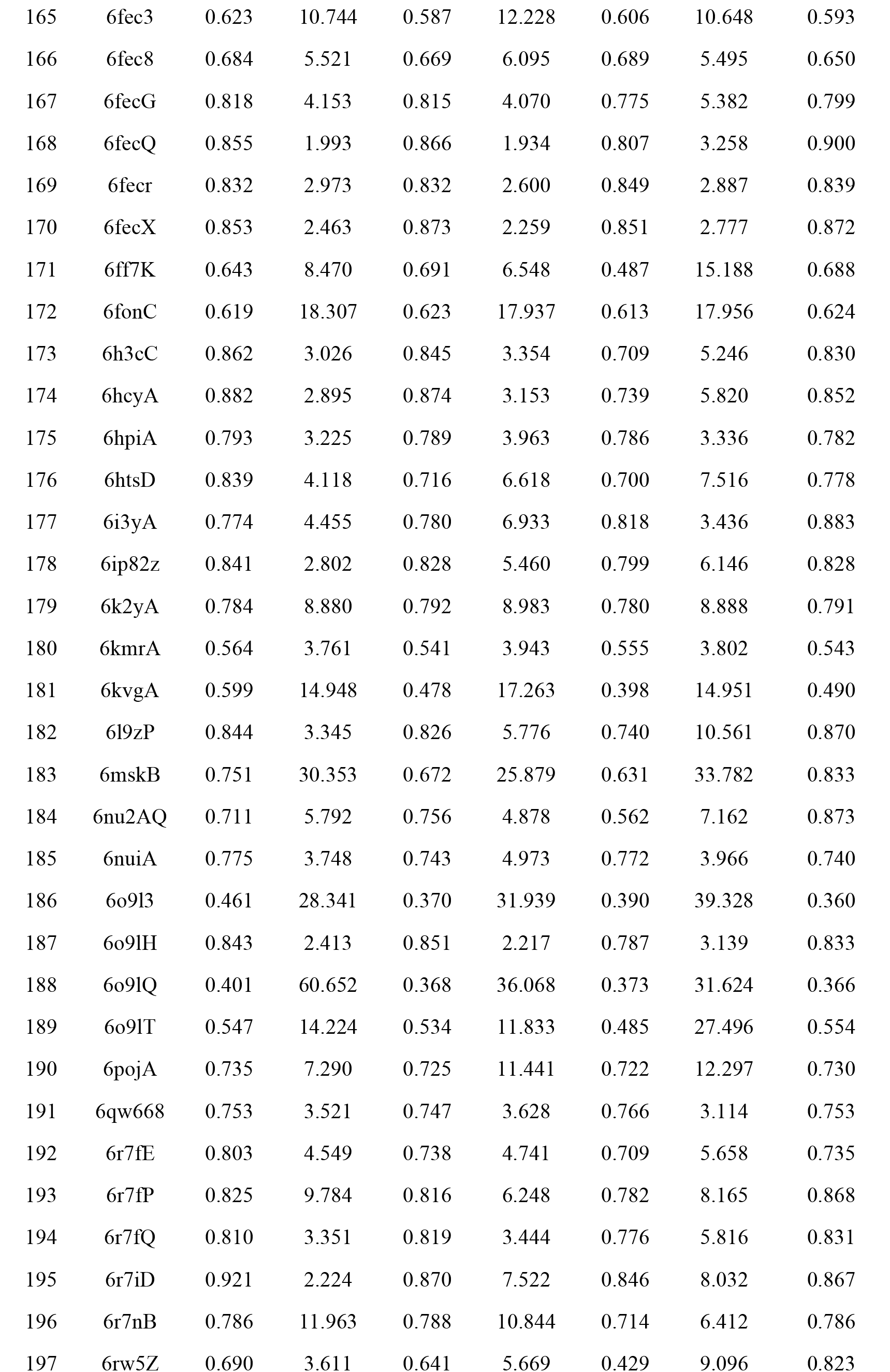
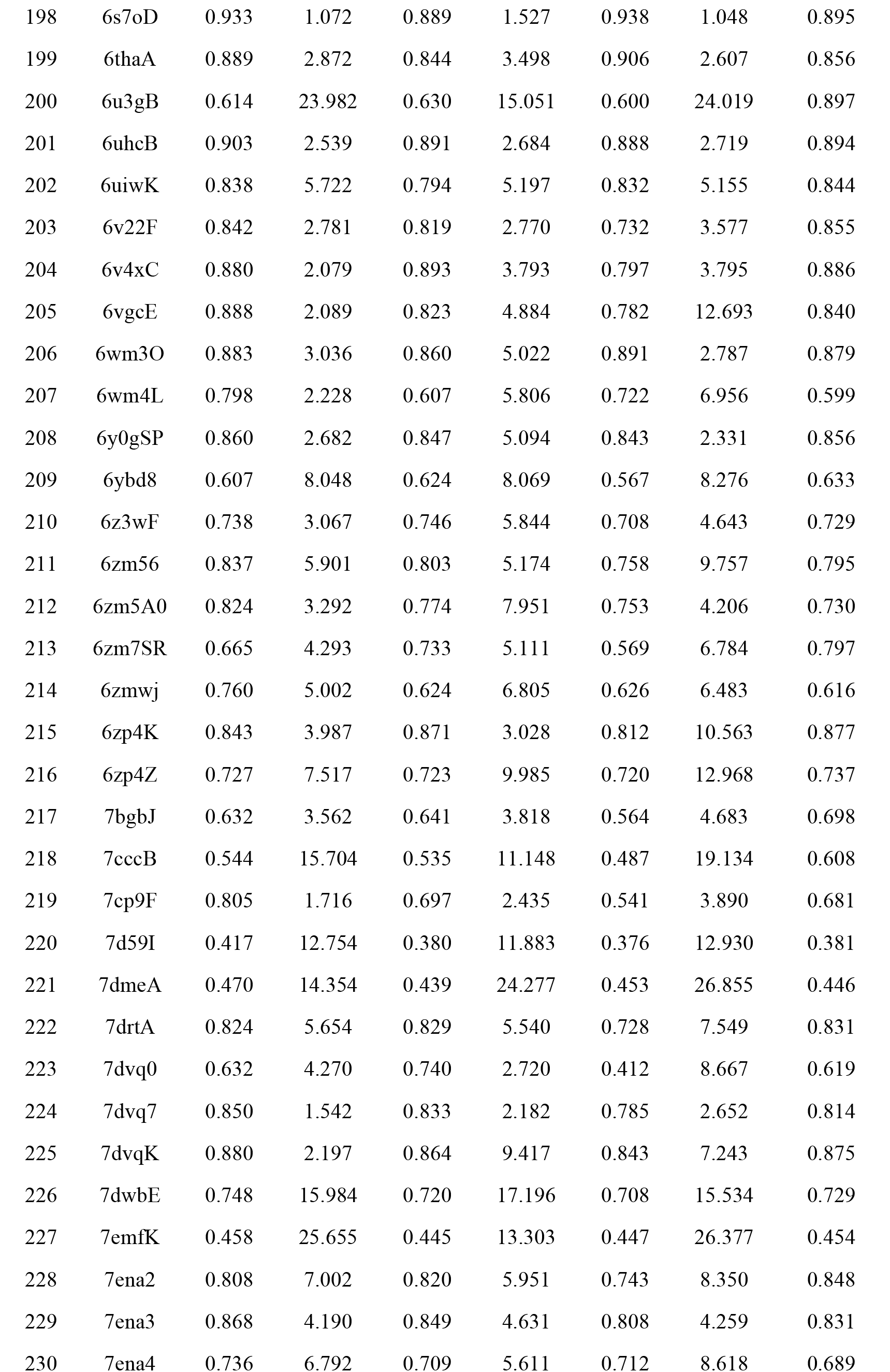
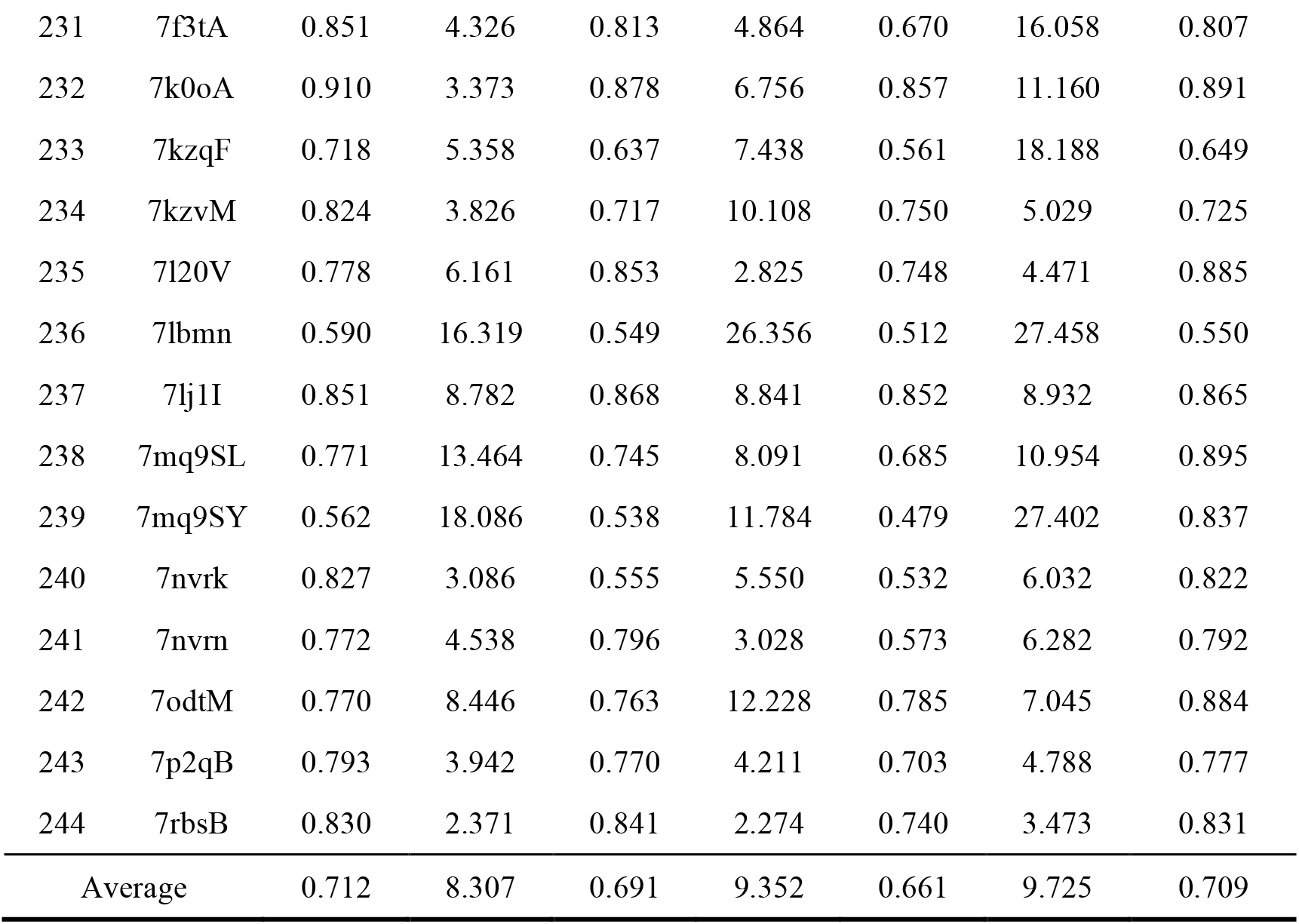
Details of experimental results with state-of-the-art methods on 244 human proteins.

| No. | PDB ID | MultiSFold |  | AlphaFold2 |  | RoseTTAFold |  | AFDB<br>(TM-score) |
| --- | --- | --- | --- | --- | --- | --- | --- | --- |
|  |  | TM-score | RMSD | TM-score | RMSD | TM-score | RMSD |  |
| 1 | 1bu9A | 0.876 | 3.622 | 0.871 | 2.641 | 0.889 | 2.964 | 0.863 |
| 2 | 1cksB | 0.727 | 16.330 | 0.749 | 16.368 | 0.708 | 16.621 | 0.756 |
| 3 | 1d7qA | 0.616 | 13.131 | 0.599 | 13.682 | 0.594 | 11.621 | 0.596 |
| 4 | 1dgnA | 0.884 | 1.611 | 0.879 | 1.679 | 0.434 | 19.490 | 0.880 |
| 5 | 1e03L | 0.902 | 7.208 | 0.897 | 10.087 | 0.906 | 8.639 | 0.886 |
| 6 | 1fwqA | 0.857 | 1.902 | 0.839 | 3.290 | 0.824 | 3.774 | 0.837 |
| 7 | 1griA | 0.568 | 5.720 | 0.465 | 19.083 | 0.500 | 10.423 | 0.474 |
| 8 | 1hp8A | 0.705 | 2.915 | 0.741 | 3.878 | 0.521 | 7.166 | 0.743 |
| 9 | 1hurA | 0.764 | 4.188 | 0.768 | 6.186 | 0.788 | 3.986 | 0.766 |
| 10 | 1j72A | 0.354 | 22.935 | 0.335 | 24.290 | 0.330 | 27.037 | 0.336 |
| 11 | 1nglA | 0.745 | 4.052 | 0.744 | 5.926 | 0.733 | 6.053 | 0.745 |
| 12 | 1nmvA | 0.679 | 8.918 | 0.665 | 16.506 | 0.654 | 16.589 | 0.665 |
| 13 | 1o0lA | 0.800 | 4.684 | 0.808 | 4.529 | 0.723 | 5.810 | 0.815 |
| 14 | 1oqyA | 0.282 | 46.304 | 0.228 | 48.408 | 0.220 | 28.466 | 0.226 |
| 15 | 1pc2A | 0.735 | 6.417 | 0.732 | 10.999 | 0.722 | 13.756 | 0.736 |
| 16 | 1q8kA | 0.688 | 4.998 | 0.570 | 7.658 | 0.547 | 13.380 | 0.592 |
| 17 | 1qklA | 0.503 | 10.884 | 0.437 | 17.805 | 0.484 | 11.706 | 0.461 |
| 18 | 1qttA | 0.843 | 2.283 | 0.855 | 2.470 | 0.814 | 3.044 | 0.850 |
| 19 | 1r6hA | 0.718 | 8.019 | 0.667 | 7.689 | 0.660 | 9.322 | 0.666 |
| 20 | 1r74B | 0.896 | 3.759 | 0.896 | 4.615 | 0.855 | 4.905 | 0.893 |
| 21 | 1ry7A | 0.872 | 4.262 | 0.869 | 8.625 | 0.852 | 9.100 | 0.863 |
| 22 | 1sgoA | 0.763 | 15.573 | 0.705 | 13.870 | 0.739 | 10.068 | 0.763 |
| 23 | 1sj6A | 0.835 | 1.784 | 0.836 | 1.804 | 0.811 | 2.119 | 0.834 |
| 24 | 1ttxA | 0.849 | 1.799 | 0.837 | 1.841 | 0.850 | 1.796 | 0.839 |
| 25 | 1w0bA | 0.765 | 5.467 | 0.745 | 5.792 | 0.760 | 6.731 | 0.743 |
| 26 | 1w0sC | 0.232 | 34.552 | 0.196 | 82.741 | 0.271 | 32.566 | 0.192 |
| 27 | 1wnjA | 0.842 | 2.700 | 0.841 | 3.168 | 0.814 | 6.619 | 0.839 |
| 28 | 1wxsA | 0.767 | 2.259 | 0.770 | 2.633 | 0.763 | 2.317 | 0.773 |
| 29 | 1xwnA | 0.854 | 5.916 | 0.844 | 5.985 | 0.838 | 6.112 | 0.844 |
| 30 | 1zj6A | 0.768 | 5.009 | 0.767 | 8.363 | 0.837 | 4.227 | 0.768 |
| 31 | 1zzaA | 0.340 | 17.565 | 0.327 | 18.984 | 0.312 | 24.445 | 0.319 |
| 32 | 2b95A | 0.848 | 2.044 | 0.841 | 2.115 | 0.842 | 2.057 | 0.837 |
|  |  | TM-score | RMSD | TM-score | RMSD | TM-score | RMSD |  |
| 33 | 2bidA | 0.627 | 7.914 | 0.620 | 6.327 | 0.605 | 8.875 | 0.605 |
| 34 | 2c9yA | 0.781 | 4.015 | 0.735 | 5.665 | 0.769 | 4.451 | 0.740 |
| 35 | 2e30A | 0.700 | 7.672 | 0.704 | 7.469 | 0.663 | 7.749 | 0.690 |
| 36 | 2g2bA | 0.677 | 11.792 | 0.676 | 18.217 | 0.670 | 13.543 | 0.675 |
| 37 | 2gdzA | 0.701 | 4.013 | 0.698 | 4.565 | 0.683 | 5.442 | 0.699 |
| 38 | 2gf5A | 0.661 | 6.397 | 0.462 | 14.293 | 0.416 | 11.247 | 0.472 |
| 39 | 2gowA | 0.544 | 15.895 | 0.554 | 14.070 | 0.544 | 11.865 | 0.557 |
| 40 | 2gpqA | 0.684 | 9.507 | 0.680 | 12.738 | 0.675 | 12.622 | 0.674 |
| 41 | 2hdeA | 0.684 | 8.661 | 0.685 | 9.200 | 0.666 | 9.643 | 0.684 |
| 42 | 2hr9A | 0.835 | 2.732 | 0.823 | 4.468 | 0.821 | 7.569 | 0.827 |
| 43 | 2hynC | 0.485 | 8.649 | 0.495 | 7.640 | 0.476 | 8.716 | 0.448 |
| 44 | 2i3bA | 0.872 | 2.236 | 0.870 | 2.498 | 0.847 | 2.691 | 0.839 |
| 45 | 2j3tB | 0.865 | 3.407 | 0.887 | 3.327 | 0.862 | 2.905 | 0.893 |
| 46 | 2jdfA | 0.635 | 3.789 | 0.627 | 3.832 | 0.611 | 4.052 | 0.626 |
| 47 | 2jnjB | 0.691 | 7.381 | 0.655 | 7.108 | 0.670 | 6.834 | 0.680 |
| 48 | 2jr7A | 0.757 | 2.949 | 0.779 | 2.582 | 0.728 | 5.878 | 0.794 |
| 49 | 2jrfA | 0.580 | 16.749 | 0.568 | 24.941 | 0.583 | 12.428 | 0.587 |
| 50 | 2k07A | 0.881 | 2.173 | 0.878 | 2.425 | 0.883 | 2.557 | 0.890 |
| 51 | 2k21A | 0.287 | 19.442 | 0.202 | 33.861 | 0.207 | 25.645 | 0.405 |
| 52 | 2k7kA | 0.891 | 1.336 | 0.864 | 2.162 | 0.843 | 3.033 | 0.863 |
| 53 | 2kg4A | 0.762 | 8.539 | 0.760 | 9.129 | 0.741 | 8.641 | 0.770 |
| 54 | 2kkwA | 0.374 | 26.970 | 0.306 | 35.431 | 0.236 | 29.424 | 0.317 |
| 55 | 2kn6A | 0.493 | 10.325 | 0.488 | 12.363 | 0.443 | 18.318 | 0.459 |
| 56 | 2kr0A | 0.317 | 33.122 | 0.299 | 26.264 | 0.335 | 34.579 | 0.304 |
| 57 | 2l2oA | 0.883 | 1.555 | 0.895 | 1.530 | 0.778 | 2.122 | 0.900 |
| 58 | 2l4sA | 0.733 | 5.249 | 0.715 | 5.944 | 0.725 | 6.437 | 0.711 |
| 59 | 2l51A | 0.726 | 3.502 | 0.644 | 4.275 | 0.720 | 3.687 | 0.650 |
| 60 | 2l6lA | 0.531 | 12.339 | 0.389 | 10.556 | 0.445 | 13.424 | 0.388 |
| 61 | 2l7bA | 0.459 | 66.651 | 0.442 | 33.400 | 0.457 | 32.072 | 0.439 |
| 62 | 2lcpA | 0.718 | 4.765 | 0.685 | 4.972 | 0.684 | 5.288 | 0.674 |
| 63 | 2lgrA | 0.899 | 1.784 | 0.879 | 4.155 | 0.884 | 3.048 | 0.879 |
| 64 | 2ljkA | 0.700 | 7.296 | 0.690 | 6.581 | 0.627 | 12.671 | 0.697 |
| 65 | 2lm5A | 0.740 | 5.666 | 0.730 | 5.620 | 0.656 | 6.106 | 0.724 |
|  |  | TM-score | RMSD | TM-score | RMSD | TM-score | RMSD |  |
| 66 | 2lomA | 0.357 | 12.492 | 0.313 | 12.676 | 0.300 | 14.501 | 0.333 |
| 67 | 2loqA | 0.287 | 10.214 | 0.242 | 15.491 | 0.231 | 16.031 | 0.279 |
| 68 | 2lorA | 0.445 | 10.107 | 0.487 | 9.013 | 0.351 | 8.127 | 0.405 |
| 69 | 2losA | 0.286 | 19.632 | 0.253 | 19.231 | 0.273 | 19.533 | 0.252 |
| 70 | 2lqnA | 0.356 | 28.162 | 0.357 | 23.908 | 0.349 | 21.769 | 0.362 |
| 71 | 2lr1A | 0.835 | 4.450 | 0.845 | 5.821 | 0.812 | 4.699 | 0.841 |
| 72 | 2m0qA | 0.277 | 30.867 | 0.254 | 35.121 | 0.269 | 28.384 | 0.268 |
| 73 | 2m0rB | 0.684 | 10.949 | 0.643 | 7.524 | 0.635 | 12.151 | 0.666 |
| 74 | 2m65A | 0.847 | 3.301 | 0.842 | 4.365 | 0.821 | 6.062 | 0.853 |
| 75 | 2mmgA | 0.822 | 7.624 | 0.811 | 14.887 | 0.834 | 5.399 | 0.801 |
| 76 | 2mouA | 0.889 | 4.324 | 0.877 | 5.771 | 0.903 | 2.084 | 0.876 |
| 77 | 2mphA | 0.813 | 5.131 | 0.516 | 11.387 | 0.697 | 5.446 | 0.497 |
| 78 | 2mt6A | 0.738 | 7.846 | 0.740 | 10.429 | 0.737 | 9.605 | 0.739 |
| 79 | 2mxcA | 0.757 | 3.840 | 0.728 | 14.478 | 0.748 | 6.779 | 0.730 |
| 80 | 2mzeA | 0.831 | 5.691 | 0.822 | 6.626 | 0.834 | 4.979 | 0.815 |
| 81 | 2n1aA | 0.605 | 8.313 | 0.559 | 17.784 | 0.568 | 13.450 | 0.556 |
| 82 | 2ndjA | 0.344 | 11.349 | 0.257 | 21.171 | 0.238 | 24.632 | 0.261 |
| 83 | 2octA | 0.474 | 16.196 | 0.479 | 16.119 | 0.474 | 16.232 | 0.477 |
| 84 | 2ozxA | 0.873 | 2.142 | 0.881 | 2.246 | 0.875 | 2.285 | 0.883 |
| 85 | 2p03A | 0.336 | 26.332 | 0.337 | 23.962 | 0.353 | 23.432 | 0.524 |
| 86 | 2qs9A | 0.529 | 6.613 | 0.538 | 6.556 | 0.541 | 6.098 | 0.538 |
| 87 | 2rmkA | 0.804 | 7.778 | 0.814 | 7.013 | 0.793 | 7.615 | 0.813 |
| 88 | 2rnbA | 0.612 | 8.118 | 0.569 | 5.299 | 0.607 | 17.025 | 0.554 |
| 89 | 2rsvA | 0.757 | 21.237 | 0.740 | 21.494 | 0.736 | 17.782 | 0.756 |
| 90 | 2rvqC | 0.526 | 17.369 | 0.516 | 13.319 | 0.504 | 13.917 | 0.515 |
| 91 | 2rvqD | 0.645 | 7.151 | 0.632 | 20.043 | 0.600 | 19.869 | 0.622 |
| 92 | 2x4dA | 0.697 | 4.077 | 0.702 | 4.001 | 0.645 | 4.703 | 0.710 |
| 93 | 2ymbD | 0.740 | 6.215 | 0.676 | 13.485 | 0.743 | 6.922 | 0.685 |
| 94 | 3epyA | 0.492 | 20.104 | 0.496 | 20.005 | 0.478 | 20.052 | 0.494 |
| 95 | 3ggrB | 0.874 | 3.016 | 0.882 | 2.937 | 0.859 | 2.921 | 0.881 |
| 96 | 3ggrC | 0.815 | 3.428 | 0.817 | 3.483 | 0.801 | 3.562 | 0.817 |
| 97 | 3j07V | 0.406 | 24.766 | 0.393 | 22.426 | 0.390 | 20.290 | 0.403 |
| 98 | 3j0sW | 0.888 | 1.865 | 0.890 | 1.839 | 0.869 | 2.125 | 0.887 |
|  |  | TM-score | RMSD | TM-score | RMSD | TM-score | RMSD |  |
| 99 | 3k2sA | 0.262 | 42.600 | 0.304 | 41.900 | 0.232 | 26.603 | 0.280 |
| 100 | 3nsiB | 0.644 | 6.924 | 0.566 | 8.703 | 0.691 | 4.980 | 0.568 |
| 101 | 3pgwF | 0.833 | 2.995 | 0.845 | 4.067 | 0.833 | 4.667 | 0.858 |
| 102 | 3pgwG | 0.836 | 2.374 | 0.840 | 2.288 | 0.840 | 2.783 | 0.841 |
| 103 | 3pgwH | 0.634 | 4.992 | 0.631 | 5.734 | 0.666 | 3.520 | 0.641 |
| 104 | 3pgwY | 0.792 | 3.452 | 0.765 | 3.729 | 0.755 | 3.762 | 0.786 |
| 105 | 3rqgC | 0.384 | 13.348 | 0.357 | 14.194 | 0.357 | 13.299 | 0.353 |
| 106 | 3vkmA | 0.565 | 14.651 | 0.563 | 14.751 | 0.569 | 16.811 | 0.565 |
| 107 | 3wsoA | 0.886 | 2.707 | 0.904 | 2.535 | 0.851 | 3.167 | 0.892 |
| 108 | 3wvzA | 0.777 | 3.044 | 0.718 | 4.450 | 0.690 | 8.649 | 0.742 |
| 109 | 4a6eA | 0.821 | 3.969 | 0.830 | 3.871 | 0.762 | 4.905 | 0.830 |
| 110 | 4aj5T | 0.601 | 7.489 | 0.569 | 10.542 | 0.551 | 6.376 | 0.565 |
| 111 | 4aqhB | 0.901 | 6.149 | 0.885 | 9.946 | 0.913 | 4.985 | 0.885 |
| 112 | 4b4oF | 0.618 | 5.698 | 0.608 | 5.822 | 0.603 | 5.788 | 0.610 |
| 113 | 4behA | 0.468 | 16.370 | 0.459 | 31.540 | 0.445 | 25.050 | 0.452 |
| 114 | 4behB | 0.465 | 15.178 | 0.425 | 24.378 | 0.444 | 20.351 | 0.434 |
| 115 | 4d5nA | 0.734 | 6.379 | 0.630 | 8.212 | 0.526 | 13.111 | 0.675 |
| 116 | 4dinB | 0.671 | 22.538 | 0.707 | 11.547 | 0.480 | 13.139 | 0.802 |
| 117 | 4djcA | 0.873 | 1.915 | 0.547 | 10.383 | 0.725 | 3.070 | 0.595 |
| 118 | 4g1tB | 0.794 | 8.232 | 0.748 | 8.303 | 0.731 | 8.360 | 0.784 |
| 119 | 4kfzB | 0.532 | 9.947 | 0.479 | 8.535 | 0.516 | 7.499 | 0.474 |
| 120 | 4uybA | 0.844 | 3.245 | 0.844 | 3.624 | 0.836 | 3.365 | 0.850 |
| 121 | 4v6xAb | 0.831 | 1.659 | 0.814 | 1.965 | 0.700 | 3.130 | 0.893 |
| 122 | 4v6xCE | 0.616 | 15.294 | 0.563 | 27.686 | 0.538 | 14.764 | 0.690 |
| 123 | 4v6xCK | 0.594 | 4.767 | 0.587 | 4.604 | 0.531 | 6.706 | 0.581 |
| 124 | 4v6xCr | 0.659 | 7.210 | 0.652 | 6.049 | 0.636 | 6.706 | 0.651 |
| 125 | 4x4wA | 0.901 | 2.841 | 0.879 | 3.705 | 0.867 | 4.681 | 0.898 |
| 126 | 5a9qT | 0.835 | 4.480 | 0.832 | 4.492 | 0.811 | 4.757 | 0.832 |
| 127 | 5aj0Au | 0.863 | 2.903 | 0.885 | 2.738 | 0.865 | 3.028 | 0.855 |
| 128 | 5ancJ | 0.632 | 8.381 | 0.645 | 6.631 | 0.423 | 14.897 | 0.634 |
| 129 | 5g04B | 0.785 | 2.923 | 0.807 | 2.483 | 0.777 | 2.300 | 0.819 |
| 130 | 5hydD | 0.618 | 4.728 | 0.601 | 5.253 | 0.624 | 4.475 | 0.599 |
| 131 | 5iy6I | 0.545 | 5.164 | 0.465 | 6.796 | 0.454 | 12.563 | 0.844 |

| No. | PDB ID | MultiSFold |  | AlphaFold2 |  | RoseTTAFold |  | AFDB |
| --- | --- | --- | --- | --- | --- | --- | --- | --- |
|  |  | TM-score | RMSD | TM-score | RMSD | TM-score | RMSD | (TM-score) |
| 132 | 5iybM | 0.779 | 6.153 | 0.797 | 4.811 | 0.764 | 6.347 | 0.889 |
| 133 | 5j0aA | 0.747 | 14.688 | 0.761 | 14.663 | 0.703 | 15.450 | 0.773 |
| 134 | 5l4kP | 0.798 | 6.579 | 0.851 | 5.186 | 0.788 | 10.374 | 0.880 |
| 135 | 5l4kQ | 0.839 | 4.457 | 0.790 | 5.143 | 0.809 | 5.919 | 0.820 |
| 136 | 5l4kU | 0.691 | 5.844 | 0.781 | 3.950 | 0.722 | 5.115 | 0.788 |
| 137 | 5l4kV | 0.738 | 5.807 | 0.651 | 8.567 | 0.601 | 20.367 | 0.873 |
| 138 | 5ln3D | 0.885 | 3.510 | 0.877 | 3.751 | 0.848 | 3.257 | 0.882 |
| 139 | 5lsjC | 0.628 | 10.827 | 0.566 | 10.196 | 0.500 | 15.847 | 0.515 |
| 140 | 5nw4h | 0.903 | 2.051 | 0.896 | 2.544 | 0.899 | 2.686 | 0.892 |
| 141 | 5tbyE | 0.605 | 6.483 | 0.607 | 5.633 | 0.642 | 6.007 | 0.567 |
| 142 | 5ujjA | 0.884 | 6.543 | 0.876 | 11.266 | 0.861 | 17.889 | 0.875 |
| 143 | 5wtqA | 0.809 | 2.132 | 0.646 | 10.297 | 0.823 | 2.147 | 0.642 |
| 144 | 5xtcj | 0.719 | 2.979 | 0.775 | 2.824 | 0.520 | 11.023 | 0.852 |
| 145 | 5xteN | 0.681 | 7.461 | 0.778 | 2.337 | 0.499 | 6.218 | 0.853 |
| 146 | 5xthh | 0.676 | 3.928 | 0.677 | 4.600 | 0.678 | 3.748 | 0.850 |
| 147 | 5xthW | 0.755 | 9.838 | 0.790 | 7.790 | 0.552 | 7.063 | 0.846 |
| 148 | 5xtiBb | 0.527 | 9.476 | 0.463 | 11.285 | 0.346 | 15.548 | 0.606 |
| 149 | 5xtiBg | 0.807 | 3.192 | 0.834 | 3.247 | 0.766 | 3.277 | 0.899 |
| 150 | 5xtiBn | 0.562 | 3.251 | 0.464 | 4.258 | 0.493 | 3.554 | 0.746 |
| 151 | 5xylA | 0.633 | 10.387 | 0.641 | 9.781 | 0.643 | 9.155 | 0.641 |
| 152 | 5y39D | 0.479 | 5.299 | 0.475 | 4.605 | 0.459 | 4.889 | 0.458 |
| 153 | 5yfgA | 0.821 | 2.871 | 0.812 | 3.092 | 0.771 | 3.557 | 0.811 |
| 154 | 5z62H | 0.864 | 1.490 | 0.885 | 1.362 | 0.834 | 1.745 | 0.814 |
| 155 | 5z62I | 0.817 | 2.066 | 0.890 | 1.194 | 0.553 | 5.089 | 0.842 |
| 156 | 5z62N | 0.599 | 3.845 | 0.567 | 3.958 | 0.459 | 8.818 | 0.570 |
| 157 | 6ahuE | 0.871 | 2.032 | 0.865 | 2.069 | 0.780 | 4.034 | 0.865 |
| 158 | 6ahuG | 0.733 | 6.404 | 0.732 | 6.333 | 0.651 | 6.546 | 0.729 |
| 159 | 6ahuH | 0.807 | 4.739 | 0.819 | 5.498 | 0.776 | 7.402 | 0.826 |
| 160 | 6bc8A | 0.877 | 3.263 | 0.891 | 3.006 | 0.851 | 4.100 | 0.897 |
| 161 | 6cmoA | 0.588 | 9.321 | 0.540 | 14.120 | 0.566 | 13.387 | 0.533 |
| 162 | 6cmoR | 0.912 | 2.099 | 0.923 | 1.940 | 0.888 | 2.457 | 0.897 |
| 163 | 6d6rI | 0.872 | 2.163 | 0.806 | 2.951 | 0.698 | 5.612 | 0.757 |
| 164 | 6dk0C | 0.883 | 3.589 | 0.881 | 4.862 | 0.870 | 3.337 | 0.884 |
|  |  | TM-score | RMSD | TM-score | RMSD | TM-score | RMSD |  |
| 165 | 6fec3 | 0.623 | 10.744 | 0.587 | 12.228 | 0.606 | 10.648 | 0.593 |
| 166 | 6fec8 | 0.684 | 5.521 | 0.669 | 6.095 | 0.689 | 5.495 | 0.650 |
| 167 | 6fecG | 0.818 | 4.153 | 0.815 | 4.070 | 0.775 | 5.382 | 0.799 |
| 168 | 6fecQ | 0.855 | 1.993 | 0.866 | 1.934 | 0.807 | 3.258 | 0.900 |
| 169 | 6fecr | 0.832 | 2.973 | 0.832 | 2.600 | 0.849 | 2.887 | 0.839 |
| 170 | 6fecX | 0.853 | 2.463 | 0.873 | 2.259 | 0.851 | 2.777 | 0.872 |
| 171 | 6ff7K | 0.643 | 8.470 | 0.691 | 6.548 | 0.487 | 15.188 | 0.688 |
| 172 | 6fonC | 0.619 | 18.307 | 0.623 | 17.937 | 0.613 | 17.956 | 0.624 |
| 173 | 6h3cC | 0.862 | 3.026 | 0.845 | 3.354 | 0.709 | 5.246 | 0.830 |
| 174 | 6hcyA | 0.882 | 2.895 | 0.874 | 3.153 | 0.739 | 5.820 | 0.852 |
| 175 | 6hpiA | 0.793 | 3.225 | 0.789 | 3.963 | 0.786 | 3.336 | 0.782 |
| 176 | 6htsD | 0.839 | 4.118 | 0.716 | 6.618 | 0.700 | 7.516 | 0.778 |
| 177 | 6i3yA | 0.774 | 4.455 | 0.780 | 6.933 | 0.818 | 3.436 | 0.883 |
| 178 | 6ip82z | 0.841 | 2.802 | 0.828 | 5.460 | 0.799 | 6.146 | 0.828 |
| 179 | 6k2yA | 0.784 | 8.880 | 0.792 | 8.983 | 0.780 | 8.888 | 0.791 |
| 180 | 6kmrA | 0.564 | 3.761 | 0.541 | 3.943 | 0.555 | 3.802 | 0.543 |
| 181 | 6kvgA | 0.599 | 14.948 | 0.478 | 17.263 | 0.398 | 14.951 | 0.490 |
| 182 | 6l9zP | 0.844 | 3.345 | 0.826 | 5.776 | 0.740 | 10.561 | 0.870 |
| 183 | 6mskB | 0.751 | 30.353 | 0.672 | 25.879 | 0.631 | 33.782 | 0.833 |
| 184 | 6nu2AQ | 0.711 | 5.792 | 0.756 | 4.878 | 0.562 | 7.162 | 0.873 |
| 185 | 6nuiA | 0.775 | 3.748 | 0.743 | 4.973 | 0.772 | 3.966 | 0.740 |
| 186 | 6o9l3 | 0.461 | 28.341 | 0.370 | 31.939 | 0.390 | 39.328 | 0.360 |
| 187 | 6o9lH | 0.843 | 2.413 | 0.851 | 2.217 | 0.787 | 3.139 | 0.833 |
| 188 | 6o9lQ | 0.401 | 60.652 | 0.368 | 36.068 | 0.373 | 31.624 | 0.366 |
| 189 | 6o9lT | 0.547 | 14.224 | 0.534 | 11.833 | 0.485 | 27.496 | 0.554 |
| 190 | 6pojA | 0.735 | 7.290 | 0.725 | 11.441 | 0.722 | 12.297 | 0.730 |
| 191 | 6qw668 | 0.753 | 3.521 | 0.747 | 3.628 | 0.766 | 3.114 | 0.753 |
| 192 | 6r7fE | 0.803 | 4.549 | 0.738 | 4.741 | 0.709 | 5.658 | 0.735 |
| 193 | 6r7fP | 0.825 | 9.784 | 0.816 | 6.248 | 0.782 | 8.165 | 0.868 |
| 194 | 6r7fQ | 0.810 | 3.351 | 0.819 | 3.444 | 0.776 | 5.816 | 0.831 |
| 195 | 6r7iD | 0.921 | 2.224 | 0.870 | 7.522 | 0.846 | 8.032 | 0.867 |
| 196 | 6r7nB | 0.786 | 11.963 | 0.788 | 10.844 | 0.714 | 6.412 | 0.786 |
| 197 | 6rw5Z | 0.690 | 3.611 | 0.641 | 5.669 | 0.429 | 9.096 | 0.823 |
|  |  | TM-score | RMSD | TM-score | RMSD | TM-score | RMSD |  |
| 198 | 6s7oD | 0.933 | 1.072 | 0.889 | 1.527 | 0.938 | 1.048 | 0.895 |
| 199 | 6thaA | 0.889 | 2.872 | 0.844 | 3.498 | 0.906 | 2.607 | 0.856 |
| 200 | 6u3gB | 0.614 | 23.982 | 0.630 | 15.051 | 0.600 | 24.019 | 0.897 |
| 201 | 6uhcB | 0.903 | 2.539 | 0.891 | 2.684 | 0.888 | 2.719 | 0.894 |
| 202 | 6uiwK | 0.838 | 5.722 | 0.794 | 5.197 | 0.832 | 5.155 | 0.844 |
| 203 | 6v22F | 0.842 | 2.781 | 0.819 | 2.770 | 0.732 | 3.577 | 0.855 |
| 204 | 6v4xC | 0.880 | 2.079 | 0.893 | 3.793 | 0.797 | 3.795 | 0.886 |
| 205 | 6vgcE | 0.888 | 2.089 | 0.823 | 4.884 | 0.782 | 12.693 | 0.840 |
| 206 | 6wm3O | 0.883 | 3.036 | 0.860 | 5.022 | 0.891 | 2.787 | 0.879 |
| 207 | 6wm4L | 0.798 | 2.228 | 0.607 | 5.806 | 0.722 | 6.956 | 0.599 |
| 208 | 6y0gSP | 0.860 | 2.682 | 0.847 | 5.094 | 0.843 | 2.331 | 0.856 |
| 209 | 6ybd8 | 0.607 | 8.048 | 0.624 | 8.069 | 0.567 | 8.276 | 0.633 |
| 210 | 6z3wF | 0.738 | 3.067 | 0.746 | 5.844 | 0.708 | 4.643 | 0.729 |
| 211 | 6zm56 | 0.837 | 5.901 | 0.803 | 5.174 | 0.758 | 9.757 | 0.795 |
| 212 | 6zm5A0 | 0.824 | 3.292 | 0.774 | 7.951 | 0.753 | 4.206 | 0.730 |
| 213 | 6zm7SR | 0.665 | 4.293 | 0.733 | 5.111 | 0.569 | 6.784 | 0.797 |
| 214 | 6zmwj | 0.760 | 5.002 | 0.624 | 6.805 | 0.626 | 6.483 | 0.616 |
| 215 | 6zp4K | 0.843 | 3.987 | 0.871 | 3.028 | 0.812 | 10.563 | 0.877 |
| 216 | 6zp4Z | 0.727 | 7.517 | 0.723 | 9.985 | 0.720 | 12.968 | 0.737 |
| 217 | 7bgbJ | 0.632 | 3.562 | 0.641 | 3.818 | 0.564 | 4.683 | 0.698 |
| 218 | 7cccB | 0.544 | 15.704 | 0.535 | 11.148 | 0.487 | 19.134 | 0.608 |
| 219 | 7cp9F | 0.805 | 1.716 | 0.697 | 2.435 | 0.541 | 3.890 | 0.681 |
| 220 | 7d59I | 0.417 | 12.754 | 0.380 | 11.883 | 0.376 | 12.930 | 0.381 |
| 221 | 7dmeA | 0.470 | 14.354 | 0.439 | 24.277 | 0.453 | 26.855 | 0.446 |
| 222 | 7drtA | 0.824 | 5.654 | 0.829 | 5.540 | 0.728 | 7.549 | 0.831 |
| 223 | 7dvq0 | 0.632 | 4.270 | 0.740 | 2.720 | 0.412 | 8.667 | 0.619 |
| 224 | 7dvq7 | 0.850 | 1.542 | 0.833 | 2.182 | 0.785 | 2.652 | 0.814 |
| 225 | 7dvqK | 0.880 | 2.197 | 0.864 | 9.417 | 0.843 | 7.243 | 0.875 |
| 226 | 7dwbE | 0.748 | 15.984 | 0.720 | 17.196 | 0.708 | 15.534 | 0.729 |
| 227 | 7emfK | 0.458 | 25.655 | 0.445 | 13.303 | 0.447 | 26.377 | 0.454 |
| 228 | 7ena2 | 0.808 | 7.002 | 0.820 | 5.951 | 0.743 | 8.350 | 0.848 |
| 229 | 7ena3 | 0.868 | 4.190 | 0.849 | 4.631 | 0.808 | 4.259 | 0.831 |
| 230 | 7ena4 | 0.736 | 6.792 | 0.709 | 5.611 | 0.712 | 8.618 | 0.689 |
|  |  | TM-score | RMSD | TM-score | RMSD | TM-score | RMSD |  |
| 231 | 7f3tA | 0.851 | 4.326 | 0.813 | 4.864 | 0.670 | 16.058 | 0.807 |
| 232 | 7k0oA | 0.910 | 3.373 | 0.878 | 6.756 | 0.857 | 11.160 | 0.891 |
| 233 | 7kzqF | 0.718 | 5.358 | 0.637 | 7.438 | 0.561 | 18.188 | 0.649 |
| 234 | 7kzvM | 0.824 | 3.826 | 0.717 | 10.108 | 0.750 | 5.029 | 0.725 |
| 235 | 7l20V | 0.778 | 6.161 | 0.853 | 2.825 | 0.748 | 4.471 | 0.885 |
| 236 | 7lbnm | 0.590 | 16.319 | 0.549 | 26.356 | 0.512 | 27.458 | 0.550 |
| 237 | 7lj1I | 0.851 | 8.782 | 0.868 | 8.841 | 0.852 | 8.932 | 0.865 |
| 238 | 7mq9SL | 0.771 | 13.464 | 0.745 | 8.091 | 0.685 | 10.954 | 0.895 |
| 239 | 7mq9SY | 0.562 | 18.086 | 0.538 | 11.784 | 0.479 | 27.402 | 0.837 |
| 240 | 7nvrk | 0.827 | 3.086 | 0.555 | 5.550 | 0.532 | 6.032 | 0.822 |
| 241 | 7nvrn | 0.772 | 4.538 | 0.796 | 3.028 | 0.573 | 6.282 | 0.792 |
| 242 | 7odtM | 0.770 | 8.446 | 0.763 | 12.228 | 0.785 | 7.045 | 0.884 |
| 243 | 7p2qB | 0.793 | 3.942 | 0.770 | 4.211 | 0.703 | 4.788 | 0.777 |
| 244 | 7rbsB | 0.830 | 2.371 | 0.841 | 2.274 | 0.740 | 3.473 | 0.831 |
| Average |  | 0.712 | 8.307 | 0.691 | 9.352 | 0.661 | 9.725 | 0.709 |

**Table S8.** CAMEO 1-month performance (2022/04/30∼2022/05/21)

| Method | ALL (N=59) |  | Easy (N=26) |  | Medium (N=30) |  | Hard (N=3) |  | Better |
| --- | --- | --- | --- | --- | --- | --- | --- | --- | --- |
|  | #TM_F | #TM_B | #TM_F | #TM_B | #TM_F | #TM_B | #TM_F | #TM_B |  |
|  | irst | est | irst | est | irst | est | irst | est |  |
| MultiSFold | 0.899 | 0.901 | 0.961 | 0.962 | 0.857 | 0.861 | 0.773 | 0.782 | - |
| PureAF2_orig | 0.892 | 0.892 | 0.954 | 0.954 | 0.853 | 0.853 | 0.774 | 0.774 | 27/56 |
| PureAF2_notemp | 0.896 | 0.896 | 0.955 | 0.955 | 0.858 | 0.858 | 0.774 | 0.774 | 30/59 |
| RoseTTAFold | 0.824 | 0.828 | 0.914 | 0.915 | 0.771 | 0.777 | 0.583 | 0.586 | 55/59 |
| IntFOLD6-TS | 0.749 | 0.765 | 0.911 | 0.919 | 0.651 | 0.672 | 0.341 | 0.371 | 40/41 |
| SWISS-MODEL | 0.729 | 0.733 | 0.910 | 0.910 | 0.608 | 0.612 | 0.334 | 0.334 | 52/58 |
| Best Single<br>Structural Template | 0.819 | 0.819 | 0.938 | 0.938 | 0.760 | 0.760 | 0.558 | 0.558 | 43/53 |

**Table S9.**
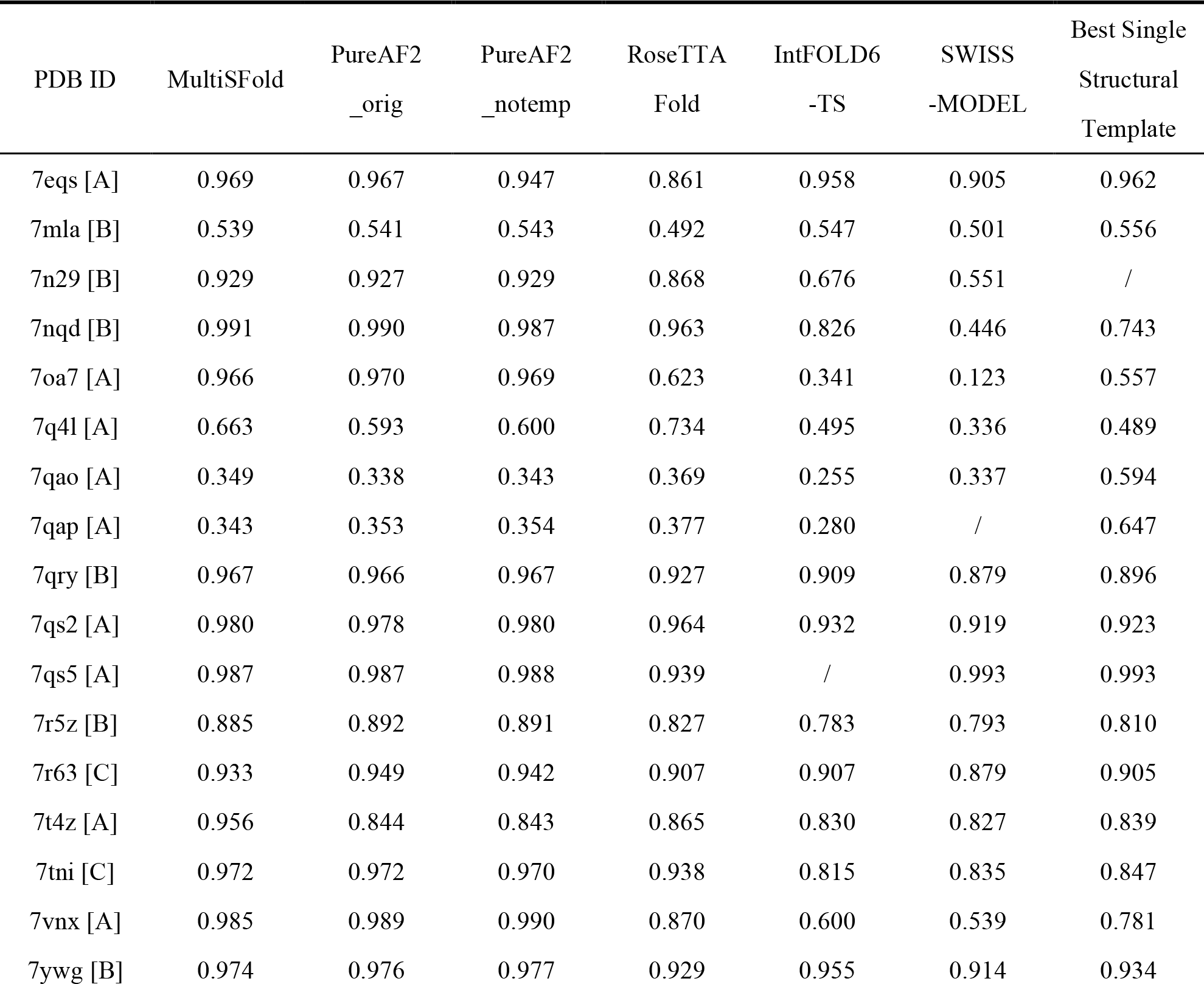
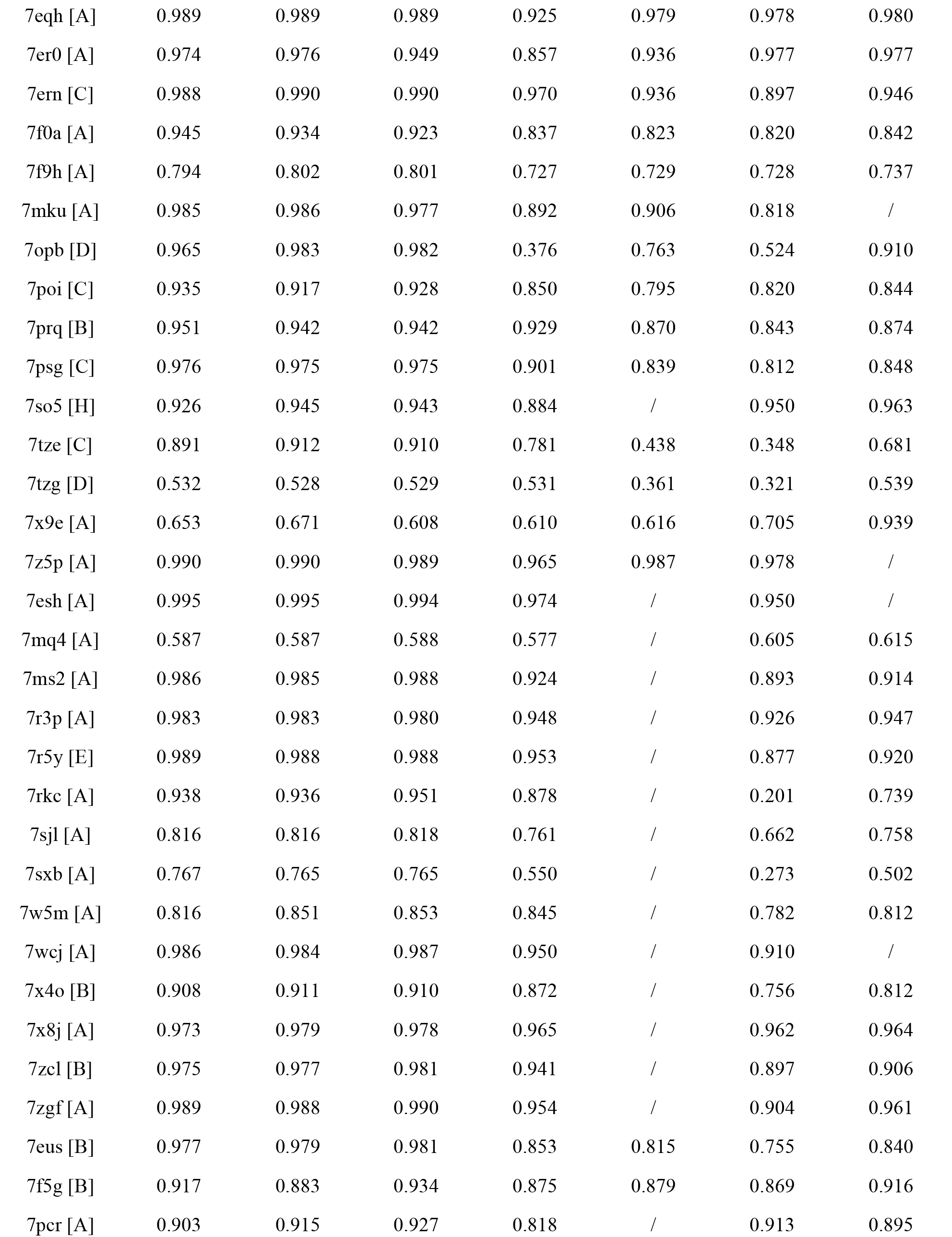
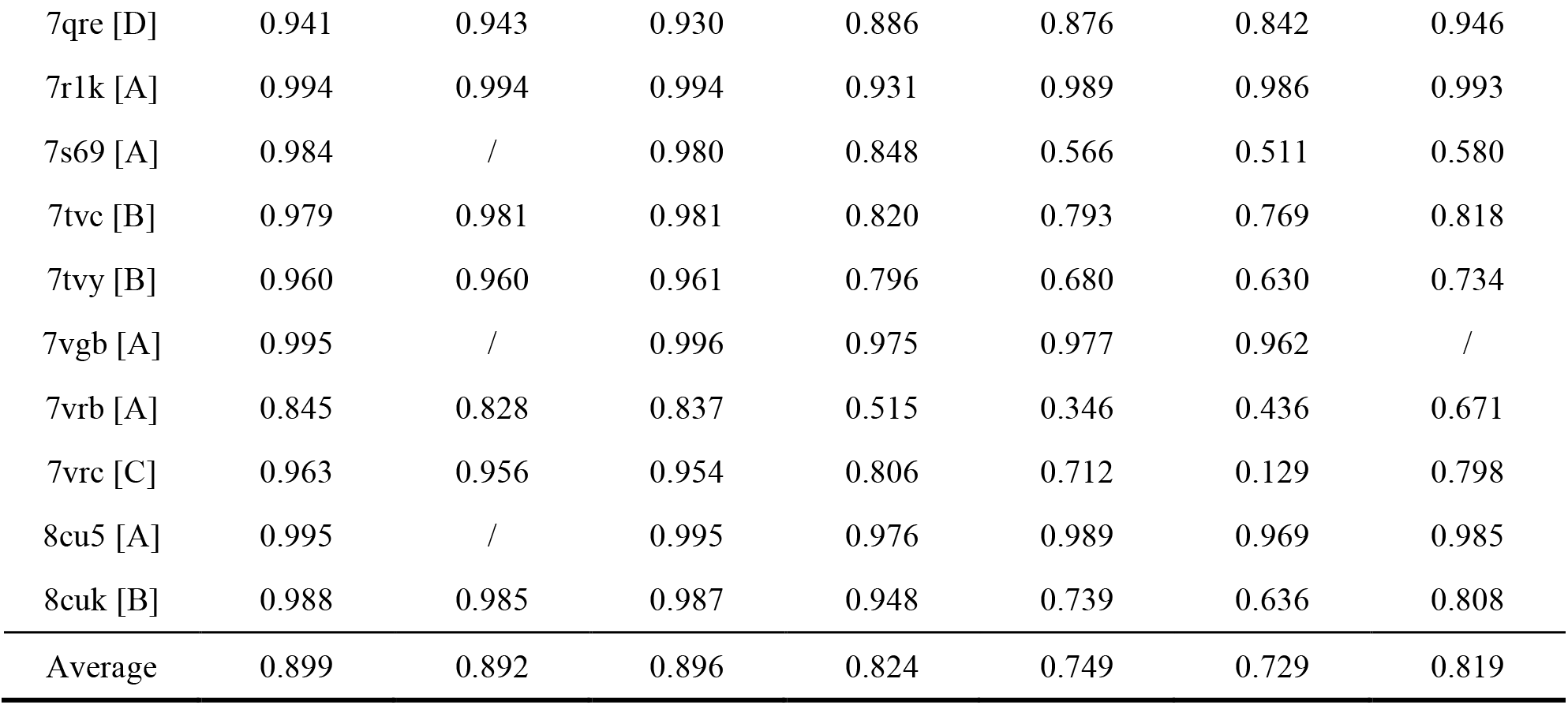
Details of CAMEO 1-month performance (2022/04/30∼2022/05/21).

**Table S10.** Summary of experimental results of component analysis.

| Method | TM-score | RMSD | #TM≥0.5 | #TM≥0.8 | P-value |
| --- | --- | --- | --- | --- | --- |
| MultiSFold | 0.712 | 8.307 | 214 | 95 | - |
| MultiSFold w/o iteration | 0.678 | 9.655 | 206 | 68 | 3.43E-33 |
| MultiSFold w/o loop | 0.690 | 9.418 | 211 | 78 | 3.16E-18 |

**Table S11.** Details of experimental results of component analysis.

| No. | PDB ID | MultiSFold |  | MultiSFold w/o iteration |  | MultiSFold w/o loop |  |
| --- | --- | --- | --- | --- | --- | --- | --- |
|  |  | TM-score | RMSD | TM-score | RMSD | TM-score | RMSD |
| 1 | 1bu9A | 0.876 | 3.622 | 0.880 | 2.436 | 0.849 | 3.951 |
| 2 | 1cksB | 0.727 | 16.330 | 0.707 | 16.155 | 0.717 | 16.583 |
| 3 | 1d7qA | 0.616 | 13.131 | 0.599 | 13.226 | 0.598 | 13.701 |
| 4 | 1dgnA | 0.884 | 1.611 | 0.886 | 1.525 | 0.864 | 1.740 |
| 5 | 1e03L | 0.902 | 7.208 | 0.893 | 9.669 | 0.887 | 8.859 |
| 6 | 1fwqA | 0.857 | 1.902 | 0.850 | 2.052 | 0.860 | 1.894 |
| 7 | 1griA | 0.568 | 5.720 | 0.574 | 5.615 | 0.567 | 5.583 |
| 8 | 1hp8A | 0.705 | 2.915 | 0.657 | 3.371 | 0.699 | 3.590 |
| 9 | 1hurA | 0.764 | 4.188 | 0.749 | 5.618 | 0.750 | 6.648 |
| 10 | 1j72A | 0.354 | 22.935 | 0.331 | 24.467 | 0.346 | 21.916 |
| 11 | 1nglA | 0.745 | 4.052 | 0.732 | 5.856 | 0.736 | 5.848 |
| 12 | 1nmvA | 0.679 | 8.918 | 0.654 | 16.534 | 0.652 | 16.706 |
| 13 | 1o0lA | 0.800 | 4.684 | 0.786 | 4.335 | 0.788 | 4.823 |
| 14 | 1oqyA | 0.282 | 46.304 | 0.241 | 41.741 | 0.221 | 39.265 |
| 15 | 1pc2A | 0.735 | 6.417 | 0.706 | 16.719 | 0.717 | 7.452 |
| 16 | 1q8kA | 0.688 | 4.998 | 0.578 | 8.408 | 0.671 | 5.220 |
| 17 | 1qklA | 0.503 | 10.884 | 0.487 | 11.715 | 0.504 | 9.106 |
| 18 | 1qttA | 0.843 | 2.283 | 0.805 | 2.786 | 0.824 | 2.431 |
| 19 | 1r6hA | 0.718 | 8.019 | 0.699 | 7.424 | 0.681 | 8.176 |
| 20 | 1r74B | 0.896 | 3.759 | 0.858 | 4.474 | 0.894 | 4.100 |
| 21 | 1ry7A | 0.872 | 4.262 | 0.852 | 7.059 | 0.864 | 3.819 |
| 22 | 1sgoA | 0.763 | 15.573 | 0.741 | 17.208 | 0.741 | 15.654 |
| 23 | 1sj6A | 0.835 | 1.784 | 0.820 | 1.883 | 0.816 | 1.915 |
| 24 | 1ttxA | 0.849 | 1.799 | 0.834 | 1.911 | 0.833 | 1.940 |
| 25 | 1w0bA | 0.765 | 5.467 | 0.753 | 5.167 | 0.747 | 6.799 |
|  |  | TM-score | RMSD | TM-score | RMSD | TM-score | RMSD |
| 26 | 1w0sC | 0.232 | 34.552 | 0.189 | 41.894 | 0.177 | 80.295 |
| 27 | 1wnjA | 0.842 | 2.700 | 0.831 | 3.797 | 0.832 | 2.541 |
| 28 | 1wxsA | 0.767 | 2.259 | 0.755 | 2.529 | 0.736 | 2.605 |
| 29 | 1xwnA | 0.854 | 5.916 | 0.847 | 5.552 | 0.856 | 5.084 |
| 30 | 1zj6A | 0.768 | 5.009 | 0.763 | 7.344 | 0.763 | 5.064 |
| 31 | 1zzaA | 0.340 | 17.565 | 0.305 | 26.437 | 0.306 | 21.760 |
| 32 | 2b95A | 0.848 | 2.044 | 0.837 | 2.034 | 0.830 | 1.940 |
| 33 | 2bidA | 0.627 | 7.914 | 0.606 | 8.567 | 0.627 | 9.276 |
| 34 | 2c9yA | 0.781 | 4.015 | 0.781 | 4.218 | 0.781 | 4.015 |
| 35 | 2e30A | 0.700 | 7.672 | 0.691 | 7.497 | 0.692 | 7.457 |
| 36 | 2g2bA | 0.677 | 11.792 | 0.653 | 15.185 | 0.657 | 12.988 |
| 37 | 2gdzA | 0.701 | 4.013 | 0.694 | 4.212 | 0.697 | 4.125 |
| 38 | 2gf5A | 0.661 | 6.397 | 0.543 | 7.531 | 0.605 | 6.695 |
| 39 | 2gowA | 0.544 | 15.895 | 0.546 | 10.782 | 0.547 | 9.779 |
| 40 | 2gpqA | 0.684 | 9.507 | 0.671 | 10.134 | 0.666 | 9.170 |
| 41 | 2hdeA | 0.684 | 8.661 | 0.678 | 9.496 | 0.686 | 8.373 |
| 42 | 2hr9A | 0.835 | 2.732 | 0.766 | 4.449 | 0.841 | 2.675 |
| 43 | 2hynC | 0.485 | 8.649 | 0.489 | 8.615 | 0.491 | 14.902 |
| 44 | 2i3bA | 0.872 | 2.236 | 0.856 | 2.418 | 0.863 | 2.344 |
| 45 | 2j3tB | 0.865 | 3.407 | 0.842 | 3.276 | 0.849 | 3.468 |
| 46 | 2jdfA | 0.635 | 3.789 | 0.630 | 3.827 | 0.634 | 3.799 |
| 47 | 2jnjB | 0.691 | 7.381 | 0.670 | 7.003 | 0.693 | 6.597 |
| 48 | 2jr7A | 0.757 | 2.949 | 0.732 | 3.912 | 0.738 | 3.802 |
| 49 | 2jrfA | 0.580 | 16.749 | 0.560 | 20.526 | 0.557 | 17.425 |
| 50 | 2k07A | 0.881 | 2.173 | 0.877 | 2.139 | 0.876 | 2.160 |
| 51 | 2k21A | 0.287 | 19.442 | 0.233 | 23.994 | 0.285 | 20.074 |
| 52 | 2k7kA | 0.891 | 1.336 | 0.875 | 2.343 | 0.897 | 1.298 |
| 53 | 2kg4A | 0.762 | 8.539 | 0.746 | 8.155 | 0.748 | 8.581 |
| 54 | 2kkwA | 0.374 | 26.970 | 0.287 | 21.956 | 0.373 | 24.915 |
| 55 | 2kn6A | 0.493 | 10.325 | 0.468 | 13.816 | 0.508 | 10.279 |
| 56 | 2kr0A | 0.317 | 33.122 | 0.287 | 42.932 | 0.292 | 68.431 |
| 57 | 2l2oA | 0.883 | 1.555 | 0.872 | 1.592 | 0.871 | 1.629 |
| 58 | 2l4sA | 0.733 | 5.249 | 0.718 | 5.751 | 0.706 | 5.249 |
|  |  | TM-score | RMSD | TM-score | RMSD | TM-score | RMSD |
| 59 | 2l51A | 0.726 | 3.502 | 0.715 | 3.263 | 0.694 | 3.611 |
| 60 | 2l6lA | 0.531 | 12.339 | 0.513 | 13.307 | 0.526 | 12.773 |
| 61 | 2l7bA | 0.459 | 66.651 | 0.448 | 67.750 | 0.440 | 66.041 |
| 62 | 2lcpA | 0.718 | 4.765 | 0.711 | 4.682 | 0.702 | 4.766 |
| 63 | 2lgrA | 0.899 | 1.784 | 0.865 | 4.627 | 0.897 | 1.801 |
| 64 | 2ljkA | 0.700 | 7.296 | 0.672 | 8.229 | 0.682 | 9.883 |
| 65 | 2lm5A | 0.740 | 5.666 | 0.720 | 5.896 | 0.717 | 5.874 |
| 66 | 2lomA | 0.357 | 12.492 | 0.312 | 14.498 | 0.361 | 12.488 |
| 67 | 2loqA | 0.287 | 10.214 | 0.251 | 13.394 | 0.293 | 8.299 |
| 68 | 2lorA | 0.445 | 10.107 | 0.387 | 9.719 | 0.449 | 10.155 |
| 69 | 2losA | 0.286 | 19.632 | 0.286 | 19.411 | 0.255 | 19.957 |
| 70 | 2lqnA | 0.356 | 28.162 | 0.348 | 25.328 | 0.349 | 36.782 |
| 71 | 2lr1A | 0.835 | 4.450 | 0.817 | 5.937 | 0.824 | 5.611 |
| 72 | 2m0qA | 0.277 | 30.867 | 0.277 | 40.671 | 0.272 | 30.475 |
| 73 | 2m0rB | 0.684 | 10.949 | 0.669 | 7.374 | 0.684 | 6.412 |
| 74 | 2m65A | 0.847 | 3.301 | 0.845 | 5.372 | 0.839 | 5.634 |
| 75 | 2mmgA | 0.822 | 7.624 | 0.797 | 12.290 | 0.803 | 15.639 |
| 76 | 2mouA | 0.889 | 4.324 | 0.883 | 3.063 | 0.885 | 4.396 |
| 77 | 2mphA | 0.813 | 5.131 | 0.784 | 5.853 | 0.813 | 5.131 |
| 78 | 2mt6A | 0.738 | 7.846 | 0.718 | 8.145 | 0.744 | 7.908 |
| 79 | 2mxca | 0.757 | 3.840 | 0.739 | 9.971 | 0.762 | 4.123 |
| 80 | 2mzeA | 0.831 | 5.691 | 0.822 | 5.972 | 0.819 | 6.450 |
| 81 | 2n1aA | 0.605 | 8.313 | 0.576 | 10.025 | 0.577 | 8.194 |
| 82 | 2ndjA | 0.344 | 11.349 | 0.295 | 19.944 | 0.322 | 9.130 |
| 83 | 2octA | 0.474 | 16.196 | 0.476 | 16.357 | 0.470 | 16.178 |
| 84 | 2ozxA | 0.873 | 2.142 | 0.862 | 2.254 | 0.857 | 2.387 |
| 85 | 2p03A | 0.336 | 26.332 | 0.318 | 30.734 | 0.343 | 26.267 |
| 86 | 2qs9A | 0.529 | 6.613 | 0.526 | 6.552 | 0.521 | 6.267 |
| 87 | 2rmkA | 0.804 | 7.778 | 0.802 | 7.658 | 0.797 | 8.781 |
| 88 | 2rnbA | 0.612 | 8.118 | 0.585 | 11.558 | 0.607 | 10.010 |
| 89 | 2rsvA | 0.757 | 21.237 | 0.749 | 16.697 | 0.752 | 23.543 |
| 90 | 2rvqC | 0.526 | 17.369 | 0.513 | 13.128 | 0.513 | 18.582 |
| 91 | 2rvqD | 0.645 | 7.151 | 0.606 | 16.804 | 0.583 | 27.315 |
|  |  | TM-score | RMSD | TM-score | RMSD | TM-score | RMSD |
| 92 | 2x4dA | 0.697 | 4.077 | 0.694 | 4.119 | 0.694 | 4.101 |
| 93 | 2ymbD | 0.740 | 6.215 | 0.686 | 8.432 | 0.740 | 6.146 |
| 94 | 3epyA | 0.492 | 20.104 | 0.488 | 20.264 | 0.483 | 20.165 |
| 95 | 3ggrB | 0.874 | 3.016 | 0.876 | 2.851 | 0.876 | 2.974 |
| 96 | 3ggrC | 0.815 | 3.428 | 0.813 | 3.436 | 0.814 | 3.449 |
| 97 | 3j07V | 0.406 | 24.766 | 0.402 | 23.252 | 0.396 | 25.286 |
| 98 | 3j0sW | 0.888 | 1.865 | 0.869 | 2.094 | 0.879 | 2.050 |
| 99 | 3k2sA | 0.262 | 42.600 | 0.228 | 39.949 | 0.262 | 42.579 |
| 100 | 3nsiB | 0.644 | 6.924 | 0.629 | 8.826 | 0.628 | 6.992 |
| 101 | 3pgwF | 0.833 | 2.995 | 0.808 | 3.391 | 0.781 | 3.659 |
| 102 | 3pgwG | 0.836 | 2.374 | 0.793 | 3.020 | 0.806 | 3.176 |
| 103 | 3pgwH | 0.634 | 4.992 | 0.617 | 5.484 | 0.622 | 4.717 |
| 104 | 3pgwY | 0.792 | 3.452 | 0.753 | 4.904 | 0.763 | 3.652 |
| 105 | 3rqgC | 0.384 | 13.348 | 0.389 | 12.433 | 0.388 | 13.469 |
| 106 | 3vkmA | 0.565 | 14.651 | 0.557 | 14.468 | 0.565 | 14.651 |
| 107 | 3wsoA | 0.886 | 2.707 | 0.870 | 2.941 | 0.891 | 2.735 |
| 108 | 3wvzA | 0.777 | 3.044 | 0.707 | 8.309 | 0.769 | 3.061 |
| 109 | 4a6eA | 0.821 | 3.969 | 0.807 | 4.389 | 0.815 | 4.050 |
| 110 | 4aj5T | 0.601 | 7.489 | 0.562 | 12.106 | 0.579 | 16.203 |
| 111 | 4aqhB | 0.901 | 6.149 | 0.887 | 7.589 | 0.904 | 6.191 |
| 112 | 4b4oF | 0.618 | 5.698 | 0.616 | 5.726 | 0.605 | 5.850 |
| 113 | 4behA | 0.468 | 16.370 | 0.451 | 31.025 | 0.463 | 22.087 |
| 114 | 4behB | 0.465 | 15.178 | 0.431 | 13.754 | 0.430 | 15.029 |
| 115 | 4d5nA | 0.734 | 6.379 | 0.721 | 6.788 | 0.711 | 5.974 |
| 116 | 4dinB | 0.671 | 22.538 | 0.640 | 15.400 | 0.665 | 20.851 |
| 117 | 4djcA | 0.873 | 1.915 | 0.765 | 2.922 | 0.671 | 3.674 |
| 118 | 4g1tB | 0.794 | 8.232 | 0.784 | 8.347 | 0.800 | 8.128 |
| 119 | 4kfzB | 0.532 | 9.947 | 0.525 | 10.979 | 0.495 | 10.352 |
| 120 | 4uybA | 0.844 | 3.245 | 0.839 | 3.298 | 0.843 | 3.245 |
| 121 | 4v6xAb | 0.831 | 1.659 | 0.759 | 2.548 | 0.748 | 2.416 |
| 122 | 4v6xCE | 0.616 | 15.294 | 0.534 | 25.653 | 0.553 | 25.712 |
| 123 | 4v6xCK | 0.594 | 4.767 | 0.505 | 5.558 | 0.582 | 5.219 |
| 124 | 4v6xCr | 0.659 | 7.210 | 0.654 | 6.771 | 0.663 | 6.352 |
|  |  | TM-score | RMSD | TM-score | RMSD | TM-score | RMSD |
| 125 | 4x4wA | 0.901 | 2.841 | 0.887 | 3.419 | 0.888 | 3.419 |
| 126 | 5a9qT | 0.835 | 4.480 | 0.833 | 4.543 | 0.829 | 4.537 |
| 127 | 5aj0Au | 0.863 | 2.903 | 0.814 | 3.218 | 0.858 | 2.669 |
| 128 | 5ancJ | 0.632 | 8.381 | 0.500 | 12.958 | 0.616 | 7.509 |
| 129 | 5g04B | 0.785 | 2.923 | 0.766 | 3.685 | 0.767 | 2.859 |
| 130 | 5hydD | 0.618 | 4.728 | 0.592 | 4.735 | 0.623 | 4.504 |
| 131 | 5iy6I | 0.545 | 5.164 | 0.450 | 14.551 | 0.504 | 9.323 |
| 132 | 5iybM | 0.779 | 6.153 | 0.762 | 8.724 | 0.783 | 6.413 |
| 133 | 5j0aA | 0.747 | 14.688 | 0.741 | 14.768 | 0.744 | 14.967 |
| 134 | 5l4kP | 0.798 | 6.579 | 0.752 | 7.763 | 0.771 | 9.503 |
| 135 | 5l4kQ | 0.839 | 4.457 | 0.770 | 7.673 | 0.796 | 4.546 |
| 136 | 5l4kU | 0.691 | 5.844 | 0.628 | 14.144 | 0.676 | 7.411 |
| 137 | 5l4kV | 0.738 | 5.807 | 0.712 | 5.769 | 0.706 | 7.517 |
| 138 | 5ln3D | 0.885 | 3.510 | 0.884 | 3.639 | 0.874 | 3.727 |
| 139 | 5lsjC | 0.628 | 10.827 | 0.613 | 15.053 | 0.497 | 16.731 |
| 140 | 5nw4h | 0.903 | 2.051 | 0.892 | 2.704 | 0.894 | 2.415 |
| 141 | 5tbyE | 0.605 | 6.483 | 0.421 | 10.222 | 0.630 | 5.820 |
| 142 | 5ujjA | 0.884 | 6.543 | 0.853 | 19.436 | 0.865 | 9.261 |
| 143 | 5wtqA | 0.809 | 2.132 | 0.793 | 2.502 | 0.747 | 2.859 |
| 144 | 5xtcj | 0.719 | 2.979 | 0.503 | 10.845 | 0.501 | 5.926 |
| 145 | 5xteN | 0.681 | 7.461 | 0.592 | 9.940 | 0.637 | 5.635 |
| 146 | 5xthh | 0.676 | 3.928 | 0.675 | 3.845 | 0.666 | 4.388 |
| 147 | 5xthW | 0.755 | 9.838 | 0.731 | 6.524 | 0.735 | 10.082 |
| 148 | 5xtiBb | 0.527 | 9.476 | 0.473 | 20.181 | 0.498 | 7.089 |
| 149 | 5xtiBg | 0.807 | 3.192 | 0.742 | 3.916 | 0.743 | 3.621 |
| 150 | 5xtiBn | 0.562 | 3.251 | 0.506 | 4.044 | 0.500 | 4.373 |
| 151 | 5xylA | 0.633 | 10.387 | 0.627 | 9.416 | 0.626 | 10.234 |
| 152 | 5y39D | 0.479 | 5.299 | 0.474 | 4.716 | 0.466 | 5.369 |
| 153 | 5yfgA | 0.821 | 2.871 | 0.794 | 3.034 | 0.820 | 2.898 |
| 154 | 5z62H | 0.864 | 1.490 | 0.807 | 2.045 | 0.835 | 1.625 |
| 155 | 5z62I | 0.817 | 2.066 | 0.749 | 2.226 | 0.827 | 1.990 |
| 156 | 5z62N | 0.599 | 3.845 | 0.579 | 3.887 | 0.570 | 3.986 |
| 157 | 6ahuE | 0.871 | 2.032 | 0.859 | 2.208 | 0.851 | 2.461 |
|  |  | TM-score | RMSD | TM-score | RMSD | TM-score | RMSD |
| 158 | 6ahuG | 0.733 | 6.404 | 0.723 | 6.298 | 0.733 | 6.193 |
| 159 | 6ahuH | 0.807 | 4.739 | 0.790 | 4.554 | 0.793 | 4.932 |
| 160 | 6bc8A | 0.877 | 3.263 | 0.866 | 3.909 | 0.871 | 3.491 |
| 161 | 6cmoA | 0.588 | 9.321 | 0.552 | 13.897 | 0.577 | 9.661 |
| 162 | 6cmoR | 0.912 | 2.099 | 0.896 | 2.446 | 0.907 | 2.166 |
| 163 | 6d6rI | 0.872 | 2.163 | 0.640 | 5.513 | 0.762 | 4.463 |
| 164 | 6dk0C | 0.883 | 3.589 | 0.868 | 5.255 | 0.881 | 3.437 |
| 165 | 6fec3 | 0.623 | 10.744 | 0.594 | 10.794 | 0.577 | 12.559 |
| 166 | 6fec8 | 0.684 | 5.521 | 0.673 | 5.697 | 0.671 | 5.729 |
| 167 | 6fecG | 0.818 | 4.153 | 0.805 | 3.626 | 0.819 | 4.305 |
| 168 | 6fecQ | 0.855 | 1.993 | 0.831 | 2.227 | 0.841 | 2.155 |
| 169 | 6fecr | 0.832 | 2.973 | 0.828 | 2.829 | 0.825 | 2.939 |
| 170 | 6fecX | 0.853 | 2.463 | 0.846 | 2.555 | 0.837 | 2.620 |
| 171 | 6ff7K | 0.643 | 8.470 | 0.581 | 8.477 | 0.602 | 8.718 |
| 172 | 6fonC | 0.619 | 18.307 | 0.619 | 18.368 | 0.616 | 18.309 |
| 173 | 6h3cC | 0.862 | 3.026 | 0.855 | 3.145 | 0.860 | 3.056 |
| 174 | 6hcyA | 0.882 | 2.895 | 0.736 | 6.247 | 0.721 | 5.535 |
| 175 | 6hpiA | 0.793 | 3.225 | 0.790 | 3.802 | 0.794 | 3.606 |
| 176 | 6htsD | 0.839 | 4.118 | 0.758 | 6.468 | 0.840 | 4.194 |
| 177 | 6i3yA | 0.774 | 4.455 | 0.752 | 3.452 | 0.717 | 8.317 |
| 178 | 6ip82z | 0.841 | 2.802 | 0.794 | 3.651 | 0.816 | 2.979 |
| 179 | 6k2yA | 0.784 | 8.880 | 0.773 | 8.766 | 0.781 | 8.852 |
| 180 | 6kmrA | 0.564 | 3.761 | 0.557 | 3.831 | 0.556 | 3.812 |
| 181 | 6kvgA | 0.599 | 14.948 | 0.443 | 20.396 | 0.555 | 17.683 |
| 182 | 6l9zP | 0.844 | 3.345 | 0.825 | 3.815 | 0.794 | 2.849 |
| 183 | 6mskB | 0.751 | 30.353 | 0.632 | 34.667 | 0.755 | 32.282 |
| 184 | 6nu2AQ | 0.711 | 5.792 | 0.642 | 6.092 | 0.733 | 7.000 |
| 185 | 6nuiA | 0.775 | 3.748 | 0.773 | 3.701 | 0.762 | 3.891 |
| 186 | 6o9l3 | 0.461 | 28.341 | 0.413 | 20.240 | 0.397 | 29.005 |
| 187 | 6o9lH | 0.843 | 2.413 | 0.805 | 2.797 | 0.843 | 2.421 |
| 188 | 6o9lQ | 0.401 | 60.652 | 0.374 | 32.038 | 0.364 | 45.776 |
| 189 | 6o9lT | 0.547 | 14.224 | 0.504 | 22.617 | 0.589 | 9.428 |
| 190 | 6pojA | 0.735 | 7.290 | 0.709 | 10.157 | 0.711 | 10.367 |
|  |  | TM-score | RMSD | TM-score | RMSD | TM-score | RMSD |
| 191 | 6qw668 | 0.753 | 3.521 | 0.733 | 3.615 | 0.744 | 3.598 |
| 192 | 6r7fE | 0.803 | 4.549 | 0.726 | 4.814 | 0.707 | 5.163 |
| 193 | 6r7fP | 0.825 | 9.784 | 0.776 | 8.438 | 0.805 | 7.944 |
| 194 | 6r7fQ | 0.810 | 3.351 | 0.785 | 3.939 | 0.791 | 3.794 |
| 195 | 6r7iD | 0.921 | 2.224 | 0.869 | 4.157 | 0.871 | 6.183 |
| 196 | 6r7nB | 0.786 | 11.963 | 0.760 | 13.252 | 0.765 | 12.341 |
| 197 | 6rw5Z | 0.690 | 3.611 | 0.633 | 4.252 | 0.569 | 6.787 |
| 198 | 6s7oD | 0.933 | 1.072 | 0.876 | 1.605 | 0.908 | 1.271 |
| 199 | 6thaA | 0.889 | 2.872 | 0.854 | 3.419 | 0.886 | 2.905 |
| 200 | 6u3gB | 0.614 | 23.982 | 0.598 | 24.795 | 0.628 | 24.066 |
| 201 | 6uhcB | 0.903 | 2.539 | 0.886 | 2.734 | 0.888 | 2.698 |
| 202 | 6uiwK | 0.838 | 5.722 | 0.782 | 6.421 | 0.825 | 5.695 |
| 203 | 6v22F | 0.842 | 2.781 | 0.781 | 3.885 | 0.824 | 2.946 |
| 204 | 6v4xC | 0.880 | 2.079 | 0.844 | 3.409 | 0.855 | 3.827 |
| 205 | 6vgcE | 0.888 | 2.089 | 0.877 | 2.208 | 0.869 | 2.277 |
| 206 | 6wm3O | 0.883 | 3.036 | 0.720 | 6.857 | 0.880 | 3.073 |
| 207 | 6wm4L | 0.798 | 2.228 | 0.700 | 6.453 | 0.798 | 2.228 |
| 208 | 6y0gSP | 0.860 | 2.682 | 0.821 | 5.017 | 0.816 | 5.148 |
| 209 | 6ybd8 | 0.607 | 8.048 | 0.600 | 8.296 | 0.612 | 7.983 |
| 210 | 6z3wF | 0.738 | 3.067 | 0.703 | 4.525 | 0.711 | 3.323 |
| 211 | 6zm56 | 0.837 | 5.901 | 0.803 | 8.413 | 0.875 | 4.211 |
| 212 | 6zm5A0 | 0.824 | 3.292 | 0.751 | 4.275 | 0.849 | 2.652 |
| 213 | 6zm7SR | 0.665 | 4.293 | 0.585 | 8.702 | 0.582 | 10.340 |
| 214 | 6zmwj | 0.760 | 5.002 | 0.687 | 5.458 | 0.636 | 6.141 |
| 215 | 6zp4K | 0.843 | 3.987 | 0.831 | 6.735 | 0.835 | 8.992 |
| 216 | 6zp4Z | 0.727 | 7.517 | 0.737 | 6.325 | 0.706 | 12.117 |
| 217 | 7bgbJ | 0.632 | 3.562 | 0.568 | 3.258 | 0.518 | 5.558 |
| 218 | 7cccB | 0.544 | 15.704 | 0.473 | 20.600 | 0.531 | 24.946 |
| 219 | 7cp9F | 0.805 | 1.716 | 0.745 | 2.244 | 0.772 | 1.738 |
| 220 | 7d59I | 0.417 | 12.754 | 0.361 | 19.512 | 0.380 | 16.969 |
| 221 | 7dmeA | 0.470 | 14.354 | 0.424 | 24.890 | 0.482 | 13.228 |
| 222 | 7drtA | 0.824 | 5.654 | 0.816 | 5.656 | 0.816 | 5.674 |
| 223 | 7dvq0 | 0.632 | 4.270 | 0.617 | 4.363 | 0.636 | 4.259 |
|  |  | TM-score | RMSD | TM-score | RMSD | TM-score | RMSD |
| 224 | 7dvq7 | 0.850 | 1.542 | 0.809 | 2.093 | 0.821 | 1.776 |
| 225 | 7dvqK | 0.880 | 2.197 | 0.867 | 8.783 | 0.877 | 2.209 |
| 226 | 7dwbE | 0.748 | 15.984 | 0.717 | 17.220 | 0.733 | 22.283 |
| 227 | 7emfK | 0.458 | 25.655 | 0.445 | 25.647 | 0.455 | 25.999 |
| 228 | 7ena2 | 0.808 | 7.002 | 0.806 | 6.816 | 0.810 | 6.142 |
| 229 | 7ena3 | 0.868 | 4.190 | 0.851 | 4.835 | 0.829 | 4.201 |
| 230 | 7ena4 | 0.736 | 6.792 | 0.699 | 10.845 | 0.703 | 7.763 |
| 231 | 7f3tA | 0.851 | 4.326 | 0.803 | 5.156 | 0.809 | 4.890 |
| 232 | 7k0oA | 0.910 | 3.373 | 0.864 | 18.451 | 0.869 | 12.098 |
| 233 | 7kzqF | 0.718 | 5.358 | 0.687 | 5.316 | 0.624 | 7.130 |
| 234 | 7kzvM | 0.824 | 3.826 | 0.676 | 16.363 | 0.702 | 5.484 |
| 235 | 7l20V | 0.778 | 6.161 | 0.690 | 9.523 | 0.741 | 6.772 |
| 236 | 7lbnm | 0.590 | 16.319 | 0.548 | 13.802 | 0.570 | 25.667 |
| 237 | 7lj1I | 0.851 | 8.782 | 0.850 | 8.898 | 0.847 | 8.693 |
| 238 | 7mq9SL | 0.771 | 13.464 | 0.726 | 5.632 | 0.746 | 13.758 |
| 239 | 7mq9SY | 0.562 | 18.086 | 0.511 | 27.453 | 0.570 | 20.497 |
| 240 | 7nvrk | 0.827 | 3.086 | 0.730 | 3.693 | 0.617 | 5.012 |
| 241 | 7nvrn | 0.772 | 4.538 | 0.626 | 5.577 | 0.585 | 5.823 |
| 242 | 7odtM | 0.770 | 8.446 | 0.755 | 13.408 | 0.749 | 15.199 |
| 243 | 7p2qB | 0.793 | 3.942 | 0.796 | 3.765 | 0.790 | 3.928 |
| 244 | 7rbsB | 0.830 | 2.371 | 0.825 | 2.492 | 0.796 | 2.558 |
| Average |  | 0.712 | 8.307 | 0.678 | 9.655 | 0.690 | 9.418 |

## References

Boocock,G.R., et al (2003) Mutations in SBDS are associated with Shwachman-Diamond syndrome. Nature Genetics, 33, 97–101.

Boehr,D.D., et al (2009) The role of dynamic conformational ensembles in biomolecular recognition. Nature Chemical Biology, 5, 789–796.

Baek,M., et al (2021) Accurate prediction of protein structures and interactions using a three-track neural network. Science, 373, 871–876.

Cournia,Z., et al (2015) Membrane Protein Structure, Function, and Dynamics: a Perspective from Experiments and Theory. The Journal of Membrane Biology, 248, 611–640.

Campbell,E., et al (2016) The role of protein dynamics in the evolution of new enzyme function. Nature Chemical Biology, 12, 944–950.

de Groot,B.L., et al (1997) Prediction of protein conformational freedom from distance constraints. Proteins: Structure Function and Bioinformatics, 29, 240251.

de Groot,B.L., et al (1998) Domain motions in bacteriophage T4 lysozyme: a comparison between molecular dynamics and crystallographic data. Proteins: Structure Function and Bioinformatics, 31, 116–127.

de Groot,B.L., et al (1999) Conformational changes in the chaperonin GroEL: new insights into the allosteric mechanism. Journal of Molecular Biology, 286, 12411249.

Drew,D. and Boudker,O. (2016) Shared Molecular Mechanisms of Membrane Transporters. Annual Review of Biochemistry, 85, 543–572.

Del Alamo,D., et al (2022) Sampling alternative conformational states of transporters and receptors with AlphaFold2. Elife, 11, e75751.

Finch,A.J., et al (2011) Uncoupling of GTP hydrolysis from eIF6 release on the ribosome causes Shwachman-Diamond syndrome. Genes & Development, 25, 917929.

Fu,L., et al (2012) CD-HIT: Accelerated for clustering the next-generation sequencing data. Bioinformatics, 28, 3150–3152.

Feng,Q.Q., et al (2022) Construct a variable-length fragment library for de novo protein structure prediction. Briefings in Bioinformatics, 23, bbac086.

Garcia,C.K., et al (1994) Molecular characterization of a membrane transporter for lactate, pyruvate, and other monocarboxylates: implications for the Cori cycle. Cell, 76, 865–873.

Greener,J.G., et al (2017) Predicting Protein Dynamics and Allostery Using Multi-Protein Atomic Distance Constraints. Structure, 25, 546–558.

Henzler-Wildman,K.A., et al (2007) Intrinsic motions along an enzymatic reaction trajectory. Nature, 450, 838–844.

Hong,Z., et al (2010) A novel method for adaptive determination clusters number based on N-order nearest neighbor. Proceedings of the 29th Chinese Control Conference, 3007-3011.

Jumper,J., et al (2021) Highly accurate protein structure prediction with AlphaFold. Nature, 596, 583–589.

Jones,D.T. and Thornton,J.M. (2022) The impact of AlphaFold2 one year on. Nature Methods, 19, 15–20.

Kabsch,W. and Sander,C. (1983) Dictionary of protein secondary structure: pattern recognition of hydrogen-bonded and geometrical features. Biopolymers: Original Research on Biomolecules, 22, 2577–2637.

Kazmier,K., et al (2017) Alternating access mechanisms of LeuT-fold transporters: trailblazing towards the promised energy landscapes. Current Opinion in Structural Biology, 45, 100–108.

Liu,J., et al (2020) CGLFold: a contact-assisted de novo protein structure prediction using global exploration and loop perturbation sampling algorithm. Bioinformatics, 36, 2443–2450.

Liu,J., et al (2022) A de novo protein structure prediction by iterative partition sampling, topology adjustment and residue-level distance deviation optimization. Bioinformatics, 38, 99–107.

Modi,V. and Dunbrack,R.L.,Jr. (2019) Defining a new nomenclature for the structures of active and inactive kinases. Proceedings of the National Academy of Sciences, 116, 6818–6827.

Peng,C.X., et al (2023) Multiple conformational states assembly of multidomain proteins using evolutionary algorithm based on structural analogues and sequential homologues. bioRxiv, doi: https://doi.org/10.1101/2023.01.15.524086.

Ritzhaupt,A., et al (1998) Identification and characterization of a monocarboxylate transporter (MCT1) in pig and human colon: its potential to transport l-lactate as well as butyrate. The Journal of physiology, 513, 719–732.

Ramanathan,A., et al (2014) Protein conformational populations and functionally relevant substates. Accounts of chemical research, 47, 149–56.

Senger,B., et al (2001) The nucle(ol)ar Tif6p and Efl1p are required for a late cytoplasmic step of ribosome synthesis. Molecular Cell, 8, 1363–1373.

Seeliger,D., et al (2007) Geometry-based sampling of conformational transitions in proteins. Structure, 15, 1482–92.

Shaw,D.E., et al (2010) Atomic-level characterization of the structural dynamics of proteins. Science, 330, 341–346.

Skolnick,J., et al (2021) AlphaFold 2: Why It Works and Its Implications for Understanding the Relationships of Protein Sequence, Structure, and Function. Journal of Chemical Information and Modeling, 61, 4827–4831.

Subramaniam,S. and Kleywegt,G.J. (2022) A paradigm shift in structural biology. Nature Methods, 19, 20–23.

Tunyasuvunakool,K. et al (2021) Highly accurate protein structure prediction for the human proteome. Nature, 596, 590-596.

Thornton,J.M., et al (2021) AlphaFold heralds a data-driven revolution in biology and medicine. Nature Methods, 27, 1666–1669.

Tunyasuvunakool,K., et al (2022) AlphaFold Protein Structure Database: massively expanding the structural coverage of protein-sequence space with high-accuracy models. Nucleic Acids Research, 50, D439–D444.

Turkan,H., et al (2022) Prediction of allosteric communication pathways in proteins. Bioinformatics, 38, 3590–3599.

Weis,F., et al (2015) Mechanism of eIF6 release from the nascent 60S ribosomal subunit. Nature Structural & Molecular Biology, 22, 914–919.

Weis,W.I. and Kobilka,B.K. (2018) The Molecular Basis of G ProteinCoupled Receptor Activation. Annual Review of Biochemistry, 87, 897–919.

Wang,N., et al (2021) Structural basis of human monocarboxylate transporter 1 inhibition by anti-cancer drug candidates. Cell, 184, 370–383.

Xie,T., et al (2020) Conformational states dynamically populated by a kinase determine its function. Science, 370, eabc2754.

Zhang,Y. and Skolnick,J. (2004) Scoring function for automated assessment of protein structure template quality. Proteins, 57, 702–710.

Zhang,Y. and Skolnick,J. (2005) TM-align: a protein structure alignment algorithm based on the TM-score. Nucleic Acids Research, 33, 2302–2309.

Zacharias, M. (2017) Predicting Allosteric Changes from Conformational Ensembles. Annual Review of Biochemistry, 25, 393–394.

Zhao,K.L., et al (2021) MMpred: a distance-assisted multimodal conformation sampling for de novo protein structure prediction. Bioinformatics, 37, 4350–4356.

Zhang,G.J., et al (2022) An overview of Multi-domain Protein Structure Prediction Methods. Journal of University of Electronic Science and Technology of China, 51, 820–829.

Zhao,K.L., et al (2022) Protein structure and folding pathway prediction based on remote homologs recognition using PAthreader. Communications Biology, 6, 243.

## References

[S1] Feng,Q.Q., et al (2022) Construct a variable-length fragment library for de novo protein structure prediction. Briefings in Bioinformatics, 23, bbac086.

## References

[S2] Zhao,K.L., et al (2021) MMpred: a distance-assisted multimodal conformation sampling for de novo protein structure prediction. Bioinformatics, 37, 4350-4356.

## References

[S3] Liu,J., et al (2020) CGLFold: a contact-assisted de novo protein structure prediction using global exploration and loop perturbation sampling algorithm. Bioinformatics, 36, 2443-2450.

[S4] Liu,J., et al (2022) A de novo protein structure prediction by iterative partition sampling, topology adjustment and residue-level distance deviation optimization. Bioinformatics, 38, 99-107.

